# RoPod, a customizable toolkit for non-invasive root imaging, reveals cell type-specific dynamics of plant autophagy

**DOI:** 10.1101/2021.12.07.471480

**Authors:** Marjorie Guichard, Sanjana Holla, Daša Wernerová, Guido Grossmann, Elena A. Minina

## Abstract

Arabidopsis root is a classic model system in plant cell and molecular biology. The sensitivity of plant roots to local environmental perturbation challenges data reproducibility and incentivizes further optimization of imaging and phenotyping tools.

Here we present RoPod, an easy-to-use toolkit for low-stress live time-lapse imaging of Arabidopsis roots. RoPod comprises a dedicated protocol for plant cultivation and a customizable 3D-printed vessel with integrated microscopy-grade glass that serves simultaneously as a growth and imaging chamber. RoPod reduces impact of sample handling, preserves live samples for prolonged imaging sessions, and facilitates application of treatments during image acquisition.

We describe a protocol for RoPods fabrication and provide illustrative application pipelines for monitoring root hair growth and autophagic activity. Furthermore, we showcase how the use of RoPods advanced our understanding of plant autophagy, a major catabolic pathway and a key player in plant fitness. Specifically, we obtained fine time resolution for autophagy response to commonly used chemical modulators of the pathway and revealed previously overlooked cell type-specific changes in the autophagy response. These results will aid a deeper understanding of the physiological role of autophagy and provide valuable guidelines for choosing sampling time during end-point assays currently employed in plant autophagy research.

## Introduction

Light-based microscopy is widely used in plant cell and molecular biology. Imaging of living plant tissues is one of the most advantageous and frequently used techniques. However, some experimental layouts can be sensitive to limitations of standard protocols. For example, prolonged incubation of a live sample mounted between microscopy glass can cause mechanical stress^1^, and gradual evaporation of the mounting medium during imaging might eventually lead to drought stress. Such drawbacks typically limit the duration of assays, time-resolution, the number of biological replicates that can be processed simultaneously and are especially impactful while studying stress-related molecular pathways.

The Arabidopsis thaliana primary root is an excellent model for live imaging, due to its predictable architecture, small size, transparency, efficient uptake of chemical compounds, and low autofluorescence. In the past decade, a number of microfluidics systems have been developed to enable time-lapse imaging of live Arabidopsis roots, e.g. RootChip^2^, RootArray^3^, Plant-in-chip^4^ device and coverslip based microfluidics device for tracking root hairs^5,6^. Such devices provide a number of significant advantages, e.g. control over root growth direction and precise timing of pulsed treatments during growth and image acquisition. However, the mechanical constraints of microfluidics chambers, albeit beneficial for some experimental layouts, might be restrictive for others. Furthermore, results from Arabidopsis grown in microfluidics chambers and on Petri plates may not always be directly comparable, as plant cultivation in a microfluidics chamber under hydroponic conditions with continuous perfusion might impact local concentration of ions, exudates, and hormones.

Time-lapse microscopy on Arabidopsis roots is typically performed to explore molecular mechanisms underpinning macroscopic phenotypes observed in plate-grown seedlings. In such case, ideally, imaging and growth conditions should be identical to ensure accurate results. In this study, we focused on establishing a user-friendly and easily accessible approach for long-term live microscopy of Arabidopsis roots under conditions most similar to the standard growth of seedlings on Petri plates. For this we developed RoPods, 3D-printed customizable chambers with integrated microscopy-grade glass, and a dedicated RoPod protocol that is also applicable for commercially available chambers.

We then illustrate the benefits of RoPod’s use for studying stress-related pathways on example of plant autophagy.

Autophagy, is a major catabolic pathway conserved in all eukaryotes^7^. It is tightly linked to sensing availability of nutrients, and serves as a gateway for multiple stress responses^7,8^. Plant autophagy is a key player regulating plant fecundity, longevity, tolerance to pathogens and abiotic stress stimuli^8–10^. Autophagy has a strong catabolic capacity and thus its activity must be tightly regulated to prevent unnecessary degradation of the cellular content^11,12^. Dynamic changes in autophagic activity and its rate, also referred to as autophagic flux^13^, are principal factors defining its physiological roles. For example, Nazio et al. demonstrated oscillatory activation of autophagy during prolonged stress suggesting it as means to keep the catabolic activity under safe and physiological threshold^14^. However, little is still known about dynamics of autophagy in plants. One of the primary reasons for this possibly was the lack of appropriate tools. A large proportion of approaches currently used for studying plant autophagy relies on standardized end-point assays that provide quick and efficient measurements, but have limited time resolution^15,16^.

By implementing RoPod we were able to perform long-term live-cell imaging of fluorescent reporters for plant autophagic activity and obtain quantitative data at high temporal resolution. This approach finally revealed the complex dynamics of plant autophagic activity and demonstrated its cell type-specific kinetics.

## Results

To reduce undesired stress during long-term time-lapse experiments, we designed the RoPod toolkit in which Arabidopsis seed germination and seedling growth occurs within a microscopy-compatible device. Here we present examples of chambers designs customized for Arabidopsis growth and root imaging, verification of the toolkit’s suggested beneficial features and illustrative use of RoPod that yielded a novel insight into plant autophagy.

### RoPod toolkit concept and design examples

In RoPod setup Arabidopsis seedlings are continuously cultivated in the commonly used 0.5xMS agar plant growth medium and remain in a consistent environment throughout growth and imaging. As a result, potential environmental changes that could affect the seedlings’ development or behavior are minimized, allowing for more accurate and reliable observations during the imaging process.

To ensure accessibility of the root for imaging, Arabidopsis seeds were placed at the edge of the agar growth medium most adjacent to bottom microscopy cover slide of a chamber and the chamber was incubated under 45° angle (**Fig. 1a**). This way, the gravity vector constantly guided root growth along the cover slide.

**Figure 1.**
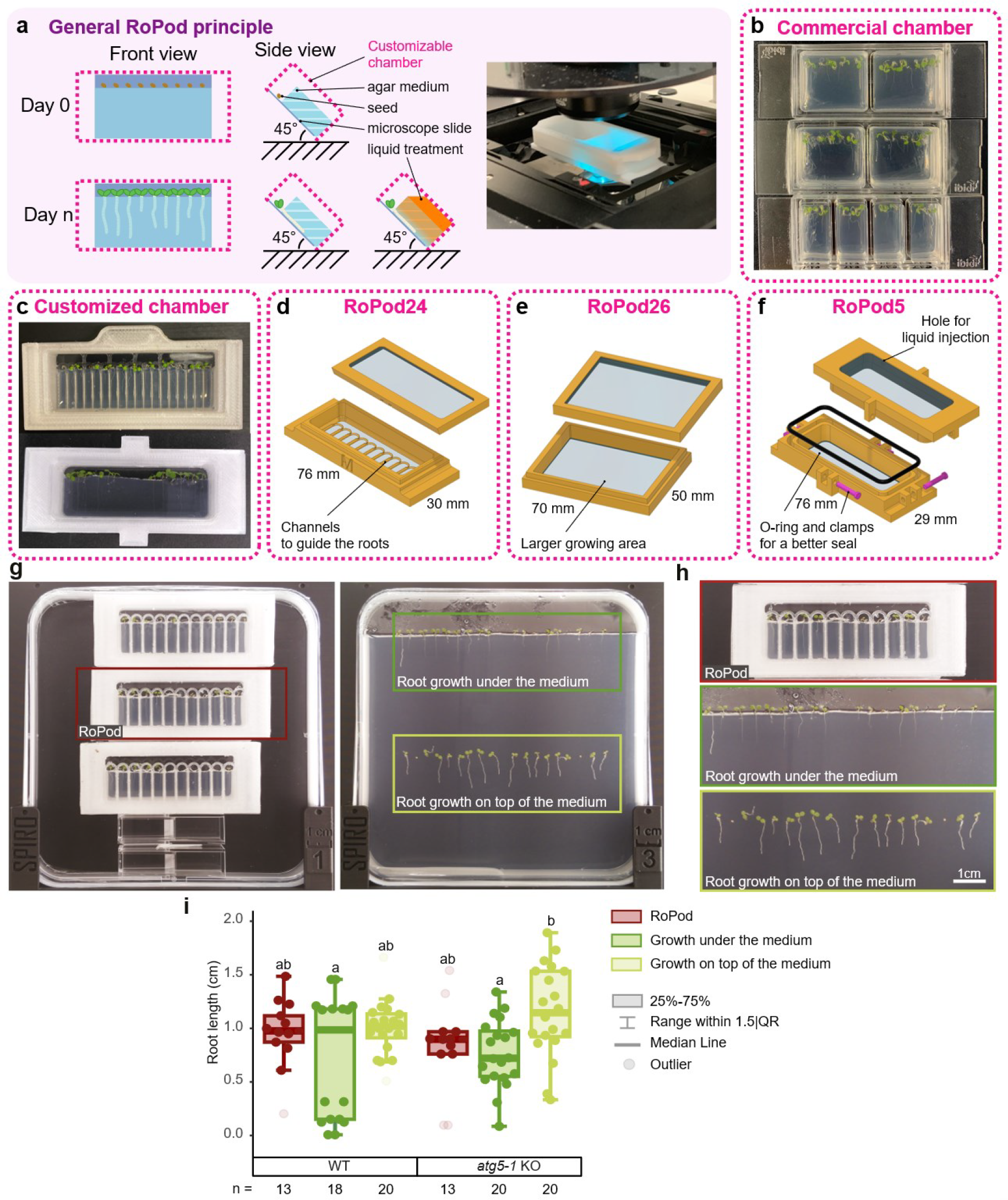
RoPod, a toolkit comprising plant growth protocol and microscopy chamber optimized for long-term time-lapse imaging of Arabidopsis roots. (**a**) the RoPod toolkit allows to grow, treat and image seedlings within the same chamber, thus keeping plants under continuous conditions that closely resemble their standard growth in the laboratory. The use of RoPods minimizes perturbations associated with remounting seedlings from growth medium onto microscopy slides. Arabidopsis seeds are plated directly into the chamber so that emerging roots grow between the agar medium and a microscope cover slide. Roots are thus immobilized in a position suitable for microscopy. The chamber itself can be either commercial (**b**), or 3D printed custom-designed (**c-f**). Custom designed chambers are optimized specifically for Arabidopsis root growth and various types of experimental layouts. For example, a RoPod chamber v24 has lanes to guide the direction of the root growth and for easy tracking of biological replicates (**d**), a larger RoPod v26 can accommodate more seedlings and enable imaging of seedlings older than 7 days (**e**), RoPod v5 is liquid-tight and allows to perform treatment of seedlings in a vertical position (**f**). (**g-i**) Growth conditions in RoPod chambers are equivalent to standard growing conditions on Petri plates, as demonstrated by comparing root growth on the top of the medium, under the medium inside the standard Petri plate and in RoPod chambers. Seeds of wild-type (WT) and autophagy-deficient *atg5-1* Arabidopsis plants expressing GFP-ATG8 were sawn on the agar medium in Petri plates or RoPods. Plates and chambers were placed on SPIRO under long day conditions and imaged for a week. (**h**) Zoomed-in insets shown in (**g**). (**i**) The chart represents data from two independent experiments. The root lengths were measured on the 7th day after seed plating. One-Way ANOVA test revealed no difference in root length of seedlings for the WT seedlings grown in RoPods and on Petri plates, significance level of 0.05, n=103.

First, we successfully tested RoPod protocol for Arabidopsis germination and growth in the commercially available microscopy chambers on the example of Ibidi and Sarstedt containers (**Fig. 1b**, see Materials and Methods). We then proceeded to further expand possible applications of the protocol by designing chambers customized specifically for Arabidopsis root growth and imaging. We aimed to design chambers that are robust, affordable, easily accessible and can be fine-tuned for various experimental layouts. Therefore, we selected to work with Fused Deposition Modeling (FDM)-based 3D printing. FDM printing enabled embedding of a microscopy cover slip directly into the plastic frame during chamber production **(Supplementary methods**), thus alleviating the need in testing phytotoxicity of different types of glues that are otherwise required to immobilize glass on the chamber. As of July 2023, the material costs for producing a single RoPod v24 comprised 0.6 Euros. Furthermore, printed-in microscopy glass remained in place even after multiple rounds of chambers’ reuse that involved washing in strong detergents followed by sterilization in 70% ethanol. As a testament of the printed design robustness, the early prototypes of RoPods have so far lasted for over two years, while being repeatedly used by multiple researchers.

The greatest benefit of the 3D printed chambers is the unlimited flexibility in the design. We conceived a number of RoPod designs to address most common requirements for various experimental layouts (**Fig. 1c-f, Supplementary methods, RoPod repository**^17^). For example, to enable easy tracking of the same biological replicates over multiple imaging sessions, we designed RoPod v24 with separate lanes that guide growth of individual roots. RoPod v24 also includes arcs to steer grow of emerging roots into the designated lanes (**Fig. 1d**). This solved the problem of some roots growing agravitropically for a short period of time after germination. To enable imaging of longer roots and a larger number of seedlings, the RoPod v26 was designed similarly to the RoPod v24 but using a bigger cover slide (**Fig. 1e**).

Commercial chambers, as well as RoPod v24 and RoPod v26 were tested to be suitable for liquid treatment when mounted on a horizontal microscope stage. For such experiments, liquid was applied on top of the agar medium and let to diffuse through the agar towards the roots at the bottom of the chamber. This layout has proven to be highly effective for short-term treatments, particularly those initiated shortly before the imaging process.

For experiments involving long-term treatments performed on a vertical microscope stage, a RoPod v5 was developed with an O-ring mounted between the lid and the chamber and clamps ensuring watertight sealing. A small aperture was included in the design to allow connection of tubing for injection of liquid treatments without opening the chamber (**Fig. 1f**).

The currently available RoPod designs have enabled us to conduct long-term time-lapse microscopy imaging spanning from several hours to several days. This extended imaging capability has facilitated the tracking and observation of various levels of biological structures, ranging from organ-level imaging to subcellular-level imaging.

### Validation of RoPod comparability with standard growth conditions

To test whether the root growth in RoPod is comparable to standard growing conditions, we assessed the root length of wild-type (WT) and the stress-sensitive autophagy-deficient *atg5-1* Arabidopsis seedlings^18^ grown and imaged on Petri plates and in RoPods (**Fig. 1g-i, Supplementary Movie S1**). For this, seeds of both genotypes were placed in square 12×12 cm Petri plates directly on the medium (to enable root growth on top of the medium) or on the edge of the medium close to the plate bottom (to enable root growth under the medium) or in RoPods following the developed protocol (**Fig. 1g-h**). The plates and chambers were then mounted on the automated imaging platform SPIRO^19^ and imaged every hour for a week. No significant effect of RoPod on root growth was observed neither for the WT nor for the mutant seedlings, confirming that root growth in RoPods is comparable to root growth on Petri plates (**Fig. 1i**).

### Validation of RoPod efficacy for minimizing undesired stress during time-lapse imaging

Next, we investigated whether the use of RoPods successfully mitigated potential stress caused by prolonged imaging sessions. For this, we focused on imaging fluorescent reporter for plant autophagy, a major catabolic pathway notoriously responsive to a plethora of stressors^20^, including common challenges associated with long-term imaging of samples mounted on microscopy glass, such as desiccation, heat and mechanical pressure^1,21–23^.We hypothesized that in the absence of any intentional stress applied to samples during imaging, the degree of autophagy induction in a sample can be used as a proxy for perturbance caused by specimen handling.

We compared induction of autophagy in Arabidopsis roots grown and imaged in RoPods versus plants grown on Petri plates and transferred on microscopy glass for imaging. For this, we used plants expressing Autophagy-related protein 8a (ATG8a) tagged with pHusion (tandem tag consisting of GFP and RFP proteins^24^) in WT or autophagy-deficient (*atg5/7*) backgrounds (**Fig. 2**). Upon activation of autophagy, ATG8 is incorporated into autophagosomes and thus changes localization from diffuse to punctate^25^. Arabidopsis seedlings grown in RoPods were scanned in the beginning and at the end of the session to ensure that no changes occurred during that time. Already after 1.5 hours of incubation, there was a significant increase in the number of ATG8a-positive puncta observed in WT plants that were transferred and mounted on microscope slides, as compared to plants grown using RoPods (**Fig. 2a and 2b**). This response was absent in autophagy-deficient mutant *atg5/atg7*, supporting the interpretation that the observed puncta were indeed autophagosomes (**Fig. 2**). This demonstrates that the use of RoPods effectively minimizes stress during time-lapse imaging.

**Figure 2.**
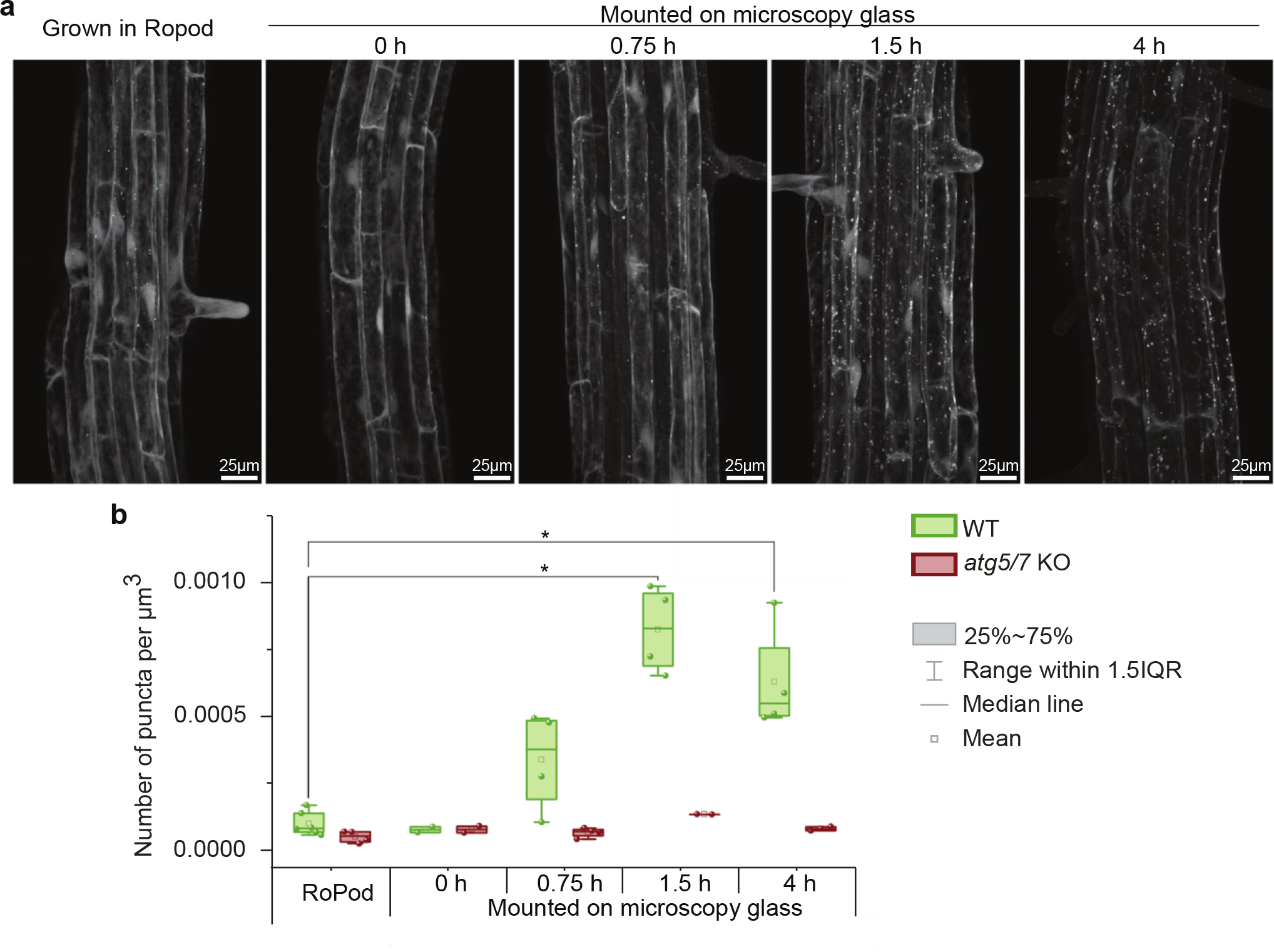
Prolonged incubation of Arabidopsis seedlings on microscopy glass upregulates autophagy in the epidermal root cells. (**a**) Arabidopsis seedlings expressing fluorescent marker for autophagy (pHusion-ATG8) in the wild-type (WT) or autophagy-deficient (*atg5/7* KO) backgrounds were grown on standard Petri plates or in the RoPod v23. 5 days-old seedlings grown on a Petri plate were mounted between a standard microscopy-grade sample slide and a cover slip in a liquid 0.5xMS medium and incubated on the bench for the designated amount of time prior to imaging using confocal microscope. Seedlings of the same genotypes grown in the RoPod were imaged using the same settings (left panel). (**b**) Quantification of pHusion-positive puncta in the root cells illustrated in (**a**) reveals gradual upregulation of autophagic activity in the roots mounted on microscopy glass. The chart comprises representative data from one out of three individual experiments. Two-tailed t-test with unequal variants, n= 35; *, p-value < 0.05.

### Assessment of chemical compound diffusion in RoPod

To test if the established RoPod protocol allows for efficient drug treatments of Arabidopsis roots, we assessed diffusion of chemical compounds in the medium inside RoPod chambers. We monitored the diffusion rate of the fluorescent dye fluorescein within horizontally positioned RoPods containing seedlings (**Fig. 3a-b**) or in RoPods containing only growth medium (**Fig. 3c**). In the chambers containing medium only, we quantified increase in fluorescence intensity at the bottom cover slip, while in the chambers containing seedlings, we measured the fluorescence accumulation at root hair tips. Quantification revealed that the dye solution added on the surface of the growth medium diffuses to the bottom coverslip and reaches the plant tissue within 15-25 min (**Fig. 3b**).

**Figure 3.**
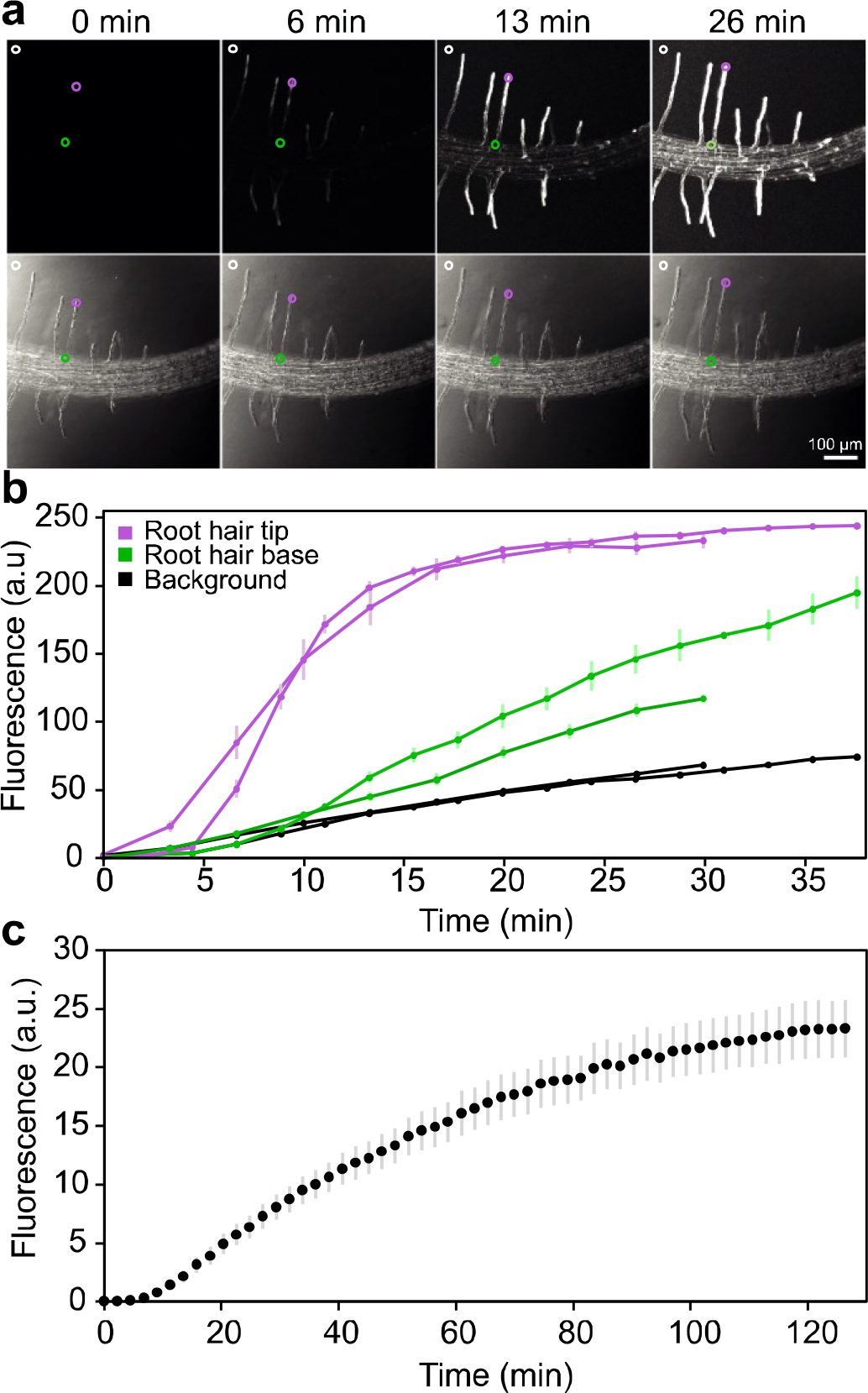
Diffusion of chemical compounds in the RoPod. (**a**) Representative time lapse images of roots treated with fluorescein in the RoPod. Arabidopsis seedlings were grown in RoPods using the described protocol. Liquid MS medium containing 2 μg ml^-1^ of fluorescein dye was added to the chambers immediately prior to the start of imaging. Maximum intensity projections of the fluorescent signal are shown on the top row, corresponding transmitted light images are shown in the bottom row. The circles indicate three types of regions of interest (ROI) analyzed for each root hair to produce the data presented in(**b**). Magenta circle, root hair tip; green circle, root hair base; white circle, background. (**b**) Quantification of the data illustrated in (**a**). Diffusion rate of fluorescein in the RoPod chamber demonstrates that chemical compounds reach root hair cells within 15 minutes of application. Data for two independent experiments is plotted for each type of analyzed ROI (**c**) Dynamics of fluorescence accumulation at the bottom coverslip of RoPod. In this experiment liquid MS medium containing 140 μg ml^-1^ fluorescein was added to RoPods containing only growth medium. The fluorescence was recorded in four separate chambers, with 2 to 3 fields of view chosen at random positions in each RoPod.

A full equilibration of the concentrations between top and bottom of chamber occurs in approximately 120 min (**Fig. 2c**). Such treatment rates are well-suited for studying relatively slow dynamic responses (tens of minutes to hours), such as plant autophagy.

### Applicability of RoPod for long term treatment and imaging illustrated by monitoring root hair growth response to changing sucrose concentrations

Next, we examined the suitability of RoPods for extended imaging sessions and the application of chemical compounds treatments during the acquisition process. For this, we used root hair growth rate as a read-out. Root hairs sustain a plethora of functions including nutrients uptake, plant anchoring in soil and symbiotic interactions (see Rongsawat *et al*., 2021 for review^26^). Their growth rate is known to be affected by mechanical stress, fluctuations in humidity and sucrose addition to the medium^27–29^. Under standard conditions, Arabidopsis root hairs elongate at the rate of approximatively 1 μm min^-1 30^ and a typical root hair reaches approximatively 100 – 400 μm in length depending on the growth conditions. Hence, the entire process of root hair formation from initiation to termination spans several hours.

We designed an experimental setup that involved addition of medium supplemented with 1% sucrose during the course of imaging and analyzed hair growth under unperturbed conditions and upon treatment. By implementing RoPod on a vertical microscope, we recorded root hair growth within a time range of 10 h, with a time resolution of 8 min, for 22 roots and 624 root hairs in total under control or sucrose-rich conditions (**Fig. 4a, Supplementary Movie S2**). To enable analysis of large imaging data, we developed a semi-automated image analysis pipeline involving designated *Fiji* macros (see Materials and Methods for details and https://github.com/AlyonaMinina/RoPod). This pipeline facilitated efficient analysis of the 35244 acquired images. The treatment with sucrose increased the final hair length around 2.3 fold (without sucrose: 102.7 +/- 5.3 μm; with sucrose: 234.7 +/- 8.0 μm (mean +/- SE)), the growth rate increased around 1.5 fold (without sucrose: 0.75 +/- 0.02 μm min^-1^; with sucrose: 1.14 +/- 0.02 μm min^-1^ (mean +/- SE)), and the growth duration extended 1.6 fold (without sucrose: 2.2 +/- 0.1 h; with sucrose: 3.5 +/- 0.01 h (mean +/- SE)) (**Fig. 4b-f**). Taken together, these results demonstrate the applicability of RoPods to provide suitable growth conditions for long-term imaging at cellular resolution, as well as the possibility to apply treatments in the course of an experiment. We also provide a complete pipeline for root hair growth analysis under physiologically relevant conditions.

**Figure 4.**
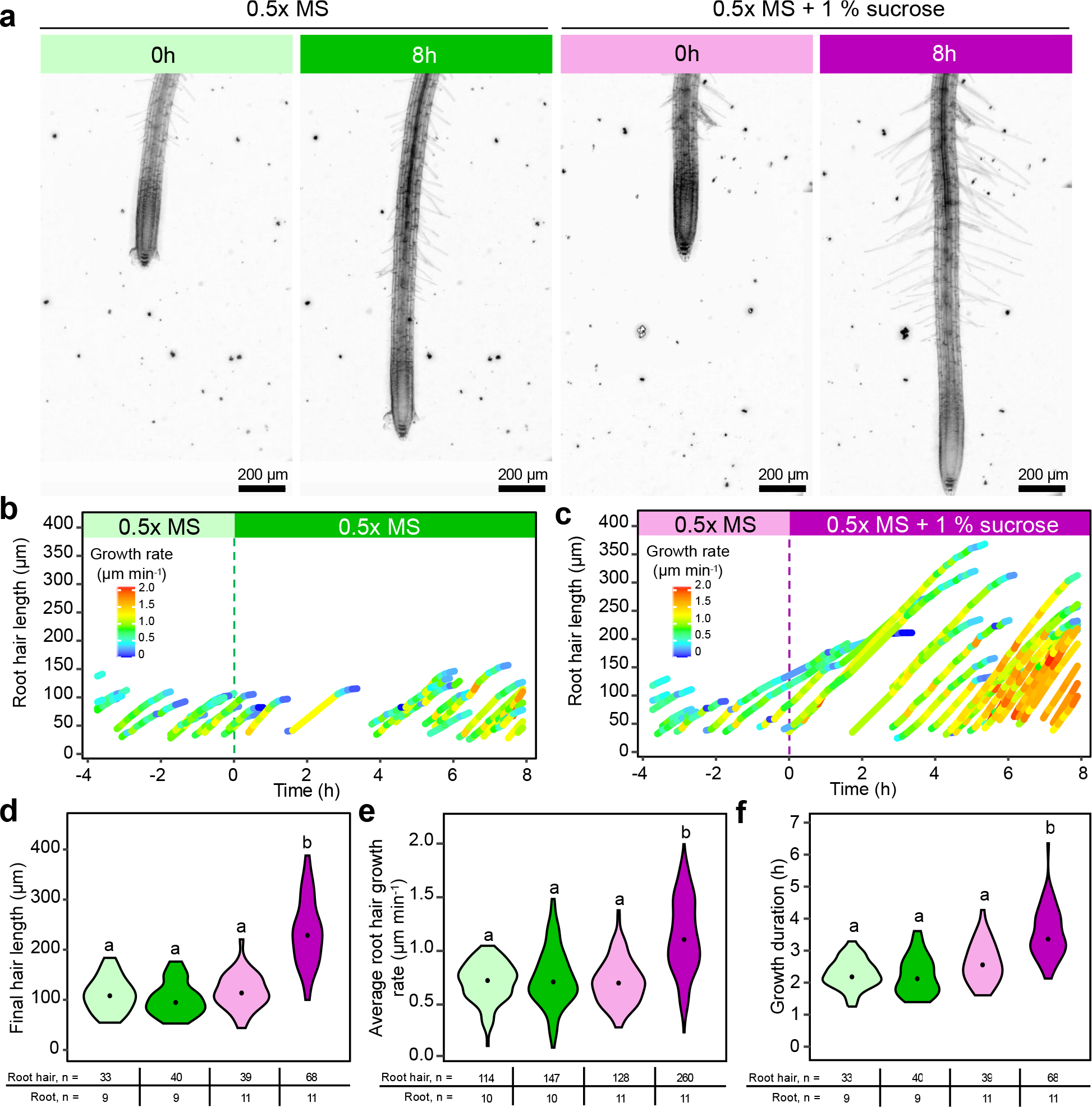
Monitoring sucrose effect on root hair elongation illustrates applicability of RoPod for long-term time-lapse imaging combined with treatments. Arabidopsis Col-0 WT seedlings were grown for one week in a RoPod5 using the described protocol. The growth of root hairs was firstly recorded for 4 h under control conditions. After that, the chambers were flooded with liquid 0.5x MS (control treatment) or liquid 0.5x MS supplemented with 1 % of sucrose (sucrose treatment). (**a**) Representative images showing roots at 0 h and 8 h of treatment. Each panel is a z-projection covering 240 μm. (**b-c**) Result of root hair tip tracking for one representative root for control (**b**) and for sucrose (**c**) treatments. The final root hair length (**d**), the root hair growth rate (**e**) and the growth duration (**f**) were calculated as described in the **supplementary Methods**. (**d-f**) The sample size used for each measurement is shown in the tables below the charts. A Dunn’s multiple comparison test, different letters designate significantly different groups, P < 0.01. The dot in the violon plots is the median of the distribution.

### Implementation of RoPod reveals cell type-specific dynamics of plant autophagy

We then proceeded to further explore advantages of RoPods implementation for studying stress-related pathways using plant autophagy as a model.

Imaging of autophagy-related structures, autophagosomes, is often used as a readout of autophagic pathway activity^25^. The above-mentioned autophagy’s sensitivity to a broad range of stress triggers makes it a prime example where minimally perturbing imaging conditions are critical to provide accurate biologically relevant results.

Furthermore, currently established protocols for measuring plant autophagic activity are end-point assays and do not provide sufficient time resolution to study autophagic flux fluctuations.

To test if the kinetics of autophagosomes accumulation could be studied in RoPod-grown plants, we performed time-lapse tracking of pHusion-tagged ATG8a in WT and autophagy-deficient *atg5/atg7 Arabidopsis thaliana* lines. Seedlings were grown in the RoPods for 5-7 days using the protocol described in the Materials and Methods. Upon induction of autophagy ATG8 is incorporated into double-membrane vesicles, autophagosomes, which sequester cargo from cytoplasm and deliver it to the lytic vacuole for degradation^7]^. Therefore, the number of autophagosomes visualized as ATG8-positive puncta is considered representative of autophagic activity^25^. The basal level of autophagic activity in cells not subjected to stress was measured in seedlings treated with 500 nM concanamycin A (ConA), a chemical compound that deactivates lytic vacuolar enzymes thereby preserving autophagosomes from degradation after they have been delivered to the vacuole^15^. To artificially induce autophagic activity, we applied treatment with 500 nM AZD 8055 (AZD), a commonly used inhibitor of TORC1, a kinase complex which under normal conditions dampens autophagic activity^15,31^. The compound stocks were diluted in 0.5x MS liquid medium and added to the seedlings in RoPod chambers immediately prior to scanning. Approximately 100 μm deep z-stacks were acquired for each biological replicate every 15 min. Maximum intensity projections of the optical slices for each biological replicate at each time point were used for further image analysis. Representative images of pHusion-ATG8a localization at 8 h (for ConA) and 3 h (for AZD) of the treatment are shown (**Fig. 5a)**. The complete time series are available in **Supplementary Movie S3** where the viability of imaged tissue was confirmed by the cell elongation and root hair formation during imaging. During the total 13h duration of recording we did not observe any increase in the number of pHusion-ATG8-positive puncta in mock-treated seedlings, confirming that imaging *per se* was not inducing autophagic activity (**Fig. 5b**).

**Figure 5.**
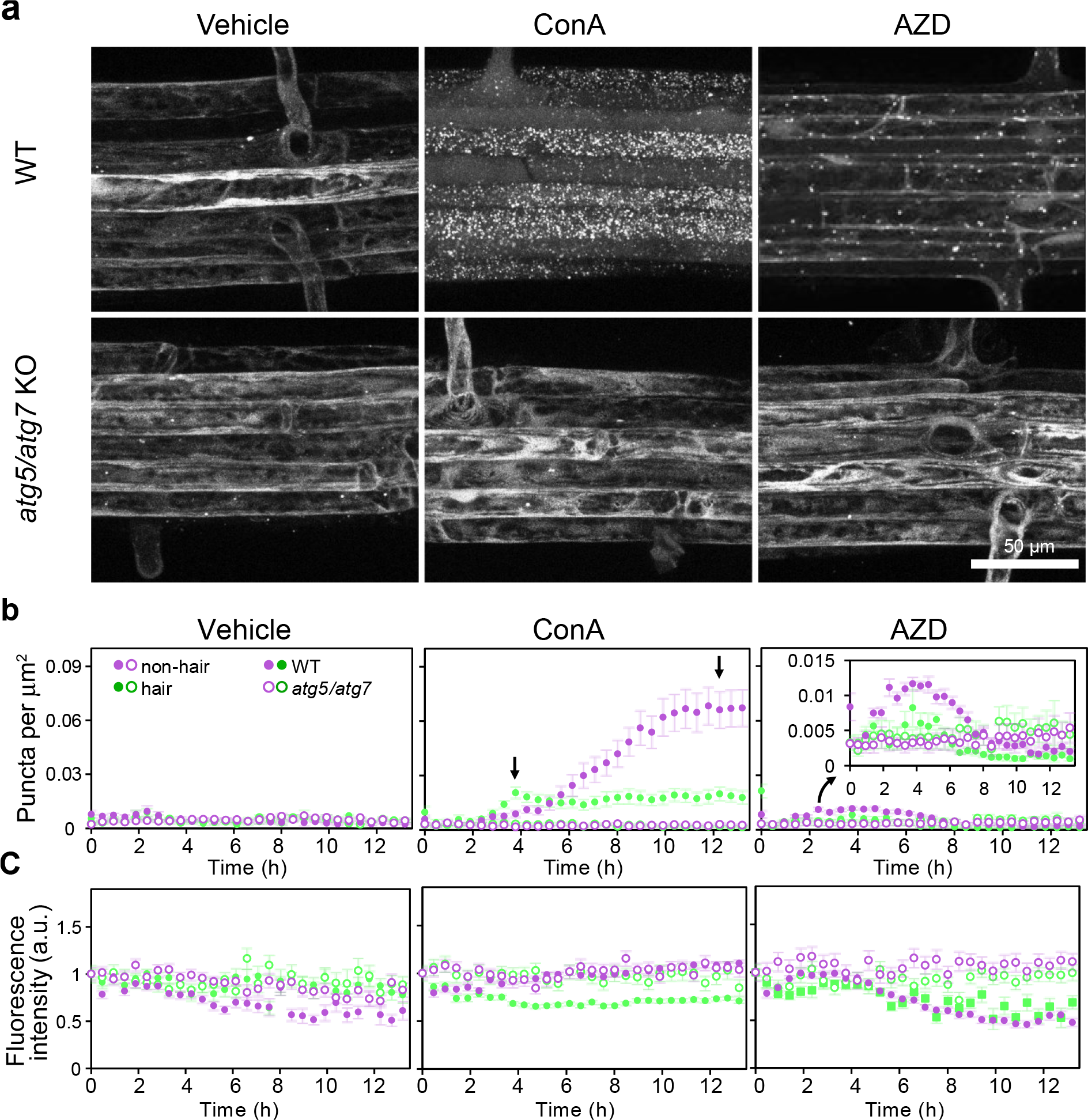
Time-resolved detection of plant autophagic activity reveals it cell type-specific dynamics. Arabidopsis seedlings expressing autophagosomal marker pHusion-ATG8a in the wild-type (WT) or autophagy-deficient (*atg5/7* KO) backgrounds were grown in the RoPods and treated with 0.5 μM AZD8055 (AZD) to induce autophagy, or with 0.5 μm Concanamycin A (ConA) to visualize basal autophagy, 0.1% DMSO (Vehicle) was used as a negative control for the treatments. Only GFP channel is shown.(**a**) Example time frames illustrating punctate localization of the autophagosomal maker pHusion-ATG8 in epidermal root cells upon drug treatment. Scale bar, 50 μm. (**b**) Quantification of the time-lapse data illustrated in (**a**) reveals early detectable but moderate basal autophagic activity in hair cells and a delayed but stronger activity in non-hair cells. The arrows on the ConA chart point out the peaks for cell-type specific autophagic activity. Additionally, statistical analysis of puncta accumulation at 0h, 4h and 12h in the hair and non-hair root cells is shown in the **Fig. S1**. Treatment with AZD reveals stronger response in the non-hair cells. The inset on the AZD chart presents a zoomed view of the data indicated by the arrow. The dots indicate mean ± SE calculated with the number of puncta counted in each cell type. (**c**) Quantification of the GFP intensity on the data illustrated in (**a**) demonstrates eventual significant decrease of signal in AZD-treated WT roots. The dots indicate mean ± SE calculated with individual measurement of each punctum. The mean intensity has been corrected with the background signal and normalized to the first time point of the record. The fluorescence intensity was measured for each vesicle counted, for a total population per cell file of 8 to 2339 vesicles, depending on the genotype, the treatment, the cell type, and the time of the record considered. For (**b**) and (**c**) each presented time point corresponds to an interval of time of up to 15 min.

Time-lapse imaging using RoPods confirmed previously observed differences in autophagic activity in hair and non-hair epidermal root cells of wild-type plants^31^. Namely, ConA treatment shorter than 5 h resulted in the higher number of fluorescent puncta accumulating in the vacuoles of the hair root cells (**Fig. 5b, Supplementary Figure S1**). Surprisingly, at later time points, the number of puncta in hair cells plateaued, while in non-hair cells it increased more than threefold (**Fig. 5b, arrows on the ConA chart, Supplementary Movie S3 and Figure S1**). We confirmed that observed changes in localization of pHusion-ATG8a were autophagy-specific by demonstrating no induced formation of pHusion-ATG8a puncta and no discernable differences between hair and non-hair cells in autophagy-deficient *atg5/7* mutant seedlings (**Fig. 5**). We then tested if the observed differences in puncta densities might be caused by the differences in the cell volume of the slowly elongating non-hair and rapidly expanding hair cells. For this, the number of puncta was quantified at 12h of ConA in the full volume of both hair and non-hair cells, resulting in an approximately 12-fold higher puncta density and 8-fold higher puncta count per cell in non-hair cells compared to hair cells (**Supplementary Figure S2**).

Consistently with the results obtained on ConA treated roots, treatment with AZD also revealed a stronger upregulation of autophagic activity in the non-hair cells, although at earlier stages (2-4h) of treatment (**Fig. 5b**). This indicates that non-hair cells might possess higher capacity not only for basal autophagy but also for stress-induced autophagic activity, albeit they show a delayed response when compared to the hair cells.

It did not escape our attention that the number of puncta in the WT and autophagy-deficient background became comparable after approximatively 6h of AZD treatment (**Fig. 5b)**. The occurrence of the few puncta in the autophagy-deficient background could be explained by association of ATG8 with protein aggregates, which could not be cleared in the absence of autophagy^32^. The decrease in the number of puncta in the WT can be explained by autophagy-based degradation of the ATG8 reporter, which would eventually drive signal intensity below detection limit. Indeed, after 4h of treatment, the mean intensity of the fluorescent marker becomes significantly lower in the cells of wild-type plants compared to the *atg5/7* knockout cells (**Fig. 5c**), rendering the data from these two types of samples not comparable.

Taken together, we found that plant autophagic activity fluctuates in a cell type-dependent manner, namely that both basal and induced autophagic activities are present at different levels in hair and non-hair root cells. These results highlight the dynamic nature of autophagy in response to specific cellular contexts and emphasize the importance of considering cell type-specific dynamics when studying plant autophagy.

## Discussion

The RoPod toolkit is by no means a replacement for currently existing solutions, but rather an addition offering important advantages for a broad range of experimental layouts. RoPod is designed to minimize physical constraints experienced by the root during growth and mitigate potential stress-responses triggered by transfer from growth conditions to a microscope stage, thus facilitating long term microscopic imaging. An important advantage of the 3D printable chambers is the ease of chamber production and customization. We designed RoPods to be compatible with commonly used microscopy equipment and publish model files to enable further customizations. For the posterity reasons, the growing collection of RoPod models is made available under open-source license in a GitHub repository^17^, which will allow to add updates even after publication of this study.

By implementing the RoPod toolkit, we provide, for the first time, data on the dynamics of basal and induced plant autophagy detected in root epidermal cells and show cell type-specific differences in their kinetics.

We implemented the commonly used ConA^25^ to visualize basal autophagic activity, which helps to catabolize superfluous cellular content and thus plays an important role in maintenance of cellular fitness and recycling of internal nutrients sources^20,33^. We observed an initially stronger basal autophagy in hair cells which later became much more intense in the neighboring non-hair cells. One could speculate, that this phenotype could be caused by different sensitivity of hair and nonhair cells to ConA. However, to the best of our knowledge, there is no empirical evidence indicating such cell type-specific ConA effect. Therefore, the cell type-specific autophagic activity pattern is most likely to stem from differences in autophagic pathway regulation in hair and non-hair root cells. Future studies are necessary to reveal molecular mechanisms underpinning this interesting phenotype. Higher basal autophagic activity might be linked to the more efficient PIN2-dependent trafficking of endogenous auxin in the non-hair root cells^34^. This plant hormone has been shown to crosstalk with autophagy^35,36^. Possibly, basal autophagic activity which is an integral part of endomembrane trafficking is dampened in the rapidly growing hair cells to conserve endomembrane trafficking capacity required for hair elongation. Future studies comparing basal autophagy at successive stages of root hair development will help to assess whether the same pattern of response is seen in fully established hair and non-hair cells. Furthermore, root hair cells that are evolved to take up nutrients from soil^37^ might also have evolved a different link between autophagy and nutrients sensing, resulting in decreasing basal autophagic activity associated with root hair development.

Application of AZD creates an artificial signal, mimicking scarcity in amino acid content^38,39^. Autophagy induced upon such treatment serves to degrade and upcycle nonessential cellular content to provide required building blocks^39^. Remarkably, non-hair cells demonstrated a higher capacity for AZD-induced autophagic activity, when compared to the neighboring hair cells of the same root. This result corroborates the hypothesis of cell-type specific roles of autophagy in the multicellular organism. Potentially, starvation-induced autophagy might be dampened in the hair cells that would still need to grow seeking for new sources of nutrients, while high autophagic activity in the neighboring non-hair cells could serve as a local source supporting such growth.

A combination of AZD and ConA treatments is frequently used in the end-point assays to assess plant autophagic activity^25]^. Typically, interpretation of such treatments is that ConA blocks the final step of the AZD-induced autophagic activity^25^. However, our results indicate that at early stages (2 - 6h) of such treatment applied to Arabidopsis roots, one might be visualizing a strong basal autophagy of root hair-cells combined with strong AZD-induced autophagy in non-hair cells.

Lastly, we demonstrate that RoPod toolkit allows to estimate the optimal time range for quantification of autophagy reporters under various conditions. For example, time-lapse detection of pHusion-ATG8a in plant root cells upon AZD-treatment illustrated the gradual decay of the reporter intensity that eventually reached level below quantitative threshold (**Fig. 5c**). Similar observation of decreased signal intensity upon prolonged AZD treatment was previously interpreted as a reduction of autophagic activity^40^. However, it cannot be excluded that autophagy-driven degradation of the fluorescent reporter, in combination with decreased efficacy of its *de novo* translation due to TORC1 deactivation, reduce detectability of the reporter eventually rendering it unsuitable for measurements. The time-resolved data on autophagic activity obtained in this study can be further used as a guideline for accurate interpretation of the commonly used end-point assays for plant autophagic activity.

In summary, our observations provide a foundation for conducting more comprehensive investigations into the cell type-specific roles of autophagy in plant physiology. Additionally, we provide a tool that will contribute to refining the interpretation of commonly used end-point assays in plant autophagy research and further broaden the range of feasible topics, facilitating studies on kinetics of plant autophagosome formation and trafficking, link between autophagy and plant circadian cycle, crosstalk between autophagy and plant cell differentiation.

Furthermore, we demonstrate a broader applicability of RoPod for minimally perturbing root imaging across the macro- and microscale. On the example of root hair growth, we showcase the suitability of RoPods not only for studying the dynamics of stress-response related pathways, but also for growth assays.

## Materials and methods

### Genetic constructs

pMDC pHusion 1 Gateway vector was generated by amplifying pHusion tag from the p16:SYP61-pHusion construct^27^ using primers TTGGTACCATGGCCTCCTCCGAGGACG and TTGGCGCGCCCCCCTTGTACAGCTCGTCCATG and introducing amplicon into pMDC32 vector ^41^ via KpnI/AscI restriction digestion sites.

At ATG8a CDS Gateway entry clone ^31^ was then recombined with the pMDC pHusion 1 Gateway vector to obtain the construct driving expression of pHusion-AtATG8a under control of double 35S promoter.

### Plant material

Autophagy deficient *atg5/atg7* double knockout line was established by crossing previously published *atg5-1*^18^ and *atg7-2*^42^ mutants.

Col-0 wild-type and *atg5/atg7* plants were transformed using GV3101 strain of *Agrobacterium tumefaciens* carrying pMDC pHusion1 AtATG8a construct as described in ^31^.

Col-0 WT and *atg5-1* plants expressing GFP-ATG8a line was described previously^10,18^.

Col-0 WT plants co-expressing vacuolar marker spL-RFP and the autophagosomal marker GFP-ATG8a were obtained by crossing the corresponding marker lines previously published in Hunter et al 2007^43^ and Thomson et al 2005^18^.

### RoPod chamber printing and plant growth protocol overview

In this study we firstly tested five types of microscopy coverslip bottom chambers from three different providers: Ibidi chambered coverslips (cat # 80421 and 80286 #1.5 polymer coverslip, hydrophobic, sterilized, Ibidi, Germany), 1-well II Chamber Slide™ System (Nunc™ Lab-Tek™, cat # C6307, Sigma-Aldrich), x-well cell culture chamber, 1-well, on coverglass (cat # 94.6190.102, Sarstedt). Plants in these chambers were grown using the same protocol as was later implemented for the custom designed chambers (**Supplementary Methods)**.

The different functional designs of RoPod chambers, as well as an illustrated step-by-step protocol for their 3D printing are presented in **Supplementary File S1 and Supplementary Methods**. Updates of the designs and of the printing protocols depending on the model of 3D printer used are available in the dedicated GitHub repository: https://github.com/AlyonaMinina/RoPod.Hardware.

Printing of the chambers was tested using Prusa MK2.5S, MK3S, MK3S+, MINI+ and Ultimaker S5 3D printers. We recommend using clear PETG filaments provided by Prusament, Sunlu or Prima Select.

The recommended protocol to grow seedlings in RoPods is presented in the **Supplementary Methods**. To reuse the chambers, the growth medium was gently removed using a Q-tip, the chambers were washed with cold water and dish soap, or in 1% SDS, abundantly rinsed with tap water followed by rinsing with MilliQ water, air dried and sterilized for 1 hour under UV light. Alternatively, chambers were sterilized by incubating for 40 minutes in 70% ethanol, followed by overnight drying in the sterile bench.

### Root length measurement for seedlings grown in RoPods and on standard Petri plates

To compare the root growth phenotypes for seedlings grown in RoPods and in standard Petri plates, 4 ml of growth medium (0.5x MS (*Duchefa*, ref. M0222); MES 10 mM (*Duchefa*, ref. M1503); 1 % sucrose; pH 5.8; 0.8 % Plant agar (*Duchefa*, ref. P1001) were pipetted into each sterile RoPod v24 and 40 ml were poured into 12cmx12cm plates. A strip of the medium was removed from the RoPods or plates using a sterile scalpel. Seeds of GFP-ATG8 expressing lines were sterilized using chlorin method (incubated in 10% Chlorin; 0.001% Tween-20 for 20 minutes then washed 6 times with sterile water) and then plated into RoPod v24, or in standard plate, either on the edge between the agar and plastic plate bottom or on top of the agar. Three RoPod v24 were placed into a 12 cm square empty plates. Both standard agar plate and plate loaded with RoPod24 were mounted on the automated imaging platform SPIRO^19^ and imaged every hour for seven days under 16h illumination (Day imaging settings: ISO 50, shutter speed 65; Night imaging settings: ISO 250 Shutter speed 10). The root length was measured manually on obtained images using *Fiji* software (ImageJ 1.53t^41^).

### Comparison of autophagic activity in roots mounted on glass or in RoPods

To assess the impact of glass-based sample mounting on the autophagic activity, RoPod v24 and standard agar plates were prepared similarly to the experiment for the root length measurement. After 6 days, seedlings growing on top of the agar medium were transferred in an empty glass bottom chamber, on a drop of 0.5x MS and then gently covered with a cover slide. To prevent fast evaporation of the liquid medium, the glass bottom chamber was covered with a lid. Both RoPod v24 with seedlings and glass bottom chamber with transferred seedlings were imaged at different end points using CLSM 800 (Carl Zeiss), Objective 40x/1.2W, excitation light 488 nm, and emission 505-560 nm. The number of puncta was quantified in *Fiji* using the workflow presented in **Supplementary File S2**.

### Fluorescein diffusion assay

To assess diffusion of chemical compounds within RoPod chambers containing plants, Arabidopsis Col-0 seeds were surface sterilized in opened 2 ml tubes for 2 h in an air-tight box containing chlorine gas, produced by the mix of 50 ml of sodium hypochlorite 10% (*Roth*, ref. 9062.3) and 2 ml of 37% hydrochloric acid (*Merck*, ref. 1.00317.1000). Seeds were then aerated for 10 min in a sterile bench to remove the left-over chlorine gas. Seeds were then transferred, with a sterile toothpick, in 2-wells *Ibidi* chambered coverslips (cat # 80286 #1.5 polymer coverslip, hydrophobic, sterilized, Ibidi, Germany) filled with 1 mL of growth medium in each well (0.5x MS (*Serva*, ref. 47515); MES 5 mM (*Sigma*, ref. M8250); 1% sucrose; pH 5.7 adjusted with KOH; 0.8 % Plant agar (*Duchefa*, ref. M1002). As the height of the Ibidi chamber is too small to accommodate nine days old seedlings, the seeds were put on the surface of the agar to be first cultivated for eight days horizontally at 21 °C, 16 h illumination, 100 μmol m^-2^ s^-1^, until the root penetrate the agar and reached the glass bottom. Then, the RoPods were incubated at 45° angle for one day before image acquisition. At the start of the recording, each well containing plants was filled with 1 ml of fluorescein at 4 μg ml^-1^ diluted in a liquid 0.5x MS (final concentration after diffusion: 2 μg ml^-1^). Imaging was performed using *Leica* DMI8 confocal microscope and HC PL APO CS2 20x objective (excitation, 488 nm, emission, 498 nm - 550 nm, pinhole size of 1 Airy Unit) (Leica Microsystems, Germany), pixel dimensions 0.48 μm px^-1^, image size 512 x 512 px, 8-bit. The root hairs that were growing at the beginning of the recording were selected for analysis. To obtain a complete view of the root hairs, a z-stack was performed to cover 92 μm height with each section separated with 10 μm. The images were then processed on *Fiji* to obtain maximum intensity projection. A circular **r**egion **o**f **i**nterest (ROI) with a 12 μm diameter was used to measure the mean grey value on the selected root hair tips, on the base of the corresponding root hair tip and in the upper left corner of the image for the mean grey value of background. The experiment was performed in two individual boxes containing 2 to 3 roots each, 2 to 8 root hairs analyzed per root.

For the fluorescein diffusion assay in the solid medium and without plants, 2-wells *Ibidi* chambers filled with 1 ml of the same 0.5x MS agar in each well. 1 ml of liquid growth medium supplemented with 140 μg ml^-1^ of fluorescein solution was added at the start of the recording (final concentration after diffusion: 70 μg ml^-1^). The same microscope settings as for the previous diffusion assay were used (pixel dimensions 1.14 μm px^-1^, image size 512 x 512 px, 8-bit). The fluorescence was recorded in 4 RoPods, with 2 to 3 fields of view chosen at random positions in each box. *Fiji* software was used for image analysis. An orthogonal projection was carried out in the middle of each acquisition area with the *Orthogonal view* tool. To define the bottom of the RoPod, the orthogonal view of the image acquired 2 h after the start of treatment was used. A rectangular ROI with the width of the orthogonal projection image and 10 μm in height was drawn at the bottom of the RoPod chamber. This ROI was reused to measure the mean grey value on the orthogonal view for each time point recorded at the same location.

### Autophagic activity assay

Growth medium (0.5x MS (*Duchefa*, ref. M0222); MES 10 mM (*Duchefa*, ref. M1503); 1 % sucrose; pH 5.8; 0.8 % Plant agar (*Duchefa*, ref. P1001)) was autoclaved for 20 min at 120 °C, cooled to approximatively 60 °C and pipetted under sterile conditions into either 4–wells *Ibidi* coverslips (cat # 80421 #1.5 polymer coverslip, hydrophobic, sterilized, Ibidi, Germany) for the experiment presented in **Figure 5** or into RoPod v24 for the experiment presented in **Supplementary figure S2**. 1ml of medium was used per well of *Ibidi* chamber and 4 ml were used for the RoPod v24.

*Arabidopsis thaliana* seeds were surface sterilized in 70 % EtOH, 0.05 % TritonX-100 for 20 min, washed three times with 96 % EtOH, air dried in a sterile bench and transferred into the chambers using a sterile toothpick. The chambers were then placed under long day growth conditions (16 h, 150 μmol m^-2^ s^-1^ light at 22 °C, 8 h dark at 20 °C). Typically, for the first four days the chambers were kept horizontally allowing the roots to reach the bottom cover slip. After this, the chambers were placed vertically to guide the root growth along the cover slip.

Chemical compounds were diluted in 0.5x MS liquid medium (the same composition as the growth medium, but lacking Plant agar) to a final concentration of 1 μM AZD8055 (*Sigma*, ref. ADV465749178, CAS number 1009298-09-2), 1 μM ConA (*Sigma*, ref. C9705, CAS number 80890-47-7) or 0.1 % DMSO for the vehicle control. The solutions were pipetted into the wells of the chambers mounted on the confocal stage in 1:1 ratio (1 volume of the medium in the well: 1 volume of the drug solution. Drug final concentration: 500 nM) right before the start of imaging.

Imaging was performed using CLSM Leica SP5 or SP8 (Leica Microsystems, Germany), 63x water immersion objective, NA=1.2, excitation light 488 nm, emission range of 498 nm - 541 nm and system-optimized pinhole size or and Zeiss CLSM 800 Objective 40x/1.2W and similar scanning settings. For this study, pHusion tag was used solely as a fluorescent reporter and ot as a pH indicator. Four to six biological replicates were imaged for wild-type and *atg* KO backgrounds in each experiment. The experiment was performed four times in two independent laboratories. Images were acquired at the beginning of the root differentiation zone, collecting approximatively 100 μm deep z-stack starting at the surface of a root (pixel dimensions 0.29 μm px^-1^, image size 848 x 848 px, 8-bit). Each replicate was imaged every 10 to 28 min for a period of 12 to 18 h.

Images in **Figure 5** were analyzed with *Fiji* and Ilastik^44^. The detailed protocol and the macro used for the analysis are presented in the **Supplementary File S3 and Supplementary Methods**. The number of vesicles was counted for each cell file type using data obtained from two independent experiments. 1 to 5 roots and 2 to 15 cell files per type were analyzed in total. The fluorescence intensity was measured for each vesicle counted, for a total population per cell file of 8 to 2339 vesicles, depending on the genotype, the treatment, the cell file type and the time of the record considered. The mean intensity has been corrected with the background signal and normalized to the first time point of the record. As the image acquisition rate varied between experiments due to different number of biological replicates, each presented time point actually corresponds to an interval of up to 15 min.

For quantification of autophagic bodies in the complete cell volume, seeds of the transgenic line co-expressing spL-RFP and GFP-ATG8a were sterilized using the above described Chlorin method and plated either on Petri plates or in the RoPod v24. 5 days old seedlings were treated with Concanamycin A for 12 h as described above and imaged using Zeiss CLSM 800 Objective 40x/1.2W to obtain tiled, z-stack encompassing complete root hair cells. Puncta quantification was performed using *Fiji* and dedicated macro as described in the **Supplementary File S4**.

### Plant growth and image analysis of sucrose treatment assay

*Arabidopsis thaliana* Col-0 wild-type seeds were surface sterilized with the chlorine gas method (see the Fluorescein section of Material and methods). The seedlings were grown in sterile RoPod v5 chambers containing 3 ml of solid growth medium (0.5x MS (*Serva*, ref. 47515); MES 5 mM (*Sigma*, ref. M8250); pH 5.7 adjusted with KOH; 0.8 % Plant agar (*Duchefa*, ref. M1002)), as described in **Supplementary Methods**. The injection hole of the RoPod v5 was closed with micropore tape. The chambers were placed into a 12×12cm dish, and positioned with an angle of 45° into a plant growth cabinet with side illumination, 16h light, 20°C for six days. The recording was made using a 90°-flipped microscope for vertical specimen mounting (*Zeiss* AxioVert 200M, *ITK* MMS-100 linear stage and Hydra controller, *Hamatsu* OrcaC11440-36U camera, *Zeiss* 10x PlanNeoFluar objective). LED bulb compatible for plant culture (LED power 150W, Chip SMD2835, 215 pcs Warm, 40pcs Red, 5pcs blue) was set to illumination intensity of 100 μmol m^-2^ s^-1^ measured at the proximity of the RoPod). Tubing connected to a syringe was inserted into the injection hole and taped on the microscope prior to imaging start in order to limit the vibration during injection. Two RoPod v5 were mounted at the same time on the microscope. A basal line was recorded for 4 h for both control and treated samples. Then, 8 ml of 0.5x MS liquid supplemented with 1 % sucrose were added in one of the RoPods (final concentration: 0.8 %) and control specimen were treated with 8 ml of 0.5x MS without sucrose. Root hair growth was recorded for an additional 8 h. For each root, two consecutive XY fields of view were recorded to obtain images of the root hairs at different stages of development throughout the recording time (pixel dimensions 0.92 μm px^-1^, image size 1200 x 1920 px, 16-bit). In addition, a Z-stack encompassing 240 μm with optical sections of 30 μm was carried out to obtain a clear view of the entire root hairs. Finally, an image was recorded every 8 min. The images were analyzed with *Fiji* software. The detailed method illustrated with figures, the macros used for the analysis, as well as a tutorial video to use those macros are available in the **Supplementary Methods, Figure S3, File S5-8 and Movie S4**. To test these macros, an example image is provided in the **Supplementary File S9**. The data produced by *Fiji* were analyzed using *R studio* software. The scripts used for the analysis are available in the **Supplementary File S10**.

## Supporting information

Supplementary file S1. RoPod models

Supplementary file S2-8. ImageJ macro used for image analysis

Supplementary file S9. Semi-automated hair tracking: training material

Supplementary file S10. R scripts used for root hair growth analysis

## Author contributions

A.M. conceived the project and the RoPod design; M.G. and G.G. contributed experimental ideas. A.M. and M.G. performed the experiments. A.M., M.G. and G.G. analyzed the data. A.M., M.G. and G.G. wrote the manuscript. D.W. and S.H. contributed to optimizing chamber design, 3D printing and imaging conditions.

## Acknowledgments

The authors are thankful to Prof. Dr. Karin Schumacher (Centre for Organismal Studies, Heidelberg University) for hosting A.M. during the development of this project and her helpful advice. The authors gratefully acknowledge financial support by a Marie Curie fellowship (MAPoPHAGY, 799433) to A.M., by a Heisenberg Professorship and a research grant by the Deutsche Forschungsgemeinschaft (DFG, GR 4559/4-1, GR 4559/5-1) and funds by Germany’s Excellence Strategy (EXC-2048/1, project ID 390686111) to G.G.

## Competing interests

The authors declare no competing interests.

## Supporting information

**Figure S1**. Dynamic changes in the basal autophagic activity detected in root hair and non-hair cells.

**Figure S2**. Root hair cells accumulate less autophagic bodies than non-hair cells under prolonged ConA treatment.

**Figure S3**. Illustration of root hair tracking procedure.

**Movie S1**. Arabidopsis seedlings grown in a RoPod chamber have normal phenotype.

**Movie S2**. Assessment of root hair length and growth rate in response to sucrose treatment.

**Movie S3**. Application of the RoPod for time-lapse imaging of pHusion-ATG8 in roots treated with autophagy modulators.

**Movie S4**. Tutorial to use the semi-automated hair tracking pipeline.

**Supplementary file S1**. RoPod models

**Supplementary file S2**. Puncta quantification on single time-point data

**Supplementary file S3**. ImageJ macro for puncta quantification on timelapse data

**Supplementary file S4**. ImageJ macro used for puncta quantification in the complete cell volume of hair and non-hair root cells.

**Supplementary file S5**. ImageJ macro for the focusing of the Z-stack acquired during the root hair imaging.

**Supplementary file S6**. ImageJ macro for the stitching of the images acquired during the root hair imaging.

**Supplementary file S7**. ImageJ macro for root hairs selection.

**Supplementary file S8**. ImageJ macro for root hair tracking.

**Supplementary file S9**. Semi-automated hair tracking: training material.

**Supplementary file S10**. R scripts used for root hair growth analysis.

**Supplementary methods:** Protocol for RoPods printing; Arabidopsis growth in RoPods; Puncta quantification on the time-lapse data; Detailed description of root hair tracking algorithm.

## Notes

https://www.alyonaminina.org/ropod

https://github.com/AlyonaMinina/RoPod

### Competing Interest Statement

The authors have declared no competing interest.

### Summary of Updates

Figure 1: RoPod concept better explained for more clarity; Figure 2, New data: Comparison between growth in RoPod and growth using standard methods; Figure 3: Some information of the supplemental data are imported into the main figure; Figure 4 : updated; Figure S2, New data: autophagic puncta quantified at an end time point, throughout the full volume of hair cells and non-hair cells

