## Supplementary figures and methods for "RoPod, a customizable toolkit for non-invasive root imaging, reveals cell type-specific dynamics of plant autophagy"

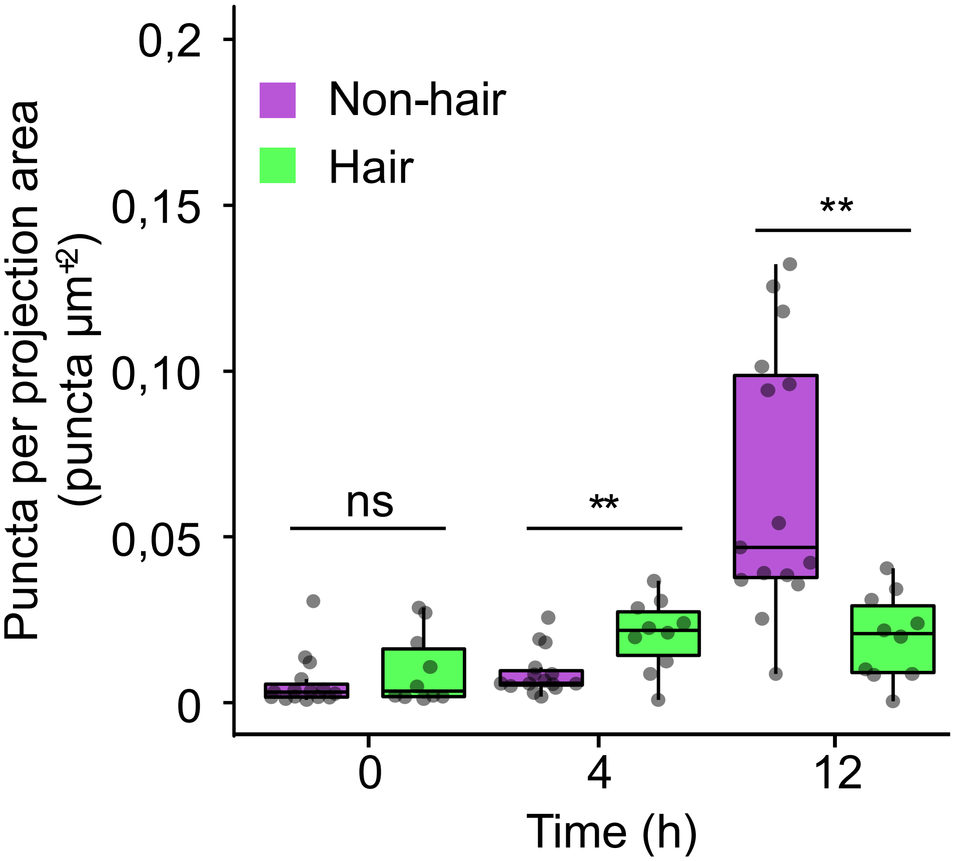


**Figure S1. Dynamic changes in the basal autophagic activity detected in root hair and non-hair cells.** Quantification of pHusion-ATG8-positive puncta per area in the epidermal root cells of WT seedlings grown in RoPods and treated with 500 nM ConA. The data represents 0h, 4h and 12h time points of the time-lapse assays shown in the **Figure 5** and **Movie S3**. T-test, p-value<0.01.

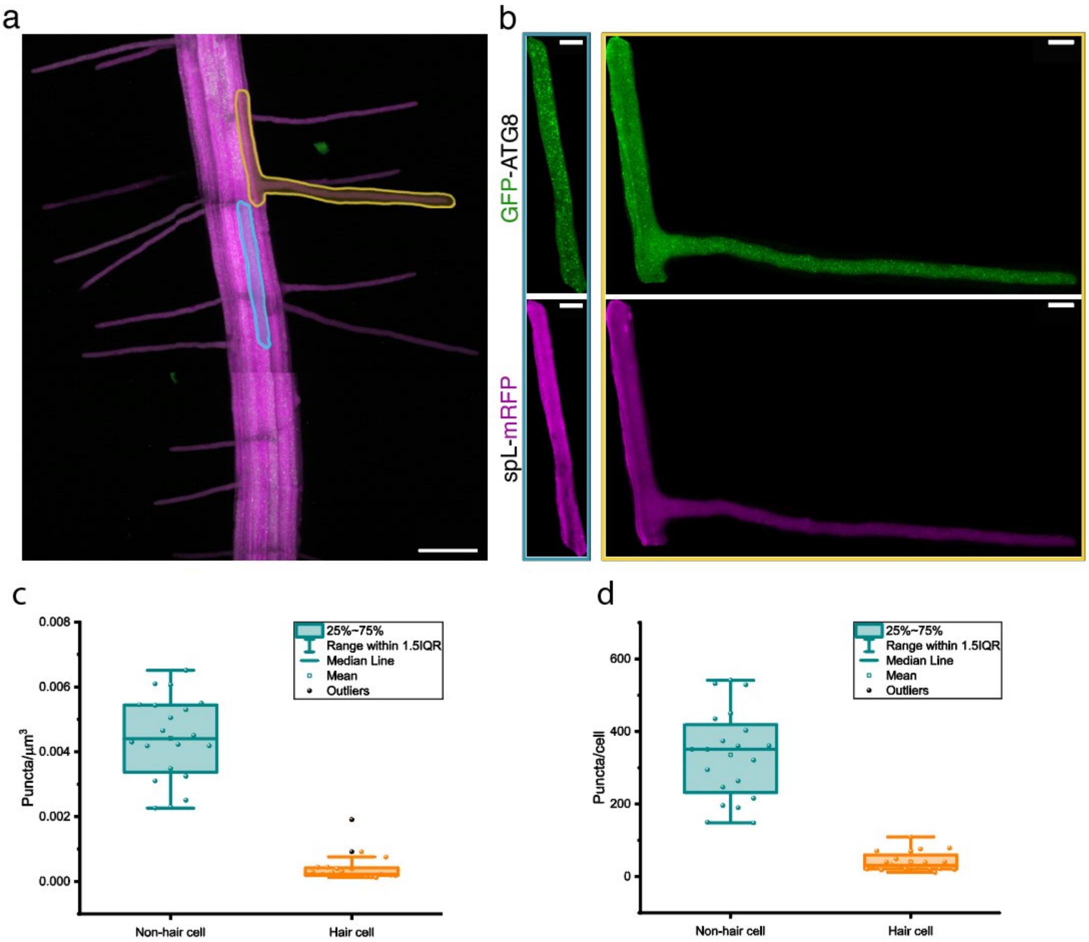


**Figure S2. Root hair cells accumulate less autophagic bodies than non-hair cells under prolonged ConA treatment.**

Arabidopsis seedlings co-expressing autophagy reporter (GFP-ATG8) with vacuolar marker (spL-mRFP) were grown in a RoPod v23.4 and treated with 0.5uM ConA overnight prior to imaging using confocal microscope. (**a**) A representative maximal projection of a tiled z-tack encompassing complete cells with vacuoles was used to segment hair (yellow outline) and non-hair (blue outline) cells; scale bar, 100 µm. (**b**) illustrates segmented non-hair (left) and hair (right) cells outlined in (**a**). Scale bars, 20 µm. The number of GFP-positive puncta was quantified per µm$^3$ (**c**) and per a cell (**d**). Charts in c and d represent data from one out of four independent experiments. Four roots were analyzed, selecting five cells for each cell type for each root, n= 20. OneWay Anova, p-value <0.01 for both charts.

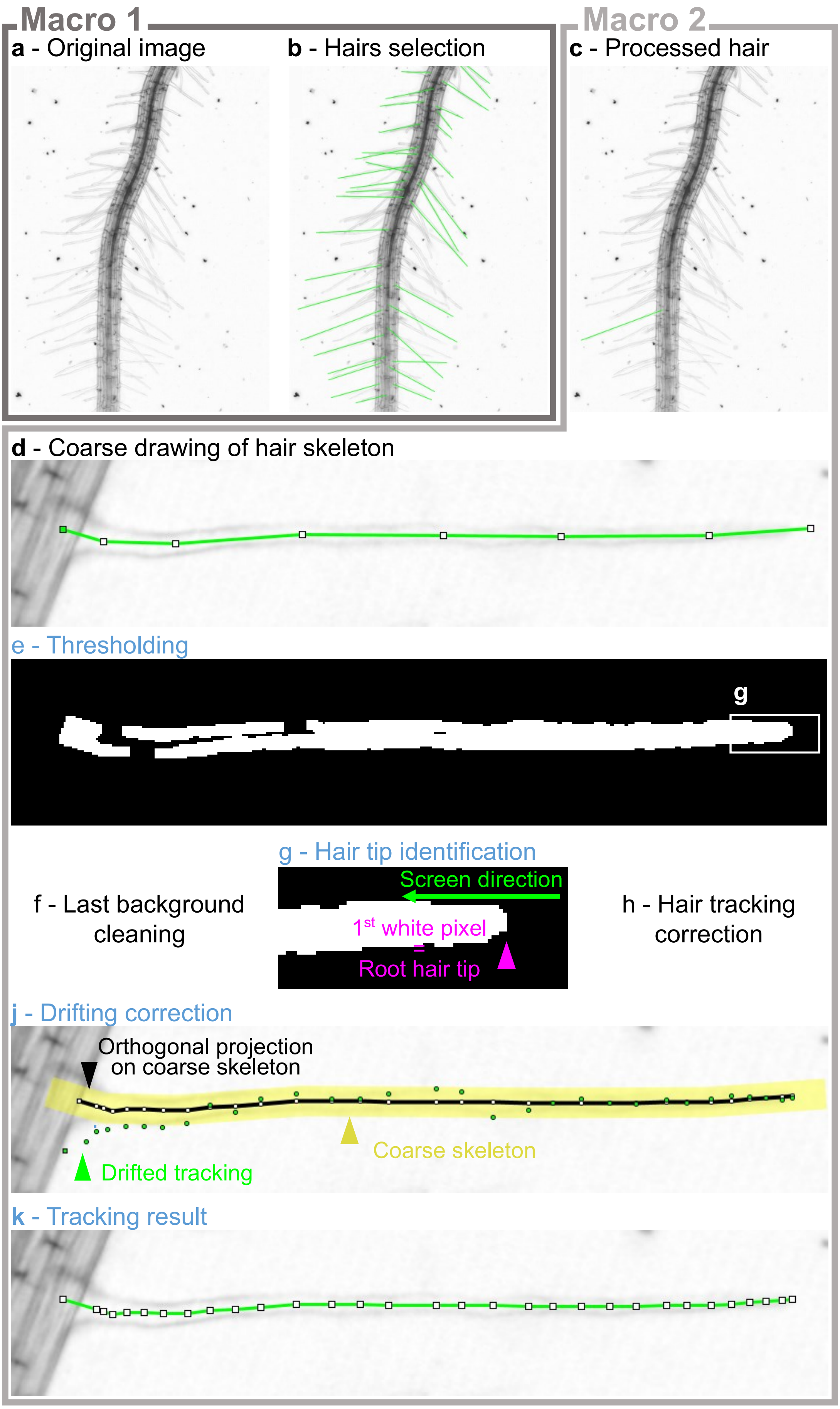


**Figure S3. Illustration of hair tracking procedure.**
(**a-b**) Selection of hairs to analyse with the first Macro. (**c-j**) Hair tracking with the second Macro. The steps indicated in blue are automatic steps. (**g**), zoomed-in inset outlined in (**e**).

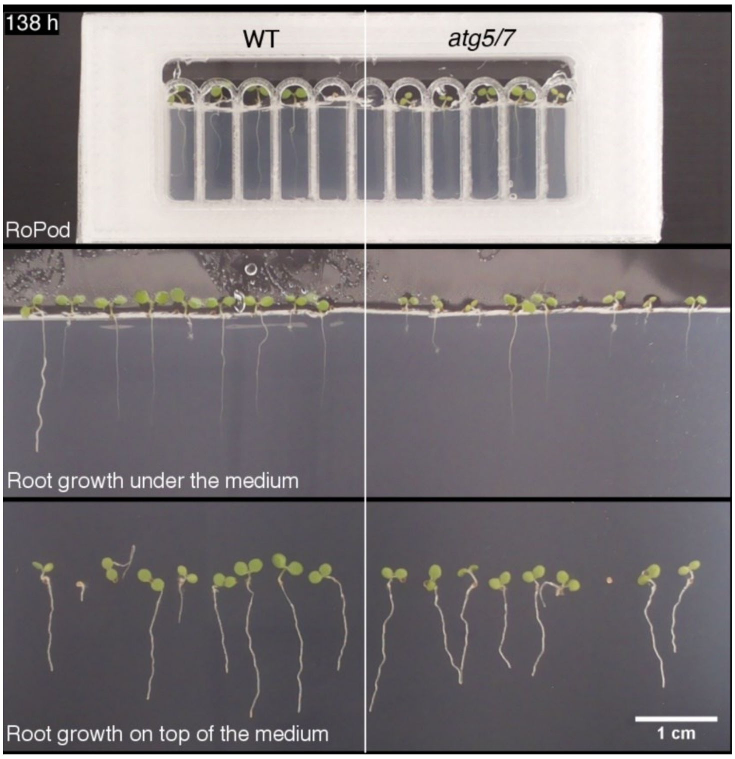


**Movie S1. Arabidopsis seedlings grown in a RoPod chamber have normal phenotype.**
The time-lapse movie of root growth illustrated and quantified in the **Fig.1g-i**. Seed germination and root elongation of seedling grown in the RoPod v24.3 are comparable to the growth under or on top of the growth medium in a Petri plate. Additionally, similar growth of wild-type WT) plants corroborates that RoPod provides favourable growth conditions for Arabidopsis plants. Images were acquired using SPIRO.

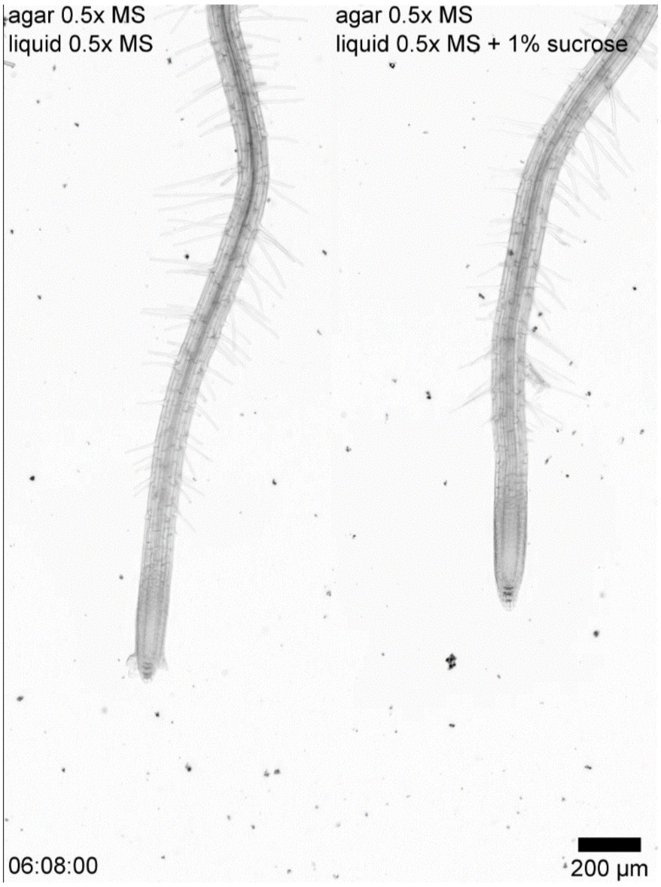


**Movie S2. Time-lapse video of root hair growth response to sucrose treatment.**
Representative video of Col-0 WT seedlings grown in RoPod5 on a vertical microscope and treated with 0.5x MS (control – left panel) or 0.5x MS supplemented with 1 % sucrose (right panel).

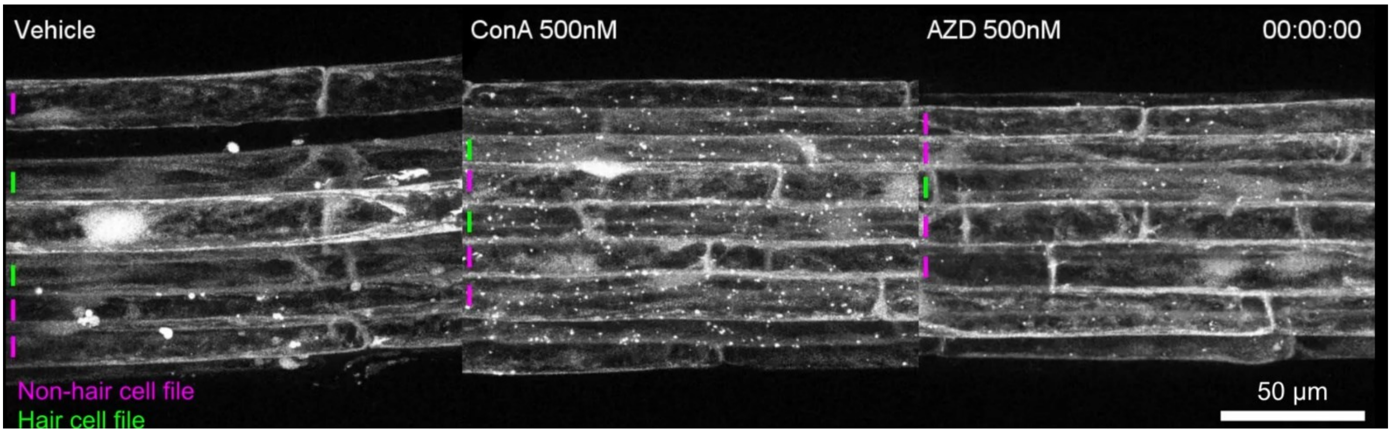


**Movie S3. Application of the RoPod for time-lapse imaging of pHusion-ATG8 in roots treated with autophagy modulators.**
Representative movies of WT seedlings expressing the autophagosomal marker pHusion-ATG8 grown in a RoPod chamber and treated with vehicle (0.01% DMSO), 500 nM Concanamycin A (ConA) or 500 nM AZD8055 (AZD). ConA blocks the final step of autophagy, i.e. degradation of the autophagic bodies in the vacuole, causing massive accumulation of the pHusion-ATG8-positive puncta in the vacuolar lumen. AZD8055 induces autophagic activity, causing incorporation of the ATG8a-based reporter into autophagosomes followed by its delivery to the vacuole and degradation. Note decrease of the fluorescent signal after ca 4h of treatment. The magenta and green lines indicate respectively hair and non-hair cell files used for quantification. Scale bar, 50 µm.

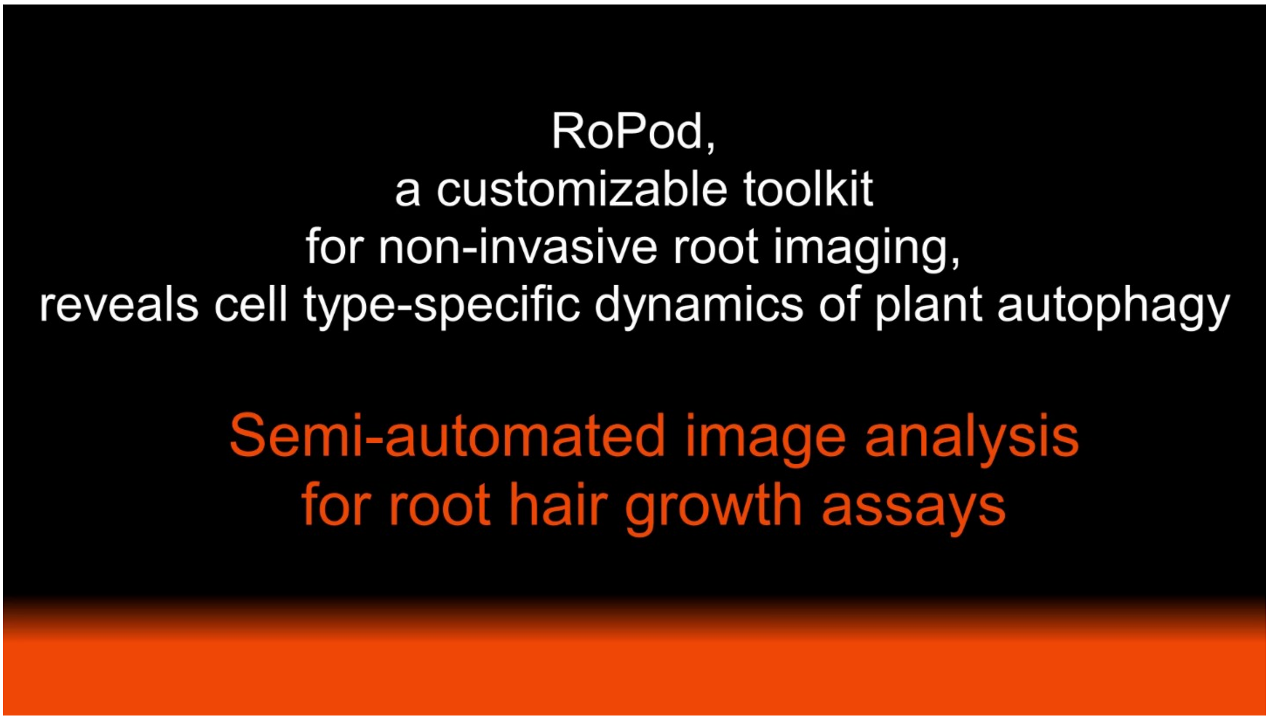


**Movie S4. Tutorial to use the semi-automated hair tracking pipeline**

#### Supplementary methods

##### Protocol for RoPods printing

Please note that updates are published on the website:
https://github.com/AlyonaMinina/RoPod.Hardware

In this video you will find a quick step-by-step demonstration how to print a RoPod

For RoPods 1-5 microscopy cover glass is printed into the plastic, thus resolving the issues with its detachment during experiments and reuse.

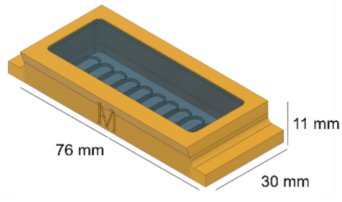


**RoPod 24**

- Optimal for imaging roots of 3-6 days old Arabidopsis seedlings
- Can accommodate up to 11 seedlings
- Separators create lanes for individual root growth
- Arcs on the top of each lane guides young root growth in the desired direction
- Chamber V = ca 6 ml
- Designed for microscopy glass 24 x 60 x 0.17mm (#1.5)

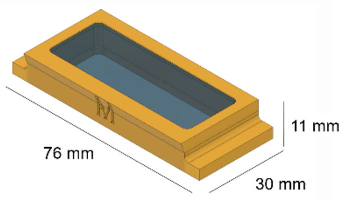


**RoPod 23**

- Optimal for imaging roots of 3-6 days old Arabidopsis seedlings
- Can accommodate up to 20 seedlings
- Chamber V = ca 6 ml
- Designed for microscopy glass 24 x 60 x 0.17mm (#1.5)

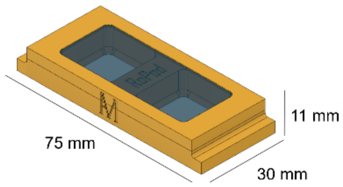


**RoPod 22**

- Optimal for imaging roots of 3-6 days old Arabidopsis seedlings
- Can accommodate 5 seedlings/well. The chamber has been originally designed for plant cell culture and protoplasts imaging
- Well V = ca 3 ml
- The chamber bottom is made out of two 24 x 24mm coverslips, the lid contains one 24 x 60mm coverslip. All coverslips are 0.17mm thick (#1.5)

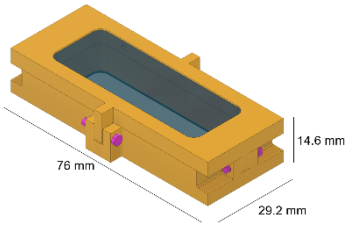


**RoPod 5**

- Optimal for imaging roots of 3-6 days old Arabidopsis seedlings
- Can accommodate up to 15 seedlings
- An insert for an o-ring and clamps are added for proper liquid sealing (a silicon tubing of ~ 2.5mm diameter and ~0.3mm thickness. The ring was close by inserting one extremity of the tube into the other extremity)
- A hole is inserted on top of the box for liquid medium injection
- Chamber V = ca 3 ml
- Designed for microscopy glass 24 x 60 x 0.17mm (#1.5)
  Recommendations for printing: On the lid, force the seam of the print on the top of the chamber (on the same side as the injection hole).
- Suitable for liquid treatment on a vertical microscope

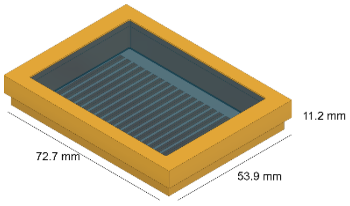


**RoPod 25**

- Optimal for imaging roots of 7 days old Arabidopsis seedlings
- Can accommodate up to 21 seedlings
- Separators create lanes for individual root growth
- Chamber V = ca 7 ml
- Designed for microscopy glass 48 x 64 x 0.17mm (#1.5)
  Recommendations for printing: use a few drops of superglue not only in the corners, but also along the sides of the glass (due to the size of the glass).

**Commercially available chambers tested for RoPod protocol:**

- Ibidi chambered coverslips (cat # 80421 and 80286 #1.5 polymer coverslip, hydrophobic, sterilized, Ibidi, Germany)
- 1-well II Chamber Slide™ System (Nunc™ Lab-Tek™ , cat # C6307, Sigma-Aldrich)
- x-well cell culture chamber, 1-well, on coverglass (cat # 94.6190.102, Sarstedt).

#### For printing RoPods you will need:

- access to a 3D printer extruder with a nozzle 0.4 mm (If you do not have access to a printer, consider finding Makerspace facility in your vicinity)
- PETG filament, white or transparent, ⌀ 1.75 mm
- 24 x 60 mm microscopy coverslip #1.5, i.e. 0.16-0.19 mm thick ( VWR, 630-2108).
- Superglue
- .3mf or .stl file for RoPods.

### Step 1.

- It is highly recommended to generate from the provided .3mf project files. However, we also provide .stl files
- If you prefer to use .stl, set the following parameters:
- follow the instructions for the chamber positioning
- Print settings-> Layer height = 0.1 mm

If possible, we recommend using variable layer height. Variable layer height feature (available for Prusa printers) allows to use small layer height beneficial for efficient embedding of the glass into the print and a larger layer height for printing the rest of the structure. This speed up the overall printing time.

- Print settings-> Fill density = 50%
- Print settings-> Fill pattern = 3D Honeycomb
- Print settings-> Layers and Perimeters-> Vertical shells-> Perimeters = 4
- Slice the model and insert a pause after 5 mm height
- You can simultaneously print up to 17 RoPods on a Prusa MK2or MK3. However we strongly recommend to start with just one.

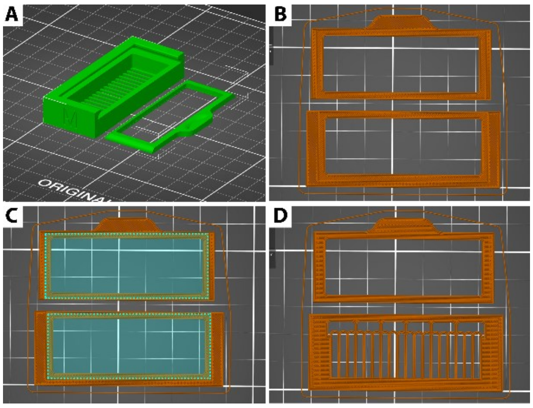


**A**. The RoPod 5 chamber and the lid should both be upside up on the print bed.
**B**. The layer with the slot for glasses should be the last one before the pause
**C**. Slot for the glasses is highlighted with the dotted line
**D**. The very next layer after the paused layer

#### Step 2. Clean the coverslip glasses

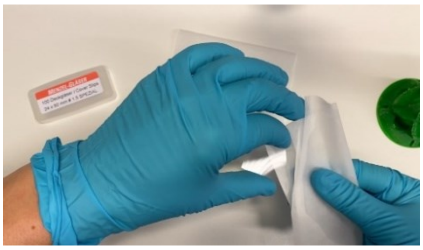

- Remove dust and fat from the glasses by wiping them clean with a paper tissue soaked in acetone

#### Step 3. Start the print

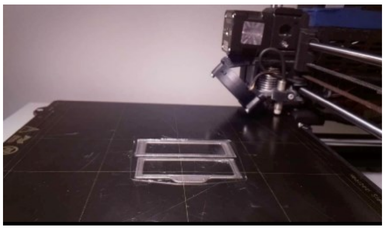

- Start the print and wait until it automatically pauses on the layer programmed int the Step 1.
- The extruder will move to the back left of the print bed while waiting for the user to resume the print

#### Step 4. Place the glasses into the slots

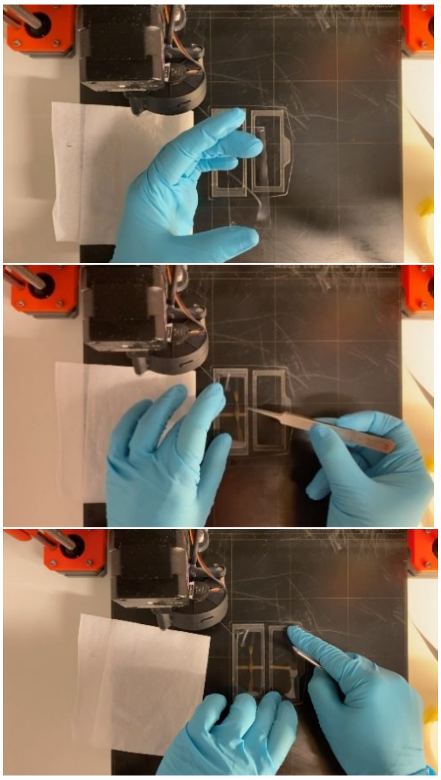

- The chambers and lids models have 0.1mm deep slots for the coverslips
- Add a drop of super glue to two corners of a glass. This will secure the glass' position during printing
- Use forceps and patience to carefully place the glass into its slot. Make sure it does not stick upwards or sideways
- Press down on the the corners with glue and hold for a few seconds. sic! wear gloves! supergluing oneself to a 90C- hot print bed is not a good way to start your day

#### Step 5. Resume the print

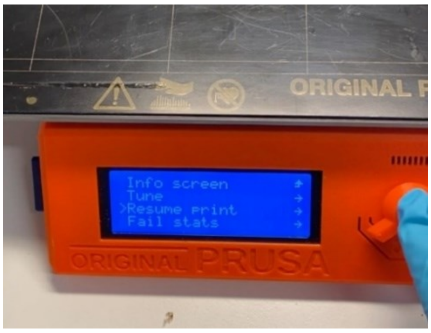


- Resume the print and make sure the the filament does not move the glass from its position and does not crack it
- When the chamber is ready make sure there is no filament sticking sideways. If needed imperfections can be removed with a scalpel

#### Protocol for Arabidopsis growth in RoPods

Shown here on the example of RoPod 24

Please note that updates are published on the website: https://www.alyonaminina.org/ropod

In this video you can find a short step-by-step demonstration of how to prep RoPod chambers and grow plants in them.

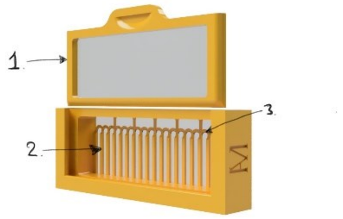


**RoPod 24 has**:
1. A watertight lid that locks by curving out. To unlock the lid, gently push its handle upwards and slide the lid inside its slot
2. Arcs to guide roots into the channels
3. Separators to prevent neighboring roots from crossing and create "lanes" for individual root growth.

### Step 1.

Sterilize the RoPod under UV in a laminar flow hood. Place the open chamber and the lid (inner side up) under the UV light, 30 minutes of exposure is usually sufficient.

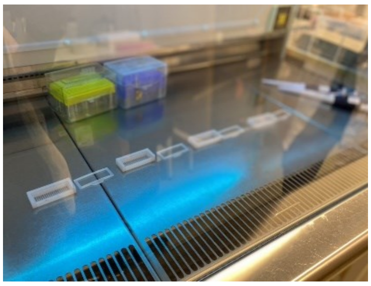

Chambers can also be sterilized by submerging in 70% Ethanol for 30 minutes and then rinsing in 95% Ethanol and leaving to air dry in a laminar flow hood. However, in this case it is crucial to ensure that ethanol completely evaporated as even traces of it will have adverse effect on seedling growth.

### Step 2.

Thaw 0.5x MS in a microwave, let it cool to 60 C and pipette 3mL into the sterile chamber. Make sure the medium is evenly distributed in the chamber and there are no bubbles

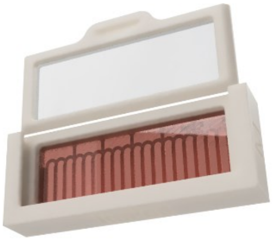

**Recommended V of the medium:**
1mL of medium per a well of the 3 –well chamber, i.e. RoPod 3
1.5 mL well of a 2-well chamber, i.e. RoPod 2
3 mL/ 1-well chamber, i.e. RoPod 5, 24 and 25

**0.5x MS:**
0.5x MS including vitamins (M0222, Duchefa)
10 mM MES (M1503, Duchefa)
1% sucrose
0.8 % Plant agar (P1001, Duchefa)
Adjust pH with KOH to 5.8
Autoclaved for 20 minutes at 120°C
Store at 4C

#### Step 3.

Let the medium solidify and then use a tip or toothpick to remove a ca 5 mm wide strip of the medium, this will give space for the areal part of seedlings. For RoPod 5 make sure arcs are exposed.

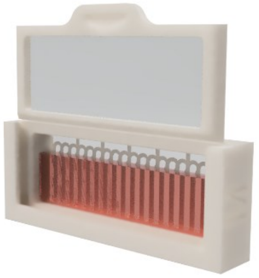

#### Step 4.

Transfer sterile seeds using a sterile toothpick onto the bottom glass by the edge of the medium. For RoPod 5 place the seeds “under” the arches, which are designed to guide roots of young seedling into the individual channels.

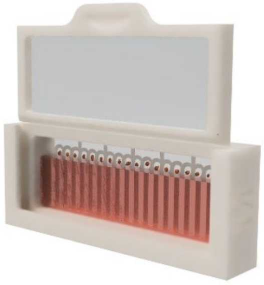

Seeds can be surface sterilized in 70% EtOH/0.05% TritonX-100 for 20 minutes, washed three times with 96% EtOH and air dried in a laminar flow hood. Sterile seeds can be stored for up to half a year.
Please note, that RoPod chambers with printed in glass have a 0.5mm plastic rim surrounding the coverslip glass, which might not allow to image with immersion objectives very close to the edges of the glass window. If you plan to use 40x and higher magnifications, it is best to not image in the first and last lanes of RoPod 5.

#### Step 5.

Close the chamber. RoPods with watertight lids do not need to be sealed individually, other RoPods require it. Place the chamber into a square Petri plate and seal it.

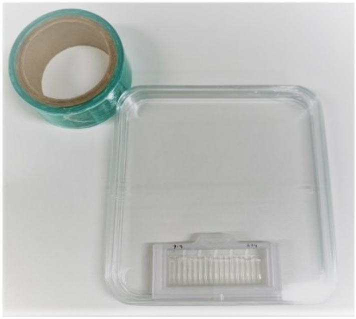

#### Step 6.

Place the plate under growth conditions, vertically or slightly tilted to guide root growth towards the coverslip bottom

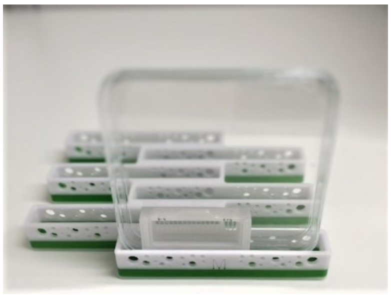

#### Puncta quantifications on the time-lapse data

detailed description of the image analysis protocol used for the autophagic puncta quantification in the time resolved data set (Figure 5)

Raw confocal image stacks were imported into *Fiji*. A maximum-intensity projection of the GFP channel was generated, the data set was rotated to orient the root horizontally and cropped. The data set was registered using the *MultiStackReg* plugin with "rigid body" settings, starting at the final slide to correct for root growth and specimen movement during imaging. The registered data sets were cropped again to the maximum area available for continuous analysis throughout the time course. ROIs were defined along hair and non-hair cells, excluding cell borders, and data sets were exported as TIFF stacks for further analysis. Autophagosomes were then identified using pixel classification approaches with the software *Ilastik*. The classifier was trained on five images (Col-0 roots treated with: DMSO, ConA, AZD; and two *atg5-1/atg7-2* roots treated with AZD). Pixel features analyzed by the software were defined according to their intensity (Gaussian smoothing with a radius of 0.3 and 0.7 pixel) and their edges (Laplacian Gaussian with of 0.7 and 1.0, and Gaussian Gradient Magnitude) with radius of 0.7). Three classes were defined for the training: vesicles, root and background. The classifier was trained by manually brushing areas of interest for each defined class. The simple segmentation produced was exported as 8-bit images in order to be processed further. In *Fiji*, regions of interest were defined to identify the hair and non-hair cell files to analyze. For each cell file ROI, its size, vesicles number, vesicles fluorescence intensity and vesicles size were quantified using an algorithm run in *Fiji* (**Supplementary File S2**).

#### Detailed description of root hair tracking helping algorithm and corresponding data analysis description

For data presented in figure 4

The images were analyzed with *Fiji* software.

To obtain a fully focused single image from the Z-stack, the macro from the **Supplementary File S5** was used. In a nut shell, the macro is using an edge detection filter and, for each subregion of the image in the XY direction, will keep for the final image only the sharpest subregion in Z direction. The algorithm has been implemented to be run on a GPU so it can run faster.

The background of the images was then corrected with the "*Substract background*" tool, and the complementary field of view of each root were then stitched with the "*3D Stitching*" plugin using the macro from the **Supplementary File S6**.

The hairs to be analyzed were then selected by drawing straight lines starting from their base and extending over the entire length of each of them (**Supplementary Fig. S3b**). A *Fiji* macro to guide this step is available (**Supplementary File S7, Movie S4**, https://github.com/AlyonaMinina/RoPod).

Then, an algorithm run in *Fiji* was used to facilitate the tracking of the tip of the hair (**Supplementary File S8, Fig. S3c-k, Movie S4**, https://github.com/AlyonaMinina/RoPod). This algorithm consists in using the straight lines defined previously and the *Straighten* tool to rotate the root hair in a horizontal position, the tip facing the right side of the image (**Supplementary Fig. S3d**). Then, a midline is drawn by the user throughout the hair (**Supplementary Fig. S3d**). This midline is used to create a mask around it in order to prevent other hairs from being processed by the algorithm (**Supplementary Fig. S3e**). A threshold is applied with the *MinError* of the *Treshold* window to obtain a white image of the root hair on a black background (**Supplementary Fig. S3e**). The white pixel positioned in the rightmost part of the image is the tip of the hair (**Supplementary Fig. S3g**). It is then important that the background is kept clear on the rightmost part of the root hair tip. For that purpose, a rough manual cleaning of the background is proposed in the algorithm (**Supplementary Fig. S3f**). This method is not suitable for monitoring hair growth for the first few minutes after the root hair bulge emergence. This is due to the thresholding method which does not differentiate between the root hair and the root body. For this reason, a manual adjustment step is proposed to correct any errors (**Supplementary Fig. S3h**). At the beginning of root hair growth, positions of bulges and young root hairs drift as the root finishes its elongation. Therefore, the absolute coordinates of the root hair tip between two time points do not only reflect the displacement of the root hair. To correct this drift, the coordinates of the root hair tip were projected orthogonally on the medium line previously drawn by the user (**Supplementary Fig. S3j-k**). The coordinates of the projected root hair tips were then used to calculate root hair displacement over time. All the data were exported into a table saved in .xls format and the root hair tip tracks were saved as ROI files.

For the data analysis in R (**Supplementary File S10**), the final root hair length and the growth duration were calculated by fitting each root hair growth profile with the sigmoidal curve $f(t)$:

$$f(t) = L_{min} + \frac{L_{max} - L_{min}}{1 + e^{(t_{50}-t)/\delta}}$$

$L_{min}$: basal line, $L_{max}$: maximum hair length, $t_{50}$ inflexion point, δ: slope factor at inflexion point.

The fitted parameters $L_{max}$ and $[t_{50} + 2x|2.5\delta|]$ were used as read out to calculate the final hair length and the growth duration respectively. The root hair growth rate was calculated during the linear growth phase, defined between 50 min after the emergence of the hair bulge and the time at which the hair does not grow more than 20 µm for 40 consecutive minutes.
