## Supplementary figures and images for "RoPod, a customizable toolkit for non-invasive root imaging, reveals cell type-specific dynamics of plant autophagy"

### 20220322_Col_sucrose_Vert_RoPod5_B2_R10_EDF-RW-GPU_Bckg-Stitch-reg_2.tif

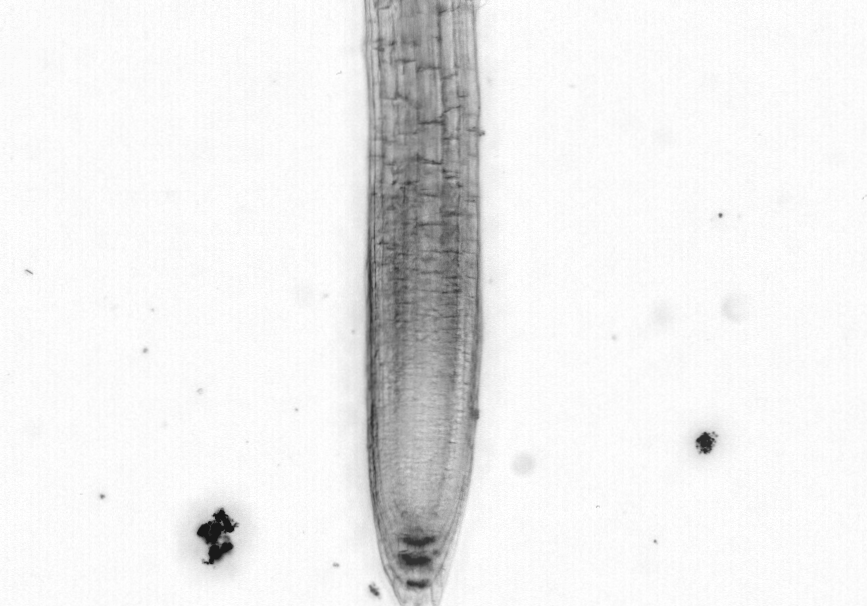

### 20220322_Col_sucrose_Vert_RoPod5_B2_R10_EDF-RW-GPU_Bckg-Stitch-reg_2_roi_01_Straigthen-BF.tif

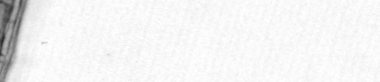

### 20220322_Col_sucrose_Vert_RoPod5_B2_R10_EDF-RW-GPU_Bckg-Stitch-reg_2_roi_02_Straigthen-BF.tif

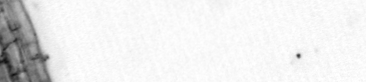

### 20220322_Col_sucrose_Vert_RoPod5_B2_R10_EDF-RW-GPU_Bckg-Stitch-reg_2_roi_03_Straigthen-BF.tif

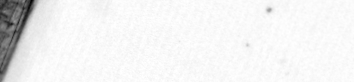

### 20220322_Col_sucrose_Vert_RoPod5_B2_R10_EDF-RW-GPU_Bckg-Stitch-reg_2_roi_04_Straigthen-BF.tif

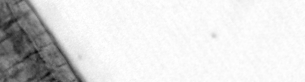

### 20220322_Col_sucrose_Vert_RoPod5_B2_R10_EDF-RW-GPU_Bckg-Stitch-reg_2_roi_05_Straigthen-BF.tif

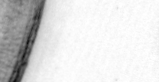

### RoPod5-2_Supl-preview_5.png

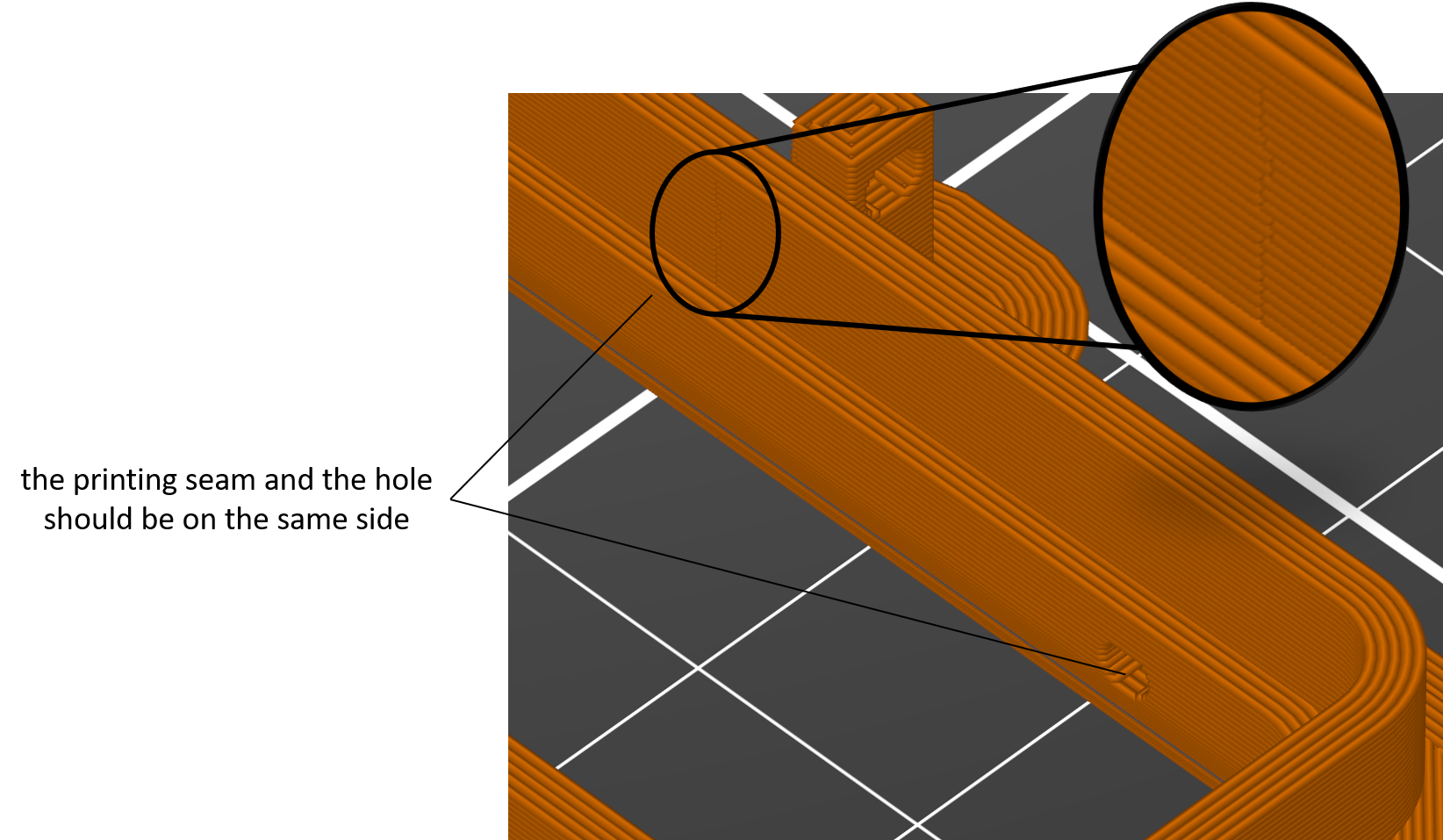
