## Supplementary file S2-8. ImageJ macro used for image analysis for "RoPod, a customizable toolkit for non-invasive root imaging, reveals cell type-specific dynamics of plant autophagy": Ropod 2022_Supplementary file S2.ijm

```
//The number of puncta in root epidermal cells was quantified in a semi-automated manner using Fiji. The following steps were achieved:
run("Split Channels"); //only GFP channel was used for puncta quantification
run("Make Substack...", "  slices=2-25"); //only epidermal cells were included into the sub stack. The number of z-slices was recorder manually for quantifying the Volume later. The z-depth of the whole stack was lifted from the file metadata in ZEN => depth of the sub stack = (total z-depth of the file/30 slices) * number of slices included into the sub stack
run("Z Project...", "projection=[Standard Deviation]"); //Standard deviation gave least noisy z-projection
run("Measure"); //a ROI was hand drawn using polygonal tool to exclude root hairs (those might significantly increase overestimation of the V) => the area was measured. Z volume in µm3 = area*z-depth of the sub stack
run("Find Maxima..."); // the number of puncta was measured using find maxima stringency =20
```
