## Supplementary file S2-8. ImageJ macro used for image analysis for "RoPod, a customizable toolkit for non-invasive root imaging, reveals cell type-specific dynamics of plant autophagy": Ropod 2022_Supplementary file S3.ijm

```
/*
 * AIM OF THE MACRO
 * 	Each measurement presented bellow are done in a ROIs defines previously by the user
 * 	Measure the mean gray value and the size of segmented elements (for example obtained with the software "Ilastik")
 * 	Measure the mean gray value of a fix round area around the middle of each segmented elements
 * 	Produce a clean result table ready to be saved
 * 	
 * BEFORE RUNNING THE MACRO
 * 	open the raw fluorescent image
 * 	open the associated segmented image
 * 	open the ROI file 
 * 	change in the code the lines label with "****" according to the needs * 	
 *	
 *CREDIT
 * Marjorie Guichard, Grossmann lab, Center for Organismal Studies, Heidelberg, Germany, 07/04/2020
 *
 */
 
//fix vesicle diameter definition
FixVesicleDiameterMicron = 2; // **** Set the region to measure around each vesicle (in micron)

//Define the images to analyse
run("Tile");
waitForUser("Select the segmented image");
SegmentedImageID = getImageID();
waitForUser("Select the NON segmented image");
NonSegmentedImageID = getImageID();
NonSegmentedImageName = getTitle();
dotIndex = indexOf(NonSegmentedImageName, ".tif");
NonSegmentedImageRootName = substring(NonSegmentedImageName, 0, dotIndex);

//get the voxel size of the image and convert the custom vesicle diameter
getVoxelSize(VxWidth, VxHeight, VxDepth, VxUnit);
FixVesicleDiameterPx = FixVesicleDiameterMicron/VxWidth;

//Extract the vesicles from the segmented images
selectImage(SegmentedImageID);
run("Select None");
run("Duplicate...", "duplicate");
SegmentedThresholdedImageID = getImageID();
setThreshold(3, 3); // ***** should be change if the gray value attributed for the vesicles at the segmentation step is different
setOption("BlackBackground", true);
run("Convert to Mask", "method=Default background=Dark black");

//Define the arrays required to store informations related to the epidermis cell files ROIs
EpROIsCoordX = newArray();
EpROIsCoordY = newArray();
EpROIsCoordStartIndex = newArray();
EpROIsCoordStopIndex = newArray();
EpROIsName = newArray();
for (EpidermisROIToAnalyse = 0; EpidermisROIToAnalyse < roiManager("count"); EpidermisROIToAnalyse++) {
	roiManager("select", EpidermisROIToAnalyse);
	EpROIsName = Array.concat(EpROIsName,Roi.getName);
	getSelectionCoordinates(xpoints, ypoints);
	EpROIsCoordStartIndex = Array.concat(EpROIsCoordStartIndex, lengthOf(EpROIsCoordX));
	EpROIsCoordX = Array.concat(EpROIsCoordX,xpoints);
	EpROIsCoordY = Array.concat(EpROIsCoordY,ypoints);
	EpROIsCoordStopIndex = Array.concat(EpROIsCoordStopIndex, lengthOf(EpROIsCoordX));
}

//set the arrays required to fill the final table
run("Set Measurements...", "area mean stack display redirect=None decimal=2");
RawVesiclesMeanGrayValue = newArray();
RawVesiclesSize = newArray();
RawVesiclesSlice = newArray();
RawVesicleImageName = newArray();
RawVesicleEpROIName = newArray();
RawVesicleEpROISize = newArray();
MiddleVesicleMeanGrayValue = newArray();
MiddleVesicleSize = newArray();


setBatchMode(true);
//loop to process each cell files
for (EpidermisROIToAnalyse = 0; EpidermisROIToAnalyse < lengthOf(EpROIsName); EpidermisROIToAnalyse++) {
	//recover the name of the current cell file
	EpidemisROICurrentName = EpROIsName[EpidermisROIToAnalyse];
	
	//select the cell file
	selectImage(NonSegmentedImageID);
	run("Clear Results");
	roiManager("deselect");
	roiManager("delete");
	EpCurrentROICoordX = Array.slice(EpROIsCoordX,EpROIsCoordStartIndex[EpidermisROIToAnalyse],EpROIsCoordStopIndex[EpidermisROIToAnalyse]);
	EpCurrentROICoordY = Array.slice(EpROIsCoordY,EpROIsCoordStartIndex[EpidermisROIToAnalyse],EpROIsCoordStopIndex[EpidermisROIToAnalyse]);
	makeSelection("Polygon", EpCurrentROICoordX, EpCurrentROICoordY);	
	
	//measure the cell file area
	run("Measure");
	EpROIsSize = getResult("Area", 0);	
	run("Clear Results");//some cleaning

	//create a segmented vesicles image without the vesicles outside from the cell file of interest 
	selectImage(SegmentedThresholdedImageID);
	run("Select None");
	run("Duplicate...", "duplicate");
	ImagesToDelete = newArray(); // for future cleaning
	ImagesToDelete = Array.concat(ImagesToDelete, getImageID()); // for future cleaning
	makeSelection("Polygon", EpCurrentROICoordX, EpCurrentROICoordY);
	run("Make Inverse");
	setBackgroundColor(0, 0, 0);
	for (sliceToProcess = 0; sliceToProcess < nSlices; sliceToProcess++) {
		setSlice(sliceToProcess+1);
		run("Clear", "slice");
	}
	run("Select None");//some cleaning

	// convert the vesicle images into ROIs and identify the number of ROIs created
	run("Analyze Particles...", "add stack");
	vesiclesROIsToSelect = newArray();
	for (vesiclesROIIndex = 0; vesiclesROIIndex < roiManager("count"); vesiclesROIIndex++) {
		vesiclesROIsToSelect = Array.concat(vesiclesROIsToSelect, vesiclesROIIndex);
	}
	
	// Measure the area and mean gray value of the vesicles in the cell file of interest
	selectImage(NonSegmentedImageID);
	roiManager("Select", vesiclesROIsToSelect);
	roiManager("Measure");

	// fill the arrays that will be used to create the final result table
	for (vesiclesROIIndex = 0; vesiclesROIIndex < roiManager("count"); vesiclesROIIndex++) {
		RawVesiclesMeanGrayValue = Array.concat(RawVesiclesMeanGrayValue, getResult("Mean", vesiclesROIIndex));
		RawVesiclesSlice = Array.concat(RawVesiclesSlice, getResult("Slice", vesiclesROIIndex));
		RawVesiclesSize = Array.concat(RawVesiclesSize, getResult("Area", vesiclesROIIndex));
		RawVesicleImageName = Array.concat(RawVesicleImageName, NonSegmentedImageName);
		RawVesicleEpROIName = Array.concat(RawVesicleEpROIName, EpidemisROICurrentName);
		RawVesicleEpROISize = Array.concat(RawVesicleEpROISize,EpROIsSize);
	}	
	
	//Measure the mean gray value in a fix region around each vesicle middle
	run("Clear Results");
	for (vesiclesROIIndex = 0; vesiclesROIIndex < roiManager("count"); vesiclesROIIndex++) {
		roiManager("select", vesiclesROIIndex);
		getSelectionBounds(x, y, width, height);
		makeOval(x+(width/2)-(FixVesicleDiameterPx/2), y+(height/2)-(FixVesicleDiameterPx/2), FixVesicleDiameterPx, FixVesicleDiameterPx);
		run("Measure");		
		// Fill the arrays used to fill the final result table
		MiddleVesicleMeanGrayValue = Array.concat(MiddleVesicleMeanGrayValue, getResult("Mean", 0));
		MiddleVesicleSize = Array.concat(MiddleVesicleSize,getResult("Area", 0));

		//some cleaning
		run("Clear Results");
	}	
	
	// Some cleaning
	for (ImageToDelete = 0; ImageToDelete < lengthOf(ImagesToDelete); ImageToDelete++) {
		selectImage(ImagesToDelete[ImageToDelete]);
		close();
	}	
}
setBatchMode("exit and display");

Table.create(NonSegmentedImageRootName+"-Vesicles analysis");
Table.setColumn("Image Name", RawVesicleImageName);
Table.setColumn("Epidermis ROI name", RawVesicleEpROIName);
Table.setColumn("Epidermis ROI size (µm²)", RawVesicleEpROISize);
Table.setColumn("Slice", RawVesiclesSlice);
Table.setColumn("Segmented Vesicle Size (µm²)", RawVesiclesSize);
Table.setColumn("Segmented Vesicle Mean Gray Value", RawVesiclesMeanGrayValue);
Table.setColumn("Fix Vesicle Size (µm²)", MiddleVesicleSize);
Table.setColumn("Fix Vesicle Mean Gray Value", MiddleVesicleMeanGrayValue);
Table.update;

// Some cleaning
roiManager("deselect");
roiManager("delete");
run("Clear Results");
selectImage(SegmentedThresholdedImageID);
close();

//The end
showMessage("Tada! You can save the result table");
```
