## Supplementary file S2-8. ImageJ macro used for image analysis for "RoPod, a customizable toolkit for non-invasive root imaging, reveals cell type-specific dynamics of plant autophagy": Ropod 2022_Supplementary file S4.ijm

```
//This is a semi-automated analysis pipline, desgined for qunatifying autophagic bodies in the whole-cell volume using tiled z-tack .czi image files.
//The procedure comprises 5 steps:
//1. Creating a overview files to aid selction of cells for analysis
//2. Use overview files to make masks for cell segmentation
//3. Manually segment individual cells
//4. Use the red channel for volume measurement
//5. Use the green channel for puncta quantification


//Clear the log window if it was open
if (isOpen("Log")){
	selectWindow("Log");
	run("Close");
	}

//1. Creating a overview files to aid selction of cells for analysis
//Pre-process the tiled z-stacks
// Macro to process the tile z-stacks
// Pick the directory with the czi images
original_dir = getDirectory("Choose directory");
 
//List all files in the directory with original files
list = getFileList(original_dir);

//create a shorter list contiaiing . czi files only
czi_list = newArray(0);
for(z = 0; z < list.length; z++) {
	if(endsWith(list[z], ".czi")) {
	czi_list = Array.concat(czi_list, list[z]);
		}
	}
print("");		
print(czi_list.length + " images were detected for analysis");


// Loop through the list of .czi files, excluding subfolders
for (i = 0; i < czi_list.length; i++){
	path = original_dir + czi_list[i];
		if (File.isFile(path)){
		print(" ");
		print("Processing image " + i+1 + " out of " + czi_list.length);
		run("Bio-Formats Windowless Importer",  "open=path");
		      
	 	// image name
		title = getTitle;
		 
		// remove redundant info from the name
		short_name = substring(title, 0, 26);
		 
		// create subfolder to save segmented cells
		sub_dir = original_dir + short_name;
		File.makeDirectory(sub_dir);
		 
		// split channels for individual adjustments
		run("Split Channels");
		 
		// prettify the GFP channel
		selectWindow("C1-" + title);
		rename("GFP-ATG8");
		run("Gaussian Blur 3D...", "x=0.5 y=0.5 z=0.5");
		 
		// prettify the RFP channel
		selectWindow("C2-" + title);
		rename("spL-RFP");
		run("Remove Outliers...", "radius=5 threshold=50 which=Bright stack");
		run("Gaussian Blur 3D...", "x=0.5 y=0.5 z=0.5");
		 
		// combine the two fluorescent channels. DIC is too messy for segmentation
		run("Merge Channels...", "c2=[GFP-ATG8] c6=[spL-RFP]");
		 
		// save the tiff file into the corresponding subfolder
		saveAs(sub_dir + "/" + short_name);
		 
		//create and save an overview file
		run("Z Project...", "projection=[Standard Deviation]");
		saveAs(sub_dir + "/" + "Overview " + short_name);
		run("Close All"); 
		}
	}
	
 waitForUser("Please continue further processing by manually. Click "Cancel" and proceed by selecting code lines and pressing Ctrl+Shift+R"); 

//2. Use the overview file to label hair and non-hair cells to be processed
//create a list of overciew files only


//3. Manually segment individual cells
// open the stack file and draw an outline of a cell using an overly brush
 
//create a selection based on the outline
run("Duplicate...", " ");
run("Flatten");
 
 setTool("wand");
//select the ROI and add it to the ROI manager
 
//remove the overlay from the original stack
run("Remove Overlay");
  
//cropped out the outlined cell
roiManager("Select", 0);
run("Duplicate...", "duplicate");
run("Clear Outside");
 
//remove irrelevant slices
run("Make Substack...", "channels=1-2 slices=1-13");
  
//save the segmented cell image
 
//split the channels
run("Split Channels");

//4. Use the red channel for volume measurement and green channel for puncta quantification
 //run manually the volume measurement using the red channel -> Analyze-> 3D object counter-> adjust the threshold
 
 //5. Use the red  green channel for puncta quantification
//make a projection and maxima search on the green channel
run("Z Project...", "projection=[Standard Deviation]");
run("Find Maxima...", "prominence=20 output=Count");
```
