## Supplementary file S2-8. ImageJ macro used for image analysis for "RoPod, a customizable toolkit for non-invasive root imaging, reveals cell type-specific dynamics of plant autophagy": Ropod 2022_Supplementary file S5.ijm

```
//Convert the strategy established by R. Wheeler in an imageJ macro running on CPU into an imageJ macro running in GPU
//Principle:
//In focus regions have sharper detail, therefore have a stronger edge detection result.
//The slice with the maximum edge detection result is most in focus.
//Build up the image from the original stack in patches according to the most in focus slice.
//Use gaussian blending to reduce the appearance of sharp edges.
//This is similar to the plugin by Michael Umorin:
//http://rsbweb.nih.gov/ij/plugins/stack-focuser.html
// input:  grayscale Z stacks with several time frames, NOT organised in hyperstack (organised in Zstrack/Time order, NOT Time/Zstack)
//Process all images contained in oe specified folder and save the processed images in another specified folder
// Marjorie Guichard - 08/04/2022 - Insitut für Zell und Interaktion Biologie - Pr. Dr. Guido Grossmann - HHU Düsseldorf - Germany -

//initialise GPU
run("CLIJ2 Macro Extensions", "cl_device=[GeForce GTX 1050]");
Ext.CLIJ2_clear();

//Get EDF parameters
UserMaxRadius = 1;
UserBlurSigma = 5;

//Stack parameters
//TotalTimeFrame = 39; // for folder "20220322_Col_sucrose_Vert_RoPod5_02_1"
TotalTimeFrame = 49; // for folder "20220322_Col_sucrose_Vert_RoPod5_01_1"
SlicePerFrame = 9;

//Input and output selection
inDir = getDir("Get INPUT directory");
outDir = getDir("Get OUTPUT directory");
filelist = getFileList(inDir);
for (FileToProcess = 0; FileToProcess < lengthOf(filelist); FileToProcess++) {
    if (endsWith(filelist[FileToProcess], ".tif")) { 
        open(inDir + File.separator + filelist[FileToProcess]);
        //get simple name
        origtitle=getTitle();
        dotIndex = indexOf(origtitle, ".ome.tif");
        RootImageName = substring(origtitle, 0, dotIndex);

		//Get image information
		run("8-bit");
		OriTitle = getTitle();
		OriSliceNumber = nSlices;
		OriWidth = getWidth();
		OriHeight = getHeight();
		OriBit = bitDepth();
		getVoxelSize(VxWidth, VxHeight, Vxdepth, Vxunit);
		
		// Create the image where the EDF of each time frames will be stored
		newImage("Untitled", OriBit+"-bit white", OriWidth, OriHeight, 1);
		rename("Final");
		
		//Process each time frame
		setBatchMode(true);
		for (FrameToProcess =0; FrameToProcess < TotalTimeFrame; FrameToProcess++) {
			showProgress(FrameToProcess, TotalTimeFrame);
			// extract the slices for this time frame
			selectWindow(OriTitle);
			run("Duplicate...", "duplicate range="+FrameToProcess*SlicePerFrame+1+"-"+FrameToProcess*SlicePerFrame + SlicePerFrame);
			UnFocusFrame = "UnFocusFrame";
			rename(UnFocusFrame);
			
			//run EDF
			//________________________
			//load Frame to process
			Ext.CLIJ2_push(UnFocusFrame);
			TempEDF = "FinalEDFOnTimeFrame";
			//Sobel filter on GPU
			SobelStack = "Sobel";
			Ext.CLIJ2_sobelSliceBySlice(UnFocusFrame, SobelStack);
			
			// Create max filter on pixel neighbours for each slice
			FinMax = "Final-max";
			Ext.CLIJ2_create3D(FinMax, OriWidth, OriHeight, nSlices, OriBit);
			for (SliceToProcess = 0; SliceToProcess < SlicePerFrame; SliceToProcess++) {
				Ext.CLIJ2_copySlice(SobelStack, SobelTemp, SliceToProcess);
				Ext.CLIJ2_maximum2DSphere(SobelTemp, MaxTemp, UserMaxRadius, UserMaxRadius);
				Ext.CLIJ2_copySlice(MaxTemp, FinMax, SliceToProcess);
				Ext.CLIJ2_release(SobelTemp);
				Ext.CLIJ2_release(MaxTemp);
			}
			Ext.CLIJ2_release(SobelStack);
				
			// z position of maximum z projection
			zPosMax = "z_position_of_maximum_z_projection";
			Ext.CLIJ2_zPositionOfMaximumZProjection(FinMax, zPosMax);
			Ext.CLIJ2_release(FinMax);
			
			//initialise z position of maximum z projection separation
			Ext.CLIJ2_create3D(zPosMaxSep, OriWidth, OriHeight, SlicePerFrame, OriBit);
			Ext.CLIJ2_threshold(zPosMax, threshTemp, SlicePerFrame-1);
			Ext.CLIJ2_copySlice(threshTemp, zPosMaxSep, SlicePerFrame-1);
			Ext.CLIJ2_release(threshTemp);
			
			//separate z position of maximum z projection in different slices
			for (SliceToProcess = SlicePerFrame-2; SliceToProcess > -1; SliceToProcess--) {
				threshold = SliceToProcess;
				Ext.CLIJ2_threshold(zPosMax, threshTemp0, threshold+1);
				Ext.CLIJ2_threshold(zPosMax, threshTemp, threshold);
				Ext.CLIJ2_subtractImages(threshTemp, threshTemp0, threshTempSub);
				Ext.CLIJ2_copySlice(threshTempSub, zPosMaxSep, SliceToProcess);
				Ext.CLIJ2_release(threshTempSub);
				Ext.CLIJ2_release(threshTemp0);
				Ext.CLIJ2_release(threshTemp);
			}
			
			//Convert to flaot
			Ext.CLIJ2_convertFloat(zPosMaxSep, zPosMaxSepFloat);
			Ext.CLIJ2_release(zPosMaxSep);
			
			//Gaussian blur
			Ext.CLIJ2_gaussianBlur2D(zPosMaxSepFloat, HeightMapBlur, UserBlurSigma, UserBlurSigma);
			Ext.CLIJ2_release(zPosMaxSepFloat);
			
			//Stack Map
			Ext.CLIJ2_multiplyImages(HeightMapBlur, UnFocusFrame, StackMap);
			Ext.CLIJ2_release(HeightMapBlur);
			
			//Sum slice
			Fin = "EDF-on-GPU";
			Ext.CLIJ2_sumZProjection(StackMap, Fin);
			Ext.CLIJ2_pull(Fin);
			//_________________________________
			run(OriBit+"-bit");
			Ext.CLIJ2_release(UnFocusFrame);
			//Store the focused image in the final stack image
			run("Concatenate...", "  title=Final open image1=Final image2=EDF-on-GPU image3=[-- None --]");	
			selectWindow("UnFocusFrame");
			close();
			Ext.CLIJ2_clear();
		}
		
		//clean the final stack image
		run("Duplicate...", "duplicate range=2-"+TotalTimeFrame+1);
		//some cleaning
		selectWindow("Final");
		close();
		selectWindow("Final-1");
		rename("Final");
		setBatchMode("exit and display");
		
		//save
		setVoxelSize(VxWidth, VxHeight, Vxdepth, Vxunit);
        saveAs("Tiff", outDir + File.separator + RootImageName+"_EDF-RW-GPU.tif");
        run("Close All");
	}
}
showMessage("Tada!");
```
