## Supplementary file S2-8. ImageJ macro used for image analysis for "RoPod, a customizable toolkit for non-invasive root imaging, reveals cell type-specific dynamics of plant autophagy": Ropod 2022_Supplementary file S6.ijm

```
/*
 * AIM OF THE MACRO
 * For all the files contained in the input folder, run this process for each root
 *  1- Concatenate the same XY position of two different time frame acquisition (repeat it for the the other XY position of the same root)
 *  2- correct the background
 *  3- stitch the 2 XY positions of the same root
 *  4- register the stitch image
 *  5- save the image in an output folder
 *  
 *  BEFORE RUNNING THE MACRO
 *  change the lines labeled with "*****"
 *  
 *  CREDIT
 *  Marjorie Guichard - Institut for Cell and Interaction Biology (Pr. Dr. Guido Grossmann lab) - Heinrich Heine Universität - Düsseldorf
 *  marjorie.guicharg at hhu.de
 *  29/03/2022
 */


inDir = getDir("select INPUT directory");
outDir = getDir("select OUTPUT directory");
filelist = getFileList(inDir);

//save the input folder as a text into the output folder
print("\\Clear");
print(inDir);
selectWindow("Log");
saveAs("Text", outDir + File.separator + "Input directory.txt");

//define list of tif file in folder
tifFileList = newArray();
for (i = 0; i < lengthOf(filelist); i++) {
    if (endsWith(filelist[i], ".tif")) { 
        tifFileList = Array.concat(tifFileList,filelist[i]);
    } 
}

NbOfFilePerGroup = 4;//**** change that if necessary
NbOfSubgroup = 2; //**** change that if necessary
NbOfFilePerSubGroup = 2; //**** change that if necessary
nbOfTifFiles = lengthOf(tifFileList); //**** change that if necessary
NbOfGroupToProcess = nbOfTifFiles/NbOfFilePerGroup; //**** change that if necessary

//process each group of files
for (GroupToProcess = 0; GroupToProcess < NbOfGroupToProcess; GroupToProcess++) {
	//define and open the images to process
	File1NbToOpen = GroupToProcess*NbOfFilePerSubGroup;
	File2NbToOpen = nbOfTifFiles/NbOfSubgroup + GroupToProcess*NbOfFilePerSubGroup;
	File3NbToOpen = GroupToProcess*NbOfFilePerSubGroup+1;
	File4NbToOpen = nbOfTifFiles/NbOfSubgroup + GroupToProcess*NbOfFilePerSubGroup+1;
	open(inDir + File.separator+tifFileList[File1NbToOpen]);
	
	//get Image information
	getVoxelSize(Vxwidth, Vxheight, Vxdepth, Vxunit);
	oriTitle = getTitle();
	GroupDescription = substring(oriTitle, 46, 52);//*****change the numbers to set the beginning and end character position of the part of the image name that should be kept
	rootGroupName = "20220322_Col_sucrose_Vert_RoPod5_"+GroupDescription;//****change that also
	
	//Proceed the opening
	rename("01-A");
	open(inDir + File.separator+tifFileList[File2NbToOpen]);
	rename("01-B");
	open(inDir + File.separator+tifFileList[File3NbToOpen]);
	rename("02-A");
	open(inDir + File.separator+tifFileList[File4NbToOpen]);
	rename("02-B");	
	
	//Concatenate the two same XY position 1 from different records and correct the background
	run("Concatenate...", "open image1=01-A image2=01-B");
	rename("01");
	run("Subtract Background...", "rolling=200 light sliding stack"); //****change the parameters if necessary
	
	//Concatenate the two same XY position 2 from different records and correct the background
	run("Concatenate...", "open image1=02-A image2=02-B");
	rename("02");
	run("Subtract Background...", "rolling=200 light sliding stack"); //****change the parameters if necessary
	
	//Stitch the 2 XY positions
	run("Pairwise stitching", "first_image=01 second_image=02 fusion_method=[Linear Blending] fused_image=01<->02 check_peaks=5 compute_overlap x=0.0000 y=0.0000 registration_channel_image_1=[Average all channels] registration_channel_image_2=[Average all channels] time-lapse_registration=[Register images adjacently over time] computation=[Save computation time (but use more RAM)] max/avg=2.50 absolute=3.50");
	run("8-bit");
	run("Grays");	
	rename("Stitch");
	
	//Fill the stitching background
	setBatchMode(true);
	for (FrameToProcess = 1; FrameToProcess < nSlices+1; FrameToProcess++) {
		setSlice(FrameToProcess);
		setThreshold(0, 1);
		run("Create Selection");
		setForegroundColor(250, 250, 250);
		run("Fill", "slice");	
		run("Select None");
	}
	setBatchMode(false);
	
	//some cleaning
	selectWindow("01");
	close();
	selectWindow("02");
	close();
	
	//Register the stitched image
	selectWindow("Stitch");
	run("Descriptor-based series registration (2d/3d + t)", "Stitch brightness_of=[Advanced ...] approximate_size=[Advanced ...] type_of_detections=[Minima only] subpixel_localization=[3-dimensional quadratic fit] transformation_model=[Rigid (2d)] number_of_neighbors=3 redundancy=1 significance=3 allowed_error_for_ransac=5 global_optimization=[All-to-all matching with range ('reasonable' global optimization)] range=5 choose_registration_channel=1 image=[Fuse and display] interpolation=[Linear Interpolation] detection_sigma=6.8375 threshold=0.0300"); //**** the sigma and threshold might need to be changed run this plugin separatly in "interactive" mode to define what value to use
	
	//Fill the registration background
	setBatchMode(true);
	for (FrameToProcess = 1; FrameToProcess < nSlices+1; FrameToProcess++) {
		setSlice(FrameToProcess);
		setThreshold(0, 1);
		run("Create Selection");
		setForegroundColor(250, 250, 250);
		run("Fill", "slice");	
		run("Select None");
	}
	setBatchMode(false);
	
	//set back the scale
	setVoxelSize(Vxwidth, Vxheight, Vxdepth, Vxunit);
	
	//save the file
	save(outDir + File.separator+rootGroupName+"_EDF-RW-GPU_Bckg-Stitch-reg.tif");
	run("Close All");
}

//The end
showMessage("Tada!");
```
