## Supplementary file S2-8. ImageJ macro used for image analysis for "RoPod, a customizable toolkit for non-invasive root imaging, reveals cell type-specific dynamics of plant autophagy": Ropod 2022_Supplementary file S7.ijm

```
/*
 * AIM OF THE MACRO
 * 	- Preselect the root hairs that will be analysed later using a straight line (Manual step)
 * 	- Save all the ROI in one file that will be used for a further process
 * 	
 * BEFORE RUNNING THE MACRO
 *	Open: a file with growing root hairs (e.g. a stiched image of a root)
 *	
 *CREDIT
 * Marjorie Guichard, Grossmann lab, Center for Organismal Studies, Heidelberg, Germany, 28/04/2022
 *
 */


//some cleaning
roiManager("Reset");

//get Image name and clean it
ImageName = getTitle();
dotIndex = indexOf(ImageName, ".tif");
RootImageName = substring(ImageName, 0, dotIndex);

//choose output directory
output = getDirectory("Choose output directory");

//Draw line over an 1st hair and store it in ROI manager
setTool("line");
setSlice(nSlices);
roiManager("show all without labels");
waitForUser("Draw a STRAIGHT line over a growing Root hair \n The line should start at the rootward end, \n and finish a bit further than the root hair tip \n Pay attention to draw the line at least after the slice when hair stop growing!!!!! \n (if you have doubt, draw on the last slice) \n save the line in ROI Manager (shortcut = t)");

//rename the ROIs
ROItoSelect = newArray();
for (ROItoProcess = 0; ROItoProcess < roiManager("count"); ROItoProcess++) {
	roiManager("select", ROItoProcess);
	if(ROItoProcess<9){
		roiManager("rename", "roi_0"+ROItoProcess+1);
	}
	else{
		roiManager("rename", "roi_"+ROItoProcess+1);
	}	
	// step useful for the further saving step
	ROItoSelect = Array.concat(ROItoSelect, ROItoProcess);
 }

//ROI manager saving
roiManager("select", ROItoSelect);
roiManager("save selected", output + File.separator + RootImageName+"_ROI.zip");

//some cleaning
run("Close All");
roiManager("deselect");
roiManager("delete");

showMessage("Tada!");
```
