## Supplementary file S2-8. ImageJ macro used for image analysis for "RoPod, a customizable toolkit for non-invasive root imaging, reveals cell type-specific dynamics of plant autophagy": Ropod 2022_Supplementary file S8.ijm

```
/* AIM OF THE MACRO
 * 	- This macro should be run on time lapse acquisitions where Regions Of Interest (ROI) have been drawn over root hair(s) (RH) of interest on the last time frame image
 * 	If there are several ROIs, this macro will loop to successively analyse thoose ROIs
 * 	1- Straighten the RHs of a bright field (BF) time lapse images using the ROIs defined before as guide
 * 	2- Manually define the RH medium line RHML
 * 		Use the RHML to mask the rest of the image so only the wanted hair is kept
 * 	3- Use the BF time lapse as a guide to track the RH tip over time
 * 		principe of the step:
 * 			2-1- threshold the BF image with MinError thresholding method
 * 			2-2- concider the white pixel that is at the more right side as the RH tip more each time frame
 * 			2-3- let the user manually check the automatic thresholding 	
 *	4- Projection of the RH tip tracking on the RHML (might be usefull to corect RH drifting at the begining of the growth ?) 
 *	5- Define the hair skeleton before and after the begining of the acquisition based on the RH tip tracking (not useful for only BF image, but usefull if there is also a Fluorescent image to analyse)
 *		4-1- If the hair didn´t start to grow at the begining of the acquisition, the RH skeleton is null
 *			and only the RH tip is used to produce the skeleton on the RHML
 * 		4-2- If the hair start to grow before the begining of the acquisition,
 * 			only is kept the region from the RH basis until the first RH tip track position defined at step 3
 * 			then the RH tip is used to define the skeleton
 * 	5- Save the straigthen BF image
 * 	6- Save the automatically or manually generated tracks of the RH position or RH skeleton
 * 	7- Calculate the RH displacement over time and save the sum up table
 * 	-The steps labeled with "******" can be modified if some adjustement are needed (look for them with the research tool. Shortcut: "ctlr + F"). Other steps can be of course modified, but this will be more tricky ^^
 * 	
 * BEFORE RUNNING THE MACRO
 *	- Open the BF image of a root (stiched or not)
 *	- Create or import already drawn ROIs covering the RH to analyse (e.g. ROIs generated with the macro "Pre-root hair tracking_ROI drawing_20190902")
 *	
 *CREDIT
 * Marjorie Guichard, Institut for cell and interaction biology, Pr. Dr. Guido Grossmann lab, Heinrich Heine Universität Düsseldorf, Germany, 28/04/2022
 *
 */

ROIWidth = 82; // ****** recommended value : twelve times the width of one hair (in pixel)
FineROIWidth = 15; // ****** recommended value : four times the width of one hair (in pixel)

//Choose output directory
outdir = getDirectory("Choose OUTPUT Directory");

//
BFID = getImageID();

//recover original image name
OriImageName = getTitle();
dotIndex = indexOf(OriImageName, ".tif");
//dotIndex = indexOf(OriImageName, "F");
OriImageRootName = substring(OriImageName, 0, dotIndex);
Dialog.create("Which title do you want to give to your file?");
Dialog.addString("", OriImageRootName,100);
Dialog.show();
FinalImageRootName = Dialog.getString();

// recover initial values
getVoxelSize(VxWidth, VxHeight, depth, unit);

//simplify image name
selectImage(BFID);
rename("BF");

ROIManagerOriSize = roiManager("count");//for future loop definition

// Select the ROI where to start the analysis
selectImage(BFID);
waitForUser("select the ROI of a RH you want to start with");
StartROI = roiManager("index");

for (i = StartROI; i < ROIManagerOriSize; i++) {//loop to screen all the ROI

	//Create the arrays to fill the final table
	ImageNameFinTable = newArray();
	StartSliceFinTable = newArray();
	DriftEndSliceFinTable = newArray();	
	hairGrowthEndinTable = newArray();
	SliceNbFinTable = newArray();
	RHTipProjectXFinTable = newArray();
	RHTipProjectYFinTable = newArray();
	HairDisplacementOverTimeUnitFinTable = newArray();
	HairCumulDisplacementOverTimeUnitFinTable = newArray();
	RHTotalLengthUnitFinTable = newArray();
	RemarkAboutTheImageFinTable = newArray();
	
	// Get some string for future process
	roiManager("select", i);
	ROIName = Roi.getName;
	
	// Straightening a single hair - BF image
	selectWindow("BF");
	roiManager("select", i);
	BFImageID = getImageID();
	TotSlices = nSlices;	
	setBatchMode(true);
	for (SliceToStraighten = 0; SliceToStraighten <  TotSlices; SliceToStraighten++) {
	selectImage(BFImageID);
	setSlice(SliceToStraighten+1);
	run("Straighten...", "title=BF line="+ROIWidth+"");
	}
	run("Images to Stack", "name=BF-concat title=[] use");
	setBatchMode("exit and display");
	run("8-bit");
	rename("BF_straighten");
	
	// get information for further steps
	StraightImageWidth = getWidth();
	StraightImageHeight = getHeight();

	//Window organisation
	getLocationAndSize(SingleRHWindowX, SingleRHWindowY, SingleRHWindowW, SingleRHWindowH);
	WindowSizeZoomFactorW = (screenWidth/3)/SingleRHWindowW;
	setLocation(SingleRHWindowW * WindowSizeZoomFactorW, (screenHeight/2)-(SingleRHWindowH * WindowSizeZoomFactorW/2), SingleRHWindowW * WindowSizeZoomFactorW, SingleRHWindowH * SingleRHWindowH);
	
	//define the begining of the hair growth	
	setSlice(nSlices/2);	
	getLocationAndSize(screenX, screenY, screenW, screenH);
	Dialog.createNonBlocking("Information about the hair");
	Dialog.setLocation(screenX+screenW,screenY);
	Dialog.addNumber("At which slice the growth is starting?", 1);
	Dialog.addNumber("Approximately, when is the hair growth finishing?", TotSlices);
	Dialog.show();
	StartSlice = Dialog.getNumber();
	HairGrowthEnd = Dialog.getNumber();

	// Manual draw of the RH medium line RHML
	setSlice(HairGrowthEnd);
	setTool("polyline");
	getLocationAndSize(screenX, screenY, screenW, screenH);
	Dialog.createNonBlocking("");
	Dialog.setLocation(screenX+screenW,screenY);
	Dialog.addMessage("Draw the root hair medium line on the last frame of root hair growth", 15, "black");
	Dialog.show();
	getSelectionCoordinates(AllRHMediumLineXCustom, AllRHMediumLineYCustom);
	
	// Save the manually defined skeleton
	selectWindow("BF_straighten");	
	setSlice(nSlices);
	makeSelection("polyline", AllRHMediumLineXCustom, AllRHMediumLineYCustom);
	roiManager("add");
	roiManager("select", ROIManagerOriSize);
	roiManager("rename", ROIName + "_RHML_custom");
	
	// Threshold the root hair	
	selectWindow("BF_straighten");
	roiManager("show none");
	run("Select None");
	run("Duplicate...", "duplicate");
	rename("Masked_RH");
	run("Gaussian Blur...", "sigma=1 stack");
	setAutoThreshold("MinError no-reset");
	run("Convert to Mask", "method=MinError background=Light black");
	
	//removal of the dust
	run("Analyze Particles...", "size=0-500 pixel show=Masks stack");//**** change the maximum particle size if necessary
	imageCalculator("Subtract create stack", "Masked_RH","Mask of Masked_RH");
	rename("temp");
	selectWindow("Masked_RH");
	close();
	selectWindow("Mask of Masked_RH");
	close();
	selectWindow("temp");
	rename("Masked_RH");
	
	//delete the surrounding of the root hair
	roiManager("select", ROIManagerOriSize);
	roiManager("Set Line Width", FineROIWidth);
	run("Line to Area");
	run("Make Inverse");
	setForegroundColor(0, 0, 0);
	for (SliceToFill = 1; SliceToFill < nSlices+1; SliceToFill++) {
		setSlice(SliceToFill);
		fill();
	}
	run("Select None");
	
	// last automatic cleaning
	run("Options...", "iterations=2 count=1 black do=Erode stack"); //******* change iteration if necessary
	run("Options...", "iterations=2 count=1 black do=Dilate stack"); //******* change iteration if necessary
	setBatchMode("exit and display");
	
	//Window organisation
	getLocationAndSize(SingleRHWindowX, SingleRHWindowY, SingleRHWindowW, SingleRHWindowH);
	WindowSizeZoomFactorW = (screenWidth/3)/SingleRHWindowW;
	setLocation(SingleRHWindowW * WindowSizeZoomFactorW+10, (screenHeight/2)-(SingleRHWindowH * WindowSizeZoomFactorW/2)+10, SingleRHWindowW * WindowSizeZoomFactorW, SingleRHWindowH * SingleRHWindowH);
		
	//check for main dust
	setTool("rectangle");
	setSlice(HairGrowthEnd);
	waitForUser("draw a rectangle around the main dusts that might disturb hair tip tracking");
	if (selectionType() != -1 ) {
		for (Slice = 1; Slice <nSlices+1; Slice ++){
			setSlice(Slice);
			setColor("black");
			fill();
		}
		run("Select None");
	}
	
	// Draw a verticale line on the left side of the image so it is not totally black	
	makeRectangle(0, 0, 1, StraightImageHeight+1);
	for (Slice = 1; Slice <nSlices+1; Slice ++){
			setSlice(Slice);
			setColor("white");
			fill();
	}	
	roiManager("show none");
	run("Select None");
	
	//Screen all slices from right to left to find the first x position with a white pixel: this is the root hair tip!
	XToScreen = StraightImageWidth-1;
	StackRHTipX = newArray(); //will store the RH tip position in X for each slice
	StackRHTipY = newArray(); //will store the RH tip position in Y for each slice
	
	for (j = 1; j < nSlices+1; j++) {
	XStatue = 0; // = 0 if there is no white pixel in one group of same X coordinates. = 1 If there is there is a white pixel
	setSlice(j);
		while (XStatue==0) {
			YPossible = newArray(); //will store all the Y position of the pixels with value of 255
			for (k = 0; k < StraightImageHeight; k++) {
				PxValue = getPixel(XToScreen, k);
				if (PxValue == 255) {
					YPossible = Array.concat(YPossible, k);
				}else {}
			}
			if (lengthOf(YPossible)== 0) {//To do if there are no white pixels in whole X column
				XStatue == 0; // To be able to loop to the next X position
				XToScreen = XToScreen - 1; //define the next X position to screen
			}
			else {//to do if there are white pixels in the whole X column
				XStatue = 1;//to end the screening
				Array.getStatistics(YPossible, min, max, RHTipY, stdDev);
				RHTipX = XToScreen;
				XToScreen = StraightImageWidth-1;
			}
		}
		StackRHTipX = Array.concat(StackRHTipX, RHTipX);
		StackRHTipY = Array.concat(StackRHTipY, RHTipY);
	}

	//Some cleaning	
	roiManager("select", ROIManagerOriSize);
	roiManager("Set Line Width", 1);
	selectWindow("Masked_RH");
	close();
	
	//Screen organisation
	selectWindow("BF_straighten");
	getLocationAndSize(WindowX, WindowY, WindowWidth, WindowHeight);
	setLocation((screenWidth-WindowWidth)/2, (screenHeight-WindowHeight)/2);

	//Check step
	ROIToDelete = newArray(); // for future cleaning
	for (j = 0; j < lengthOf(StackRHTipX); j++) {
		setSlice(j+1);
		makePoint(StackRHTipX[j], StackRHTipY[j]);
		roiManager("add");
		roiManager("select", ROIManagerOriSize+1+j);
		roiManager("rename", j+1);
		Roi.setPosition(0,j+1,0); // step needed because the "add" tool of roi Manager isn't recording the slice position in macro mode...
		roiManager("Update"); // step needed because the "add" tool of roi Manager isn't recording the slice position in macro mode...
		ROIToDelete = Array.concat(ROIToDelete, ROIManagerOriSize+1+j);//for future cleaning
	}
	roiManager("Show All");
	setSlice(HairGrowthEnd);
	
	// for the checking process to not start from zero if there is a mistake
	StackRHTipX2= newArray();
	StackRHTipX2 = Array.concat(StackRHTipX2, StackRHTipX);
	StackRHTipY2= newArray();
	StackRHTipY2 = Array.concat(StackRHTipY2, StackRHTipY);
	
	//initiate checking loop
	Dialog.createNonBlocking("");
	Dialog.addMessage("From slice "+StartSlice+" to "+HairGrowthEnd+", are you happy with the tracking?", 20, "magenta");
	Dialog.addRadioButtonGroup("", newArray("Yes", "No"), 1, 2, "No");
	Dialog.show();
	if (Dialog.getRadioButton() == "yes") {
		IsUserHappy = true;
		IniIsUserHappy = true;
	}else {
		IsUserHappy = false;
		IniIsUserHappy = false;
	}
	
	//checking loop
	while (IsUserHappy==false) {
		//some cleaning
		roiManager("select", ROIToDelete);
		roiManager("delete");
		ROIToDelete = newArray(); // for future cleaning
		
		//create array where to store the custom RH tip selections
		StackRHTipXCustom = newArray();
		StackRHTipYCustom = newArray();
		showMessage("Then adjust it for each slice. Ready?");
		for (j = 0; j < lengthOf(StackRHTipX2); j++) {
			if (j+1<StartSlice || j+1>HairGrowthEnd ) {				
				setSlice(j+1);
				makePoint(StackRHTipX2[j], StackRHTipY2[j]);
				getSelectionCoordinates(xpoints, ypoints);
				StackRHTipXCustom = Array.concat(StackRHTipXCustom, xpoints);
				StackRHTipYCustom = Array.concat(StackRHTipYCustom, ypoints);
			}else {
				setTool("point");
				setSlice(j+1);
				makePoint(StackRHTipX2[j], StackRHTipY2[j]);
				waitForUser("Adjust the tip position in slice "+j+1);
				getSelectionCoordinates(xpoints, ypoints);
				StackRHTipXCustom = Array.concat(StackRHTipXCustom, xpoints);
				StackRHTipYCustom = Array.concat(StackRHTipYCustom, ypoints);
			}
		}
		//prepare the last checking
		for (j = 0; j < lengthOf(StackRHTipX2); j++) {
			setSlice(j+1);
			makePoint(StackRHTipXCustom[j], StackRHTipYCustom[j]);
			roiManager("add");
			roiManager("select", ROIManagerOriSize+1+j);
			roiManager("rename", j+1);
			Roi.setPosition(0,j+1,0); // step needed because the "add" tool of roi Manager isn't recording the slice position in macro mode...
			roiManager("Update"); // step needed because the "add" tool of roi Manager isn't recording the slice position in macro mode...
			ROIToDelete = Array.concat(ROIToDelete, ROIManagerOriSize+1+j);//for future cleaning
		}
		// last checking
		roiManager("Show All");
		setSlice(HairGrowthEnd);
		Dialog.createNonBlocking("");
		Dialog.addMessage("From slice "+StartSlice+" to "+HairGrowthEnd+", are you happy with the tracking?", 20, "magenta");
		Dialog.addRadioButtonGroup("", newArray("yes", "No"), 1, 2, "yes");
		Dialog.show();
		if (Dialog.getRadioButton() == "yes") {
			IsUserHappy = true;
		}else {
			IsUserHappy = false;
		}
	
		//for the checking process to not start from zero if there is a mistake
		StackRHTipX2= newArray();
		StackRHTipX2 = Array.concat(StackRHTipX2, StackRHTipXCustom);
		StackRHTipY2= newArray();
		StackRHTipY2 = Array.concat(StackRHTipY2, StackRHTipYCustom);
	}//end of RH tip tracking checking loop
	
	//some cleaning
	roiManager("select", ROIToDelete);
	roiManager("delete");
	
	//Store the automatic RH selection
	roiManager("show none");
	run("Select None");
	makeSelection("multipoint",StackRHTipX, StackRHTipY);
	roiManager("add");
	roiManager("select", ROIManagerOriSize+1);
	roiManager("rename", ROIName + "_Root hair tip track_auto");
	
	//Store the custom RH selection if user happy with the automatic tracking and implement the manually labeled drifting hairs
	if(IniIsUserHappy==true){
		StackRHTipXCustom = StackRHTipX;
		StackRHTipYCustom = StackRHTipY;
		for (SliceToProcess = 0; SliceToProcess < StartSlice; SliceToProcess++) {// set the hair position before beginning of hair growth
			StackRHTipXCustom[SliceToProcess] = StackRHTipXCustom[StartSlice-1];
			StackRHTipYCustom[SliceToProcess] = StackRHTipYCustom[StartSlice-1];			
		}
		for (SliceToProcess = HairGrowthEnd-1; SliceToProcess < TotSlices; SliceToProcess++) {// set the hair position after end of hair growth
			StackRHTipXCustom[SliceToProcess] = StackRHTipXCustom[HairGrowthEnd-1];
			StackRHTipYCustom[SliceToProcess] = StackRHTipYCustom[HairGrowthEnd-1];
		}
	}

	else {

		for (SliceToProcess = 0; SliceToProcess < StartSlice; SliceToProcess++) {// set the hair position before beginning of hair growth
			StackRHTipXCustom[SliceToProcess] = StackRHTipXCustom[StartSlice-1];
			StackRHTipYCustom[SliceToProcess] = StackRHTipYCustom[StartSlice-1];			
		}
		for (SliceToProcess = HairGrowthEnd-1; SliceToProcess < TotSlices; SliceToProcess++) {// set the hair position after end of hair growth
			StackRHTipXCustom[SliceToProcess] = StackRHTipXCustom[HairGrowthEnd-1];
			StackRHTipYCustom[SliceToProcess] = StackRHTipYCustom[HairGrowthEnd-1];
		}
	}
	roiManager("show none");
	run("Select None");
	makeSelection("multipoint",StackRHTipXCustom, StackRHTipYCustom);
	roiManager("add");
	roiManager("select", ROIManagerOriSize+2);
	roiManager("rename", ROIName + "_Root hair tip track_custom");

	//Get RH tip projection on RH final medium line (RHFML) for each time point
	//method: analyse triangle ABC formed by RH position (A), and one segment of RH Medium Line (BC)
	// (B): segment side closer from RH basis ; (C): segment side farther from RH basis
	// AH = heigh of the triangle (emerging from A)
	// (H) = instersection between AH and BC
	// AI = median of the triangle (emerging from A)
	// (I) = instersection between AI and BC
	
	// initiation parameters
	HXPointAllRH = newArray();
	HYPointAllRH = newArray();

	//Loop to project the different RH tip positions
	for (RHTip = 0; RHTip < lengthOf(StackRHTipXCustom); RHTip++) {
		RHMedianeFromRML = newArray();
		//Find from which RHFML segment the RH tip is closer (look for AI minimum length)
		for (c=0; c<lengthOf(AllRHMediumLineXCustom)-1; c++)
		{
			ABLength = length( StackRHTipXCustom[RHTip], StackRHTipYCustom[RHTip], AllRHMediumLineXCustom[c], AllRHMediumLineYCustom[c] );
			ACLength = length( StackRHTipXCustom[RHTip], StackRHTipYCustom[RHTip], AllRHMediumLineXCustom[c+1], AllRHMediumLineYCustom[c+1] );
			BCLength = length( AllRHMediumLineXCustom[c], AllRHMediumLineYCustom[c], AllRHMediumLineXCustom[c+1], AllRHMediumLineYCustom[c+1] );
			AILength = AI( ABLength, ACLength, BCLength );
			RHMedianeFromRML = Array.concat(RHMedianeFromRML, AILength);
		}
		RHMedianeFromRMLRanked = Array.rankPositions(RHMedianeFromRML);
		RHMedianeFromRMLMinRank = RHMedianeFromRMLRanked[0];
	
		//Find H coordinate on the closer RML segment
		HXPoint = HCoorX( StackRHTipXCustom[RHTip], StackRHTipYCustom[RHTip], AllRHMediumLineXCustom[RHMedianeFromRMLMinRank], AllRHMediumLineYCustom[RHMedianeFromRMLMinRank], AllRHMediumLineXCustom[RHMedianeFromRMLMinRank+1], AllRHMediumLineYCustom[RHMedianeFromRMLMinRank+1]);
		HXPointAllRH = Array.concat(HXPointAllRH, HXPoint);
		HYPoint = HCoorY( StackRHTipXCustom[RHTip], StackRHTipYCustom[RHTip], AllRHMediumLineXCustom[RHMedianeFromRMLMinRank], AllRHMediumLineYCustom[RHMedianeFromRMLMinRank], AllRHMediumLineXCustom[RHMedianeFromRMLMinRank+1], AllRHMediumLineYCustom[RHMedianeFromRMLMinRank+1]);
		HYPointAllRH = Array.concat(HYPointAllRH, HYPoint);
	}
	//
	roiManager("show none");
	run("Select None");
	makeSelection("polyline", HXPointAllRH, HYPointAllRH);
	roiManager("add");
	roiManager("select", ROIManagerOriSize+3);
	roiManager("rename", ROIName + "_RHML_custom_RH tip projection");

	//define the RH medium line at the beginning of the acquisition
	AllRHXCustomSlice1 = newArray();
	AllRHYCustomSlice1 = newArray();
	j = 0;
	if(StartSlice==1){
		while (AllRHMediumLineXCustom[j]+10 < HXPointAllRH[0]) {
			AllRHXCustomSlice1 = Array.concat(AllRHXCustomSlice1, AllRHMediumLineXCustom[j]);
			AllRHYCustomSlice1 = Array.concat(AllRHYCustomSlice1, AllRHMediumLineYCustom[j]);
			j = j+1;
		}
		// finish the RH medium line at the beginning of the acquisition
		AllRHXCustomSlice1 = Array.concat(AllRHXCustomSlice1, HXPointAllRH[0]);
		AllRHYCustomSlice1 = Array.concat(AllRHYCustomSlice1, HYPointAllRH[0]);
		//Make selection of RHML before beginning of RH growth or recording
		roiManager("show none");
		run("Select None");
		makeSelection("polyline", AllRHXCustomSlice1, AllRHYCustomSlice1);
	}else{
		AllRHXCustomSlice1 = Array.concat(AllRHXCustomSlice1, AllRHMediumLineXCustom[0]);
		AllRHYCustomSlice1 = Array.concat(AllRHYCustomSlice1, AllRHMediumLineYCustom[0]);
		//Make selection of RHML before beginning of RH growth or recording
		roiManager("show none");
		run("Select None");
		makeSelection("point", AllRHXCustomSlice1, AllRHYCustomSlice1);
	}
	roiManager("add");
	roiManager("select", ROIManagerOriSize+4);
	roiManager("rename", ROIName + "_RHML_custom_before_growth_start");
	
	//Make polyline of RHML after beginning of acquisition
	HXPointAllRHAfterStart = newArray();
	HYPointAllRHAfterStart = newArray();
	if(StartSlice>1){
		HXPointAllRHAfterStart = Array.concat(HXPointAllRHAfterStart,HXPointAllRH);
		HYPointAllRHAfterStart = Array.concat(HYPointAllRHAfterStart,HYPointAllRH);
		for (Slice = 1; Slice < StartSlice+1; Slice++) {
			HXPointAllRHAfterStart[Slice-1] = AllRHMediumLineXCustom[0];
			HYPointAllRHAfterStart[Slice-1] = AllRHMediumLineYCustom[0];
		}
	}else{
		HXPointAllRHAfterStart = Array.concat(HXPointAllRHAfterStart,HXPointAllRH);
		HYPointAllRHAfterStart = Array.concat(HYPointAllRHAfterStart,HYPointAllRH);
	}
	roiManager("show none");
	run("Select None");
	makeSelection("polyline", HXPointAllRHAfterStart, HYPointAllRHAfterStart);
	roiManager("add");
	roiManager("select", ROIManagerOriSize+5);
	roiManager("rename", ROIName + "_RHML_custom_after_growth_start");
		
	//Saving and cleaning
	selectWindow("BF_straighten");
	save(outdir+File.separator+FinalImageRootName+"_"+ROIName+"_Straigthen-BF.tif");
	run("Select None");
	close();
	
	//Define ROI to save
	ROIManagerFinSize = roiManager("count");
	ROIToSave = newArray();
	for (j = ROIManagerOriSize; j < ROIManagerFinSize; j++) {
		ROIToSave = Array.concat(ROIToSave, j);
	}
	roiManager("select", ROIToSave);
	roiManager("save selected", outdir+File.separator+FinalImageRootName+"_"+ROIName+"_RH-tip-track.zip");
	//some cleaning
	roiManager("delete");
	
	//ask the user if there is a remark to do on this image
	Dialog.create("General remarks");
	Dialog.addMessage("If you have a remark to do about this image, write it there");
	Dialog.addString("R", "None",100);
	Dialog.show();
	ImageRemark = Dialog.getString();

	//calculate the length of the hair at the beginning of the record if the hair already grew
	RHTotalLengthIniUnit = LengthOfSelection(AllRHXCustomSlice1, AllRHYCustomSlice1)*VxWidth;
	//set the initial values of the table
	ImageNameFinTable = Array.concat(ImageNameFinTable, FinalImageRootName+"_"+ROIName+"_Straigthen-BF.tif");
	StartSliceFinTable = Array.concat(StartSliceFinTable , StartSlice);
	hairGrowthEndinTable = Array.concat(hairGrowthEndinTable , HairGrowthEnd);
	SliceNbFinTable = Array.concat(SliceNbFinTable, 1);
	RHTipProjectXFinTable = Array.concat(RHTipProjectXFinTable, HXPointAllRH[0]);
	RHTipProjectYFinTable = Array.concat(RHTipProjectYFinTable, HYPointAllRH[0]);
	HairDisplacementOverTimeUnitFinTable = Array.concat(HairDisplacementOverTimeUnitFinTable, 0);
	HairCumulDisplacementOverTimeUnitFinTable = Array.concat(HairCumulDisplacementOverTimeUnitFinTable,0);
	RHTotalLengthUnitFinTable = Array.concat(RHTotalLengthUnitFinTable, RHTotalLengthIniUnit);
	RemarkAboutTheImageFinTable = Array.concat(RemarkAboutTheImageFinTable, ImageRemark);
	
	// Measure the displacement of the hair over time
	//loop to fill the measurement table
	for (j = 0; j < lengthOf(HXPointAllRH)-1; j++) {		
		SegmentBetweenTwoTimePointsX = newArray(HXPointAllRHAfterStart[j],HXPointAllRHAfterStart[j+1]);
		SegmentBetweenTwoTimePointsY = newArray(HYPointAllRHAfterStart[j],HYPointAllRHAfterStart[j+1]);
		HairDisplacementInSliceUnit = LengthOfSelection(SegmentBetweenTwoTimePointsX, SegmentBetweenTwoTimePointsY)*VxWidth;
		ImageNameFinTable = Array.concat(ImageNameFinTable, FinalImageRootName+"_"+ROIName+"_Straigthen-BF.tif");
		StartSliceFinTable = Array.concat(StartSliceFinTable , StartSlice);		
		hairGrowthEndinTable = Array.concat(hairGrowthEndinTable , HairGrowthEnd);
		SliceNbFinTable = Array.concat(SliceNbFinTable, j+2);		
		RHTipProjectXFinTable = Array.concat(RHTipProjectXFinTable, HXPointAllRH[j+1]);
		RHTipProjectYFinTable = Array.concat(RHTipProjectYFinTable, HYPointAllRH[j+1]);
		HairDisplacementOverTimeUnitFinTable = Array.concat(HairDisplacementOverTimeUnitFinTable, HairDisplacementInSliceUnit);
		HairCumulDisplacementOverTimeUnitFinTable = Array.concat(HairCumulDisplacementOverTimeUnitFinTable, HairCumulDisplacementOverTimeUnitFinTable[j]+HairDisplacementOverTimeUnitFinTable[j+1]);
		RHTotalLengthUnitFinTable = Array.concat(RHTotalLengthUnitFinTable, RHTotalLengthUnitFinTable[j]+HairDisplacementOverTimeUnitFinTable[j+1]);
		RemarkAboutTheImageFinTable = Array.concat(RemarkAboutTheImageFinTable, ImageRemark);
	}
	
	//Set and save the result table
	Table.create("Results summary");
	Table.setColumn("Image_name", ImageNameFinTable);
	Table.setColumn("Slice_when_RH_Start_growing",StartSliceFinTable);
	Table.setColumn("Approximal_Slice_when_RH_Stop_growing",hairGrowthEndinTable);
	Table.setColumn("Slice_number", SliceNbFinTable);
	Table.setColumn("Hair_Tip_projected_X_position", RHTipProjectXFinTable);
	Table.setColumn("Hair_Tip_projected_Y_position", RHTipProjectYFinTable);
	Table.setColumn("Hair_displacement_(micron)", HairDisplacementOverTimeUnitFinTable);
	Table.setColumn("Hair_cumulative_displacement_(micron)", HairCumulDisplacementOverTimeUnitFinTable);
	Table.setColumn("Hair_Length_(micron)", RHTotalLengthUnitFinTable);
	Table.setColumn("Image_remark", RemarkAboutTheImageFinTable);
	Table.save(outdir+File.separator+FinalImageRootName+"_"+ROIName+"_Hair-length.xls");
	Table.reset("Results summary");
	
	showMessage("end of "+ROIName+" processing");
}//end of loop to screen the ROI


showMessage("Tada!"); // The end

// ************************ FUNCTIONS DEFINITION ************************
// ______1st SET OF FUNCTIONS (FOR hair length measurement)______
function LengthOfSelection(SelectionArrayX, SelectionArrayY){
	AB=0;
	for (a = 0; a < lengthOf(SelectionArrayX)-1; a++) {
		ABSegment = sqrt(pow(SelectionArrayX[a]-SelectionArrayX[a+1],2)+pow(SelectionArrayY[a]-SelectionArrayY[a+1],2));
		AB = AB + ABSegment;
	}
	return AB;
}

// ______2ND SET OF FUNCTIONS (To make a projection of the RH tip on the final RH medium line)______
//Function to return length between 2 points with cartesian coordinates
function length( x_0, y_0, x_1, y_1 )
	{
		return sqrt( pow( x_0 - x_1, 2 ) + pow( y_0 - y_1, 2 ) );
	}

//Function to return length of median extract from A of ABC triangle 
function AI( AB, AC, BC )
	{
		return sqrt((pow( AB, 2 )+pow( AC, 2 )-(0.5*pow( BC, 2 )))/2);
	}

//Functions to return H coordinate in ABC triangle
function HCoorX( Xa, Ya, Xb, Yb, Xc, Yc)
	{
		D = Xa*(Xc-Xb)+Ya*(Yc-Yb);
		E = Xc-Xb;
		F = Yc-Yb;
		G = (Yc-Yb)/(Xc-Xb);
		H = Yb - Xb*((Yc-Yb)/(Xc-Xb));
		return (-D + (F*H))/(-E-(G*F));
	}
	
function HCoorY( Xa, Ya, Xb, Yb, Xc, Yc)
	{
		D = Xa*(Xc-Xb)+Ya*(Yc-Yb);
		E = Xc-Xb;
		F = Yc-Yb;
		G = (Yc-Yb)/(Xc-Xb);
		H = Yb - Xb*((Yc-Yb)/(Xc-Xb));
		return (G*((-D+(F*H))/(-E-(G*F))))+H;
	}
```
