## Supplementary file S10. R scripts used for root hair growth analysis for "RoPod, a customizable toolkit for non-invasive root imaging, reveals cell type-specific dynamics of plant autophagy": 20220420_sucrose_treatment_Verto_RoPod5_analysis2_fit-and-growth-time_clean.R

```
IniHairTable <- read.csv("F:/Microscopy/Others/20220322_sucrose_treatment_Vertosc_RoPod5/6-analysis/20220322_Col_sucrose_Vert_RoPod5_EDF-RW-GPU_group.csv", sep=";")

# To organize / merge the table
# To use the tool needed to extract information from the table 
# To make plots
# To modify string shape in the graph
library(tidyverse)
library(dplyr)
library(ggplot2)

woRemark <- filter(IniHairTable, IniHairTable$Image_remark == "None")

# function needed for visualization purposes
sigmoid = function(params, x) {
  params[1] / (1 + exp((params[2]-x)/params[3]))
}

# function needed for visualization purposes in nested data
sigmoid_for_nested_data = function(params, x) {
  y_result = params[1] / (1 + exp((params[2]-x)/params[3]))
  return(y_result)
}

#function needed to create the groups for violin plots
Violon_group_function = function(Col_RH_Growth_end, Col_RH_Growth_start, Col_Treatment){
  if(Col_RH_Growth_end <= 4.66666666666666666666666666666667*60 & Col_Treatment == "1/2MS"){
    return("Ctrl_BT")
  }else if (Col_RH_Growth_end <= 4.66666666666666666666666666666667*60 & Col_Treatment == "1/2MS_sucrose"){
    return("Treat_BT")
  }else if (Col_RH_Growth_start > 4.66666666666666666666666666666667*60 &  Col_Treatment == "1/2MS"){
    return("Ctrl_AT")
  }else if (Col_RH_Growth_start > 4.66666666666666666666666666666667*60 &  Col_Treatment == "1/2MS_sucrose"){
    return("Treat_AT")
  }else {
    return(NA)
  }
}

Violon_group_factor_function = function(Col_RH_Growth_end, Col_RH_Growth_start, Col_Treatment){
  if(Col_RH_Growth_end <= 4.66666666666666666666666666666667*60 & Col_Treatment == "1/2MS"){
    return(1)
  }else if (Col_RH_Growth_end <= 4.66666666666666666666666666666667*60 & Col_Treatment == "1/2MS_sucrose"){
    return(3)
  }else if (Col_RH_Growth_start > 4.66666666666666666666666666666667*60 &  Col_Treatment == "1/2MS"){
    return(2)
  }else if (Col_RH_Growth_start > 4.66666666666666666666666666666667*60 &  Col_Treatment == "1/2MS_sucrose"){
    return(4)
  }else {
    return(NA)
  }
}


#Prepare the data, select the proper hairs and do the fit for all the remaining hairs
NestedHairForFit <- woRemark %>%
  group_by(woRemark$Root_ROI)%>%
  nest()%>%
  #shift the time column so the growth rate coloration fit with the slop of the future curve
  mutate(data = map(data,~.x %>%mutate('Time.min.shift'= lag(Time.min.))))%>%
  #Calculate the time relative to the begining of the treatment
  mutate(data = map(data,~.x %>%mutate('Time.hours.relative'= Time.hours.-4.66666666666666666666666666666667)))%>%
  #Calculate the time between 2 time frames
  mutate(data = map(data,~.x %>%mutate('Time_bw_2_frame'= Time.min.-Time.min.shift)))%>%
  #extract the information about RH growth start
  mutate('Slice_when_RH_Start_growing'=map(data, ~ .x$Slice_when_RH_Start_growing[1]))%>%
  #extract the information about RH growth end
  mutate('Approximal_Slice_when_RH_Stop_growing'=map(data, ~ .x$Approximal_Slice_when_RH_Stop_growing[1]))%>%
  #remove hairs that started to grow before the beginning of the acquisition
  filter(Slice_when_RH_Start_growing != 1)%>%
  #remove hairs that hasn't finished to grow before the end of the acquisition
  filter(Approximal_Slice_when_RH_Stop_growing != 88)%>%
  # Define fitting initial parameters
  # max hair length measured
  mutate('Lmax_ini'=map(data, ~ max(.x$Hair_cumulative_displacement_.micron.)))%>%
  # time when 1/2 of max hair length occurred
  mutate('d50_ini'=map2(.x = data, .y = Lmax_ini, ~ .x$Time.min.[which(abs(.x$Hair_cumulative_displacement_.micron.-(.y/2)) == min(abs(.x$Hair_cumulative_displacement_.micron.-(.y/2))))]))%>%
  # a slope defined between 1/2 of max hair length and and bit later
  mutate('delta_ini' = map(.x = d50_ini, ~0.1*.x))%>%
  #extract the information for fit in x
  mutate('x' = map(.x = data, ~ .x$Time.min.))%>%
  #extract the information for fit in y
  mutate('y' = map(.x = data, ~.x$Hair_cumulative_displacement_.micron.))%>%
  # do the fit
  mutate('fit_model' = pmap(
    list(
      xForFit = x,
      yForFit = y,
      Lmax_ini_ForFit = Lmax_ini,
      d50_ini_ForFit = d50_ini,
      delta_ini_ForFit = delta_ini),
    function(xForFit, yForFit, Lmax_ini_ForFit, d50_ini_ForFit, delta_ini_ForFit) nls(
      yForFit~Lmax/(1 + exp((d50 - xForFit)/delta)),
      start=list(Lmax=Lmax_ini_ForFit,d50=d50_ini_ForFit,delta=delta_ini_ForFit)
    )))%>%
  # get the final equation of the fit
  mutate('params' = map(.x = fit_model, ~coef(.x)))%>%
  # Define hair growth start and arrest
  mutate('RH_Growth_start' = map(.x = params, ~.x[2]-2.5*.x[3]))%>%
  # Define hair growth start and end
  mutate('RH_Growth_end' = map(.x = params, ~.x[2]+2.5*.x[3]))%>%
  # Calculate time difference
  mutate('RH_Growth_duration' = map2(.x = RH_Growth_start, .y = RH_Growth_end, ~.y-.x))%>%
  # Remove the name of the growth duration number
  mutate("RH_Growth_duration" = map(.x = RH_Growth_duration, ~as.numeric(.x["d50"])))%>%
  #Extract the treatment information
  mutate("Treatment" = map(.x = data, ~.x$Treatment[1]))%>%
  # generate the values for future plots of the sigmoid fits
  mutate(data = map2(.x = data, .y = params, ~.x %>%mutate("y_sigmoid" = sigmoid_for_nested_data(.y, .x$Time.min.))))%>%
  # create before / after groups
  mutate("Violon_groups" = pmap(
    list(
      Col_RH_Growth_end_n = RH_Growth_end,
      Col_RH_Growth_start_n = RH_Growth_start,
      Col_Treatment_n = Treatment
    ),
    function(Col_RH_Growth_end_n, Col_RH_Growth_start_n, Col_Treatment_n) Violon_group_function(Col_RH_Growth_end_n,Col_RH_Growth_start_n,Col_Treatment_n)
  ))

# plot the fit for all selected hairs
UnestedFit <- NestedHairForFit %>%
  # remove duplicated names for future unesting
  select(-c("Treatment","Slice_when_RH_Start_growing","Approximal_Slice_when_RH_Stop_growing"))%>%
  unnest(cols = data)%>%
  filter(Violon_groups !="NA")

UnestedFit <- UnestedFit[names(UnestedFit) !='woRemark$Root_ROI']
UnestedFit$Violon_groups <- as.character(UnestedFit$Violon_groups)
UnestedFit$Violon_groups <- factor(UnestedFit$Violon_groups, levels = c("Ctrl_BT","Ctrl_AT","Treat_BT","Treat_AT"))

Xaxis <- seq(-4,12,by = 2)
Yaxis <- seq(0,400, by = 50)
UnestedFitPlot <- UnestedFit %>% ggplot()+
  geom_point(mapping = aes(x = Time.hours.relative, y = Hair_cumulative_displacement_.micron. , group = Root_ROI, colour =Root_ROI))+
  geom_line(mapping = aes(x = Time.hours.relative, y = y_sigmoid , group = Root_ROI, colour =Root_ROI))+
  facet_wrap(vars(Violon_groups))+
  labs( x = "Time (hours)", y = "Hair length (µm)", colour = "Growth rate\n (µm/min)")+
  scale_x_continuous(breaks= Xaxis, labels = Xaxis, minor_breaks = NULL, limits = c(-5,8), expand = expansion(mult = c(0, .02)))+
  scale_y_continuous(breaks= Yaxis, labels = Yaxis, minor_breaks = NULL, limits = c(0,400), expand = expansion(mult = c(0.02, .02)))+
  theme(
    text = element_text(family = "GraphFont", face="plain", size=10, color = "black"),
    axis.text = element_text(family = "GraphFont", face="plain", size=10, color = "black"),
    panel.background = element_rect(fill = "White", colour = "black"),
    panel.grid.major = element_blank(),
    panel.grid.minor = element_blank(),
    legend.position = "none")
UnestedFitPlot

# plot the fit for all selected hairs in the different violon groups
Violon_groups_to_check <- "Treat_AT"
UnestedFit_Violon_groups_to_check <- UnestedFit %>%
  filter(Violon_groups == Violon_groups_to_check)

Xaxis <- seq(-4,12,by = 2)
Yaxis <- seq(0,400, by = 50)
UnestedFit_Violon_groups_to_check_plot <- UnestedFit_Violon_groups_to_check %>% ggplot()+
  geom_point(mapping = aes(x = Time.hours.relative, y = Hair_cumulative_displacement_.micron. , group = Root_ROI, colour =Root_ROI))+
  geom_line(mapping = aes(x = Time.hours.relative, y = y_sigmoid , group = Root_ROI, colour =Root_ROI))+
  facet_wrap(vars(Root))+
  labs( x = "Time (hours)", y = "Hair length (µm)", colour = "Growth rate\n (µm/min)", title = Violon_groups_to_check)+
  scale_x_continuous(breaks= Xaxis, labels = Xaxis, minor_breaks = NULL, limits = c(-5,8), expand = expansion(mult = c(0, .02)))+
  scale_y_continuous(breaks= Yaxis, labels = Yaxis, minor_breaks = NULL, limits = c(0,400), expand = expansion(mult = c(0.02, .02)))+
  theme(
    text = element_text(family = "GraphFont", face="plain", size=10, color = "black"),
    axis.text = element_text(family = "GraphFont", face="plain", size=10, color = "black"),
    panel.background = element_rect(fill = "White", colour = "black"),
    panel.grid.major = element_blank(),
    panel.grid.minor = element_blank(),
    legend.position = "none")+
  geom_vline(xintercept=0, linetype="dotted", color = "black", size=0.5)
UnestedFit_Violon_groups_to_check_plot


#group data by treatment types and before/after treatment types
ViolonGroupNestedHairForFit <- NestedHairForFit %>%
  mutate("Violon_groups" = pmap(
    list(
      Col_RH_Growth_end_n = RH_Growth_end,
      Col_RH_Growth_start_n = RH_Growth_start,
      Col_Treatment_n = Treatment
    ),
    function(Col_RH_Growth_end_n, Col_RH_Growth_start_n, Col_Treatment_n) Violon_group_function(Col_RH_Growth_end_n,Col_RH_Growth_start_n,Col_Treatment_n)
  ))%>%
  mutate("Violon_groups_factors" = pmap(
    list(
      Col_RH_Growth_end_n = RH_Growth_end,
      Col_RH_Growth_start_n = RH_Growth_start,
      Col_Treatment_n = Treatment
    ),
    function(Col_RH_Growth_end_n, Col_RH_Growth_start_n, Col_Treatment_n) Violon_group_factor_function(Col_RH_Growth_end_n,Col_RH_Growth_start_n,Col_Treatment_n)
  ))%>%
  #remove hairs labelled NA
  filter(Violon_groups != "NA")%>%
  # convert columns as vector
  mutate("Violon_groups" = map_chr(.x = Violon_groups, ~.x))%>%
  mutate("RH_Growth_duration" = map_dbl(.x = RH_Growth_duration, ~.x))%>%
  mutate("Violon_groups_factors" = map_dbl(.x = Violon_groups_factors, ~.x))

#calculate median of each violon groups
ViolonGroupMed <- ViolonGroupNestedHairForFit%>%
  group_by(Violon_groups)%>%
  nest()%>%
  mutate("Mediane" = map(.x = data, ~median(.x$RH_Growth_duration)))%>%
  mutate("Mediane" = map_dbl(.x = Mediane, ~.x))%>%
  #extract violon group factors
  mutate("Med_group_factors" = map(.x = data, ~.x$Violon_groups_factors[[1]]))%>%
  mutate("Med_group_factors" = map_dbl(.x = Med_group_factors, ~.x))
  
#___________________
#statistical tests
library(FSA) #install.packages("FSA")
library(rcompanion)

#do test
GrowthDurationStat <- pairwise.wilcox.test(x = ViolonGroupNestedHairForFit$RH_Growth_duration, g= ViolonGroupNestedHairForFit$Violon_groups_factors, p.adjust.method = "bonferroni", paired = FALSE)


#do post test
GrowthDurationStatForPostTest <- cbind("Violon_groups_factors" = ViolonGroupNestedHairForFit$Violon_groups_factors, "RH_Growth_duration" = ViolonGroupNestedHairForFit$RH_Growth_duration )%>%
  as.tibble()

GrowthDurationStatForPostTest$RH_Growth_duration <- as.double(GrowthDurationStatForPostTest$RH_Growth_duration)
GrowthDurationStatForPostTest$Violon_groups_factors <-   as.integer(GrowthDurationStatForPostTest$Violon_groups_factors)
  
GrowthDurationDunn <-  dunnTest(GrowthDurationStatForPostTest$RH_Growth_duration ~ GrowthDurationStatForPostTest$Violon_groups_factors, data = GrowthDurationStatForPostTest, method="bh")

#get statistical groups letters
PH_GrowthDurationDunn <- cldList(comparison = GrowthDurationDunn$res$Comparison,
                                     p.value = GrowthDurationDunn$res$P.adj,
                                     threshold = 0.01,
                                     remove.space = TRUE)
PH_GrowthDurationDunn_Letter <- PH_GrowthDurationDunn$Letter

# to set the position of the letters in the graph
GrowthDurationStatForPostTest_MAX <- aggregate(data = GrowthDurationStatForPostTest, RH_Growth_duration ~ Violon_groups_factors, max)
GrowthDurationStatForPostTest_MAX$"letter" <- PH_GrowthDurationDunn_Letter
colnames(GrowthDurationStatForPostTest_MAX) <- c("group","max","letter")

#____________________
#count number of hairs and roots used for the growth duration analysis
Population_used <- ViolonGroupNestedHairForFit %>%
  select(c(`woRemark$Root_ROI`, Violon_groups_factors))%>%
  group_by(Violon_groups_factors)%>%
  nest()%>%
  mutate("data" = map(.x = data, ~separate(data = .x, col = `woRemark$Root_ROI`, into = c("box", "root","hair"), sep = "_")))%>%
  mutate("data" = map(.x = data, ~as.data.frame(.x)))%>%
  mutate("Root_hairs_population" = map(.x = data, ~length(.x$hair)))%>%
  mutate("Roots_population" = map(.x = data, ~length(unique(.x$root))))%>%
  # change into a list of numbers without name
  mutate("Root_hairs_population" = as.numeric(Root_hairs_population))%>%
  mutate("Roots_population" = as.numeric(Roots_population))


#______________
#do violin plot
Yaxis <- seq(0,7, by = 1)
Growth_duration_plot <- 
  ggplot()+
  geom_violin(data = ViolonGroupNestedHairForFit, mapping = aes(x=Violon_groups_factors, y = RH_Growth_duration/60, fill = Violon_groups), size=0.75)+
  labs(data = ViolonGroupNestedHairForFit, x = "", y = "Growth duration (h)")+
  scale_fill_manual(values=c('springgreen4','palegreen','magenta','plum1'))+
  scale_y_continuous(breaks= Yaxis, labels = Yaxis, minor_breaks = NULL, limits = c(0,7))+
  geom_point(data = ViolonGroupMed, mapping = aes(x=Med_group_factors, y = Mediane/60)) +
  geom_text(data = GrowthDurationStatForPostTest_MAX, aes(x= group, y = max/60+0.5, label = letter), size = 9)+
  geom_text(data = Population_used, aes(x= Violon_groups_factors, y = rep(x = 0.5, times = 4), label = Roots_population), size = 5, colour = "red")+
  geom_text(data = Population_used, aes(x= Violon_groups_factors, y = rep(x = 0, times = 4), label = Root_hairs_population), size = 5, colour = "blue")+
  theme(
    text = element_text(family = "GraphFont", face="plain", size=20, color = "black"),
    axis.text = element_text(family = "GraphFont", face="plain", size=20, color = "black"),
    panel.background = element_rect(fill = "White", colour = "black"),
    panel.grid.major = element_blank(),
    panel.grid.minor = element_blank(),
    axis.text.x=element_blank(),
    legend.position = "none")
Growth_duration_plot

ggsave(Growth_duration_plot, device = "svg", filename = paste("growth-duration_",Sys.Date(),".svg"))
```
