## Supplementary file S10. R scripts used for root hair growth analysis for "RoPod, a customizable toolkit for non-invasive root imaging, reveals cell type-specific dynamics of plant autophagy": 20220422_sucrose_treatment_Verto_RoPod5_analysis1_clean.R

```
IniHairTable <- read.csv("F:/Microscopy/Others/20220322_sucrose_treatment_Vertosc_RoPod5/6-analysis/20220322_Col_sucrose_Vert_RoPod5_EDF-RW-GPU_group.csv", sep=";")

# To organize / merge the table
# To use the tool needed to extract information from the table 
# To make plots
# To modify string shape in the graph
library(tidyverse)
library(dplyr)
library(ggplot2)

woRemark <- filter(IniHairTable, IniHairTable$Image_remark == "None")

#_______________________________________
#analysis with growth rate calculation and time filter - at the beginning AND the end of hair growth
NestedHair <- woRemark %>% group_by(woRemark$Root_ROI)%>% nest()
NestedHair4 <- NestedHair %>%
  #shift the time column so the growth rate coloration fit with the slop of the future curve
  mutate('Time.min.shift'=map(data, ~ lag(.x$Time.min.)))%>%
  mutate('data'=map2(data,Time.min.shift,~mutate(.x,Time.min.shift = .y)))%>%
  select(`woRemark$Root_ROI`,data)%>%
  #Calculate the time relative to the begining of the treatment
  mutate('Time.hours.relative'=map(data, ~ .x$Time.hours.-4.66666666666666666666666666666667))%>%
  mutate('data'=map2(data,Time.hours.relative,~mutate(.x,Time.hours.relative = .y)))%>%
  select(`woRemark$Root_ROI`,data)%>%
  #Calculate the time between 2 time frames
  mutate('Time_bw_2_frame'=map(data, ~ .x$Time.min.-.x$Time.min.shift))%>%
  mutate('data'=map2(data,Time_bw_2_frame,~mutate(.x,Time_bw_2_frame = .y)))%>%
  select(`woRemark$Root_ROI`,data)%>%
  #calculate the growht rate
  mutate('RH_growth_rate_micron_per_min'=map(data, ~ .x$Hair_displacement_.micron./.x$Time_bw_2_frame))%>%
  mutate('data'=map2(data,RH_growth_rate_micron_per_min,~mutate(.x,RH_growth_rate_micron_per_min = .y)))%>%
  select(`woRemark$Root_ROI`,data)%>%
  #shift the growth rate column so the growth rate coloration fit with the slop of the future curve
  mutate('RH_growth_rate_micron_per_min_shift'=map(data, ~ lead(.x$RH_growth_rate_micron_per_min)))%>%
  mutate('data'=map2(data,RH_growth_rate_micron_per_min_shift,~mutate(.x,RH_growth_rate_micron_per_min_shift = .y)))%>%
  select(`woRemark$Root_ROI`,data)%>%
  #Find out at which global time RH start to grow
  mutate('Time_RH_start_growth'=map(data, ~ ifelse(.x$Slice_when_RH_Start_growing==.x$Slice_number,.x$Time.min., -1)))%>%
  mutate('data'=map2(data,Time_RH_start_growth,~mutate(.x,Time_RH_start_growth = .y)))%>%
  select(`woRemark$Root_ROI`,data)%>%
  # copy the time of RH growth start in each row
  mutate('data' = map(data,~mutate(.x,Time_RH_start_growth = max(.x$Time_RH_start_growth))))%>%
  # calculate time relative to hair growth start
  mutate('Time_relative_to_RH_start_growth_min' = map(data,~.x$Time.min.-.x$Time_RH_start_growth))%>%
  mutate('data'=map2(data,Time_relative_to_RH_start_growth_min,~mutate(.x,Time_relative_to_RH_start_growth_min = .y)))%>%
  select(`woRemark$Root_ROI`,data)

#create function that will find the hair growth during the 40 last minutes
growthEnd <- function(dataFrameToCompute, TimeDifColumn, DisplacementColumn, TimeOffset){
  dataFrameOut <- dataFrameToCompute
  for (RowToCompute in 1:(nrow(dataFrameToCompute))) {
    TimeColToPRocess <-as.double(which(colnames(dataFrameToCompute)==TimeDifColumn))
    DisplacementColToPRocess <- as.double(which(colnames(dataFrameToCompute)==DisplacementColumn))
    CumTimeInLoop <- 0
    WhileLoopIndice <- 0
    CumDisplacementInLoop <- 0
    while (CumTimeInLoop < TimeOffset) {
      if((RowToCompute - WhileLoopIndice-1)<=0){
        CumDisplacementInLoop <- NA
        CumTimeInLoop <- TimeOffset+1
      }else{
        CumTimeInLoop <- CumTimeInLoop + dataFrameToCompute[[RowToCompute - WhileLoopIndice, TimeColToPRocess]]
        CumDisplacementInLoop <- CumDisplacementInLoop + dataFrameToCompute[[RowToCompute - WhileLoopIndice, DisplacementColToPRocess]]
        WhileLoopIndice <- WhileLoopIndice+1
      }
    }
    dataFrameOut[RowToCompute,'Displacement_during_time_offset'] <- CumDisplacementInLoop
    a<-CumDisplacementInLoop
  }
  return(dataFrameOut)
}

#analysis with growth rate calculation and time filter - only at the end of hair growth
NestedHair5 <- NestedHair4
prgs = txtProgressBar(min = 0, max = nrow(NestedHair5), initial = 0, style = 3) 
for (SubTableToProcess in 1:nrow(NestedHair5)) {
  setTxtProgressBar(prgs,SubTableToProcess)
  temp <- NestedHair5$data[[SubTableToProcess]]
  NestedHair5$data[[SubTableToProcess]] <- growthEnd(temp, 'Time_bw_2_frame', 'Hair_displacement_.micron.', 40)
}

# unest table
UnNested4 <- unnest(NestedHair5, cols = data)
UnNested4 <- UnNested4[names(UnNested4) !='woRemark$Root_ROI']

# remove time before 50 min after hair growth start
UnNested4afterRHStart <- filter(UnNested4, UnNested4$Time_relative_to_RH_start_growth_min >=50)
# remove time after hair growth start end
UnNested4LinearGrowth <- filter(UnNested4afterRHStart, UnNested4afterRHStart$Displacement_during_time_offset >=20)

# separated control and treated conditions
controlFromUnNested4 <- filter(UnNested4LinearGrowth,UnNested4LinearGrowth$Treatment == "1/2MS")
treatmentFromUnNested4 <- filter(UnNested4LinearGrowth, UnNested4LinearGrowth$Treatment=="1/2MS_sucrose")

# Do graph for control condition
Xaxis <- seq(-4,12,by = 2)
Yaxis <- seq(0,400, by = 50)
Selected_root_ctrl <- "B1_R07"

SingleRootControlPlot <- controlFromUnNested4 %>%
  filter(Root==Selected_root_ctrl)%>%
  ggplot()+
  geom_line(mapping = aes(x=Time.hours.relative, y = Hair_Length_.micron., colour = RH_growth_rate_micron_per_min_shift, group = Root_ROI ), lineend = "round", size=2)+
  scale_size_manual(values = c(2,4))+
  scale_colour_gradientn(colours = c("blue","cyan","green","yellow","orange","red"), limits = c(0,2.1))+
  labs( x = "Time (hours)", y = "Hair length (µm)", colour = "Growth rate\n (µm/min)", title = paste(controlFromUnNested4$Treatment[1],Selected_root_ctrl, sep = " \\ "))+
  scale_x_continuous(breaks= Xaxis, labels = Xaxis, minor_breaks = NULL, limits = c(-5,8), expand = expansion(mult = c(0, .02)))+
  scale_y_continuous(breaks= Yaxis, labels = Yaxis, minor_breaks = NULL, limits = c(0,400), expand = expansion(mult = c(0.02, .02)))+
  theme(
    text = element_text(family = "GraphFont", face="plain", size=20, color = "black"),
    axis.text = element_text(family = "GraphFont", face="plain", size=20, color = "black"),
    panel.background = element_rect(fill = "White", colour = "black"),
    panel.grid.major = element_blank(),
    panel.grid.minor = element_blank(),
    legend.key = element_blank())+
  geom_vline(xintercept=0, linetype="dotted", color = "black", size=0.5)

SingleRootControlPlot
ggsave(SingleRootControlPlot, device = "svg", filename = paste(Selected_root_ctrl,"_beg-end-filter_control_",Sys.Date(),".svg"))

# Do graph for treated condition
Xaxis <- seq(-4,12,by = 2)
Yaxis <- seq(0,400, by = 50)
Selected_root_treat <- "B2_R09"

SingleRootTreatmentPlot <- treatmentFromUnNested4 %>%
  filter(Root==Selected_root_treat)%>%
  ggplot()+
  geom_line(mapping = aes(x=Time.hours.relative, y = Hair_Length_.micron., colour = RH_growth_rate_micron_per_min_shift, group = Root_ROI ), lineend = "round", size=2)+
  scale_size_manual(values = c(2,4))+
  scale_colour_gradientn(colours = c("blue","cyan","green","yellow","orange","red"), limits = c(0,2.1))+
  labs( x = "Time (hours)", y = "Hair length (µm)", colour = "Growth rate\n (µm/min)", title = paste(treatmentFromUnNested4$Treatment[1],Selected_root_treat, sep = " \\ "))+
  scale_x_continuous(breaks= Xaxis, labels = Xaxis, minor_breaks = NULL, limits = c(-5,8), expand = expansion(mult = c(0, .02)))+
  scale_y_continuous(breaks= Yaxis, labels = Yaxis, minor_breaks = NULL, limits = c(0,400), expand = expansion(mult = c(0.02, .02)))+
  theme(
    text = element_text(family = "GraphFont", face="plain", size=20, color = "black"),
    axis.text = element_text(family = "GraphFont", face="plain", size=20, color = "black"),
    panel.background = element_rect(fill = "White", colour = "black"),
    panel.grid.major = element_blank(),
    panel.grid.minor = element_blank(),
    legend.key = element_blank())+
  geom_vline(xintercept=0, linetype="dotted", color = "black", size=0.5)

SingleRootTreatmentPlot
ggsave(SingleRootTreatmentPlot, device = "svg", filename = paste(Selected_root_treat,"_beg-end-filter_treatment_",Sys.Date(),".svg"))

#calculate average growth rate for each hair in the linear growth area
controlFromUnNested4 <- filter(UnNested4LinearGrowth,UnNested4LinearGrowth$Treatment == "1/2MS")
treatmentFromUnNested4 <- filter(UnNested4LinearGrowth, UnNested4LinearGrowth$Treatment=="1/2MS_sucrose")

controlFromUnNested4BeforeT <- filter(controlFromUnNested4,controlFromUnNested4$Time.hours.relative <=0)
controlFromUnNested4AfterT <- filter(controlFromUnNested4,controlFromUnNested4$Time.hours.relative >0)
treatmentFromUnNested4BeforeT <- filter(treatmentFromUnNested4,treatmentFromUnNested4$Time.hours.relative <=0)
treatmentFromUnNested4AfterT <- filter(treatmentFromUnNested4,treatmentFromUnNested4$Time.hours.relative >0)

NestedcontrolFromUnNested4BeforeT <- controlFromUnNested4BeforeT %>% group_by(controlFromUnNested4BeforeT$Root_ROI)%>% nest()
NestedcontrolFromUnNested4BeforeT <-mutate(NestedcontrolFromUnNested4BeforeT,'Growth_rate_average_in_linear_growth'=map(data, ~ mean(.x$RH_growth_rate_micron_per_min)))
colnames(NestedcontrolFromUnNested4BeforeT) <- c("Root_ROI","data","Growth_rate_average_in_linear_growth_Ctrl_BT")
NestedcontrolFromUnNested4BeforeT <- transmute(NestedcontrolFromUnNested4BeforeT, Growth_rate_average_in_linear_growth_Ctrl_BT = map_dbl(Growth_rate_average_in_linear_growth_Ctrl_BT, 1))
NestedcontrolFromUnNested4BeforeT <- select(NestedcontrolFromUnNested4BeforeT, "Root_ROI", "Growth_rate_average_in_linear_growth_Ctrl_BT")

NestedcontrolFromUnNested4AfterT <- controlFromUnNested4AfterT %>% group_by(controlFromUnNested4AfterT$Root_ROI)%>% nest()
NestedcontrolFromUnNested4AfterT <-mutate(NestedcontrolFromUnNested4AfterT,'Growth_rate_average_in_linear_growth'=map(data, ~ mean(.x$RH_growth_rate_micron_per_min)))
colnames(NestedcontrolFromUnNested4AfterT) <- c("Root_ROI","data","Growth_rate_average_in_linear_growth_Ctrl_AT")
NestedcontrolFromUnNested4AfterT <- transmute(NestedcontrolFromUnNested4AfterT, Growth_rate_average_in_linear_growth_Ctrl_AT = map_dbl(Growth_rate_average_in_linear_growth_Ctrl_AT, 1))
NestedcontrolFromUnNested4AfterT <- select(NestedcontrolFromUnNested4AfterT, "Root_ROI", "Growth_rate_average_in_linear_growth_Ctrl_AT")

NestedtreatmentFromUnNested4BeforeT <- treatmentFromUnNested4BeforeT %>% group_by(treatmentFromUnNested4BeforeT$Root_ROI)%>% nest()
NestedtreatmentFromUnNested4BeforeT <-mutate(NestedtreatmentFromUnNested4BeforeT,'Growth_rate_average_in_linear_growth'=map(data, ~ mean(.x$RH_growth_rate_micron_per_min)))
colnames(NestedtreatmentFromUnNested4BeforeT) <- c("Root_ROI","data","Growth_rate_average_in_linear_growth_Treat_BT")
NestedtreatmentFromUnNested4BeforeT <- transmute(NestedtreatmentFromUnNested4BeforeT, Growth_rate_average_in_linear_growth_Treat_BT = map_dbl(Growth_rate_average_in_linear_growth_Treat_BT, 1))
NestedtreatmentFromUnNested4BeforeT <- select(NestedtreatmentFromUnNested4BeforeT, "Root_ROI", "Growth_rate_average_in_linear_growth_Treat_BT")

NestedtreatmentFromUnNested4AfterT <- treatmentFromUnNested4AfterT %>% group_by(treatmentFromUnNested4AfterT$Root_ROI)%>% nest()
NestedtreatmentFromUnNested4AfterT <-mutate(NestedtreatmentFromUnNested4AfterT,'Growth_rate_average_in_linear_growth'=map(data, ~ mean(.x$RH_growth_rate_micron_per_min)))
colnames(NestedtreatmentFromUnNested4AfterT) <- c("Root_ROI","data","Growth_rate_average_in_linear_growth_Treat_AT")
NestedtreatmentFromUnNested4AfterT <- transmute(NestedtreatmentFromUnNested4AfterT, Growth_rate_average_in_linear_growth_Treat_AT = map_dbl(Growth_rate_average_in_linear_growth_Treat_AT, 1))
NestedtreatmentFromUnNested4AfterT <- select(NestedtreatmentFromUnNested4AfterT, "Root_ROI", "Growth_rate_average_in_linear_growth_Treat_AT")

averageCombine <- merge(NestedcontrolFromUnNested4BeforeT,NestedcontrolFromUnNested4AfterT, by = "Root_ROI", all.x = TRUE, all.y = TRUE)
averageCombine <- merge(averageCombine,NestedtreatmentFromUnNested4BeforeT, by = "Root_ROI", all.x = TRUE, all.y = TRUE)
averageCombine <- merge(averageCombine,NestedtreatmentFromUnNested4AfterT, by = "Root_ROI", all.x = TRUE, all.y = TRUE)
averageCombine <- pivot_longer(averageCombine, cols = 2:5, names_to ="Treatment",values_to = "Average_RH_growth_rate")

for (rowToChange in 1:nrow(averageCombine)) {
  if(averageCombine[rowToChange,2] == "Growth_rate_average_in_linear_growth_Ctrl_BT"){
    averageCombine[rowToChange,"group_number"] <- 1
  }
  if(averageCombine[rowToChange,2] == "Growth_rate_average_in_linear_growth_Ctrl_AT"){
    averageCombine[rowToChange,"group_number"] <- 2
  }
  if(averageCombine[rowToChange,2] == "Growth_rate_average_in_linear_growth_Treat_BT"){
    averageCombine[rowToChange,"group_number"] <- 3
  }
  if(averageCombine[rowToChange,2] == "Growth_rate_average_in_linear_growth_Treat_AT"){
    averageCombine[rowToChange,"group_number"] <- 4
  }
}

#statistical tests
library(FSA) #install.packages("FSA")
library(rcompanion)
statAverageCombineForStat <- pairwise.wilcox.test(x = averageCombine$Average_RH_growth_rate, g= averageCombine$group_number, p.adjust.method = "bonferroni", paired = FALSE)

  #data preparation for post test
AverageCombineForStat2 <- averageCombine
AverageCombineForStat2 <- na.omit(AverageCombineForStat2, AverageCombineForStat2$Average_RH_growth_rate)
AverageCombineForStat2 <- as.tibble(AverageCombineForStat2)
temp <- as.double(AverageCombineForStat2$Average_RH_growth_rate)
temp2 <- as.integer(AverageCombineForStat2$group_number)
  #change list to single value in average column
  for (RowToProcess in 1:nrow(AverageCombineForStat2)) {
    AverageCombineForStat2[[RowToProcess,'Average_RH_growth_rate_factor']] <- temp[[RowToProcess]]
    AverageCombineForStat2[[RowToProcess,'group_number_numeric']] <- temp2[[RowToProcess]]
  }
  #post test
DunnTest_AverageCombineForStat2 <- dunnTest(Average_RH_growth_rate_factor ~ group_number_numeric, data = AverageCombineForStat2, method="bh")
  #get statistical groups letters
PH_AverageCombineForStat2 <- cldList(comparison = DunnTest_AverageCombineForStat2$res$Comparison,
                       p.value = DunnTest_AverageCombineForStat2$res$P.adj,
                       threshold = 0.01,
                       remove.space = TRUE)
PH_AverageCombineForStat2_Letter <- PH_AverageCombineForStat2$Letter
  # to set the position of the letters in the graph
AverageCombineForStat2_MAX <- aggregate(data = AverageCombineForStat2, Average_RH_growth_rate_factor ~ group_number_numeric, max)
AverageCombineForStat2_MAX$"letter" <- PH_AverageCombineForStat2_Letter
colnames(AverageCombineForStat2_MAX) <- c("group","mean","letter")

# find median of each group
AverageCombineForSMed <- averageCombine
AverageCombineForSMed <- na.omit(AverageCombineForSMed, AverageCombineForSMed$Average_RH_growth_rate)
AverageCombineForSMedNested <- AverageCombineForSMed %>%
  group_by(AverageCombineForSMed$group_number)%>%
  nest()%>%
  mutate('Med'=map(data, ~ median(.x$Average_RH_growth_rate)))
colnames(AverageCombineForSMedNested) <- c("Med_group_number","data","Med")
AverageCombineForSMedNested <- transmute(AverageCombineForSMedNested, Med = map_dbl(Med, 1))

AverageCombineForSMedNested$Med_group_number <- as.integer(AverageCombineForSMedNested$Med_group_number)

#Do violon plot
averageCombineplot <-
  ggplot()+
  geom_violin(data = averageCombine, mapping = aes(x=group_number, y = Average_RH_growth_rate, fill = Treatment), size=0.75)+
  labs(data = averageCombine, x = "", y = "Average root hair \n growth rate (µm min-1)")+
  scale_fill_manual(values=c('springgreen4','palegreen','magenta','plum1'))+
  geom_point(data = AverageCombineForSMedNested, mapping = aes(x=Med_group_number, y = Med)) +
  geom_text(data = AverageCombineForStat2_MAX, aes(x= group, y = mean+0.2, label = letter), size = 9)+
  theme(
    text = element_text(family = "GraphFont", face="plain", size=20, color = "black"),
    axis.text = element_text(family = "GraphFont", face="plain", size=20, color = "black"),
    panel.background = element_rect(fill = "White", colour = "black"),
    panel.grid.major = element_blank(),
    panel.grid.minor = element_blank(),
    axis.text.x=element_blank(),
    legend.position = "none")
averageCombineplot
  
ggsave(averageCombineplot, device = "svg", filename = paste("beg-end-filter_mean-RH-growth-rate_",Sys.Date(),".svg"))

#get samples sizes
AverageCombineForStat2_Pop_RH_Ctrl_BT <- AverageCombineForStat2 %>%
  filter(group_number=="1")%>%
  nrow()
AverageCombineForStat2_Pop_RH_Ctrl_AT <- AverageCombineForStat2 %>%
  filter(group_number=="2")%>%
  nrow()
AverageCombineForStat2_Pop_RH_Tret_BT <- AverageCombineForStat2 %>%
  filter(group_number=="3")%>%
  nrow()
AverageCombineForStat2_Pop_RH_Tret_AT <- AverageCombineForStat2 %>%
  filter(group_number=="4")%>%
  nrow()
```
